# Neural dynamics of emergent social roles in collective foraging by mice

**DOI:** 10.1101/2025.04.24.650547

**Authors:** Jaehyun Lee, Gyu-Hwan Lee, Seo Young Kim, DaYoung Jung, Jaehoon Kim, Jee Hyun Choi

## Abstract

We investigated how neural dynamics support role differentiation in group foraging by mice cohabiting with a predator-like spider robot serving as a mobile food platform. While solitary foraging behaviors were consistent among mice, group foraging revealed distinct roles–working, participating, and freeriding–that solidified over time, leading to an imbalance (Gini = 0.5) across five groups. Recordings from the medial prefrontal cortex (mPFC), nucleus accumbens (NAc), and basolateral amygdala (BLA) revealed enhanced beta power in active foragers. Workers exhibited a stronger fast beta drive from mPFC to BLA compared to participants, and role prediction accuracy based on spectral features improved over trials. Our findings highlight how neural dynamics underlie the emergence of distinct roles in collective action by mice.

**One-Sentence Summary:** Distinct foraging roles emerge in group-housed mice, driven by fast beta (24–32 Hz) activity in the mPFC-NAc-BLA circuit.

## Main text

In nature, group survival often relies on collective action to secure resources such as food and safety. However, when some individuals benefit without participation, conflicts arise between self-interest and group welfare, threatening the group through phenomena such as the freerider problem^1,2^ and the tragedy of the commons^3^. Societies nonetheless persist owing in large part to individuals who voluntarily participate^4^. This seemingly paradoxical behavior is often attributed to mechanisms such as genetic altruism^5^ or mutual altruism^6^ or altruistic punishment theories^7^. However, these theories do not address the real-time cognitive processes driving such participation. Alternately, in game-theoretical approaches, participation is guided by the individual rationality condition (IRC)^8^, which posits that individuals engage when perceived rewards outweigh costs. Mancur Olson expanded on this, noting that individual perceptions of these conditions vary, leading to diverse behaviors in collective actions^2^. However, the dynamics of voluntary participation under the IRC and its neural basis remain poorly understood.

To explore the individuality during collective action, we devised a group foraging task for mice, recording brain activity in the medial prefrontal cortex (mPFC), nucleus accumbens (NAc), and basolateral amygdala (BLA), which are linked to executive function, reward processing, and fear, respectively. The experiment takes place in a colosseum-like cage, where a group of mice cohabits with a predator-like robot restricted to the inner zone (Movie S1). The task starts by placing a snack on the robot; a mouse must venture into the robot zone, climb the robot to retrieve the snack, and then bring it out to a safe zone devoid of the robot, where the snack is shared with the other mice. The snacks provide about 40% of their daily calories per 10 trials. Our investigation focused on how individual roles within this collective foraging behavior – specifically, the propensity for food retrieval – evolve over time.

## Behavioral divergence in group foraging

Five groups with different member of mice compositions conducted the task, spanning five to fourteen consecutive days, completing between 50 to 162 trials of the group foraging task (**Supplementary Table 1, Supplementary Fig. 1**). In particular, Group A underwent extensive experiments, including pre– and post-foraging tube tests and individual training, to ensure all members achieved a comparable level of foraging proficiency (**Supplementary Fig. 2**, acting latency, *ps* > 0.716; retrieval time, *ps* > 0.163), making this group the primary focus of our analysis.

In the solitary task, all mice retrieved food (**Movie S2**); however, in group settings, behaviors diverged into working (retrieving food), participating (entering the robot zone without retrieving), and freeriding (staying outside the zone), reflecting distinct roles observed across tasks where any food brought out was shared among all (**Fig. 1a**). Over the course of two weeks, the foraging effort was concentrated on a single mouse (**Fig. 1b**) and the work and freeriding tendencies increased in the top worker and freeriding mice, respectively (**Fig. 1c**). In all tested groups, both participating and freeriding mice were observed, with freeriding mice consistently present (**Fig. 1d**). Regarding work ranks, as determined by worker trial percentages, the top workers averaged 72.2 ± 8.9% of trials, compared to 15.9 ± 4.1% for the secondary workers across five groups (**Fig. 1e**). A hidden Markov model based on role sequence data revealed distinct transitions across trials (**Fig. 1f-h**). Top workers were 4.5 times more likely to transition to roles involving more work (P_more_) than less work (P_less_) (*p* = 0.009). In contrast, participants and freeriders were more likely to transition to less work, with P_more_ being 0.7 and 0.3 times weaker than P_less_, respectively (*p* = 0.503 and p < 0.001).

**Figure 1.**
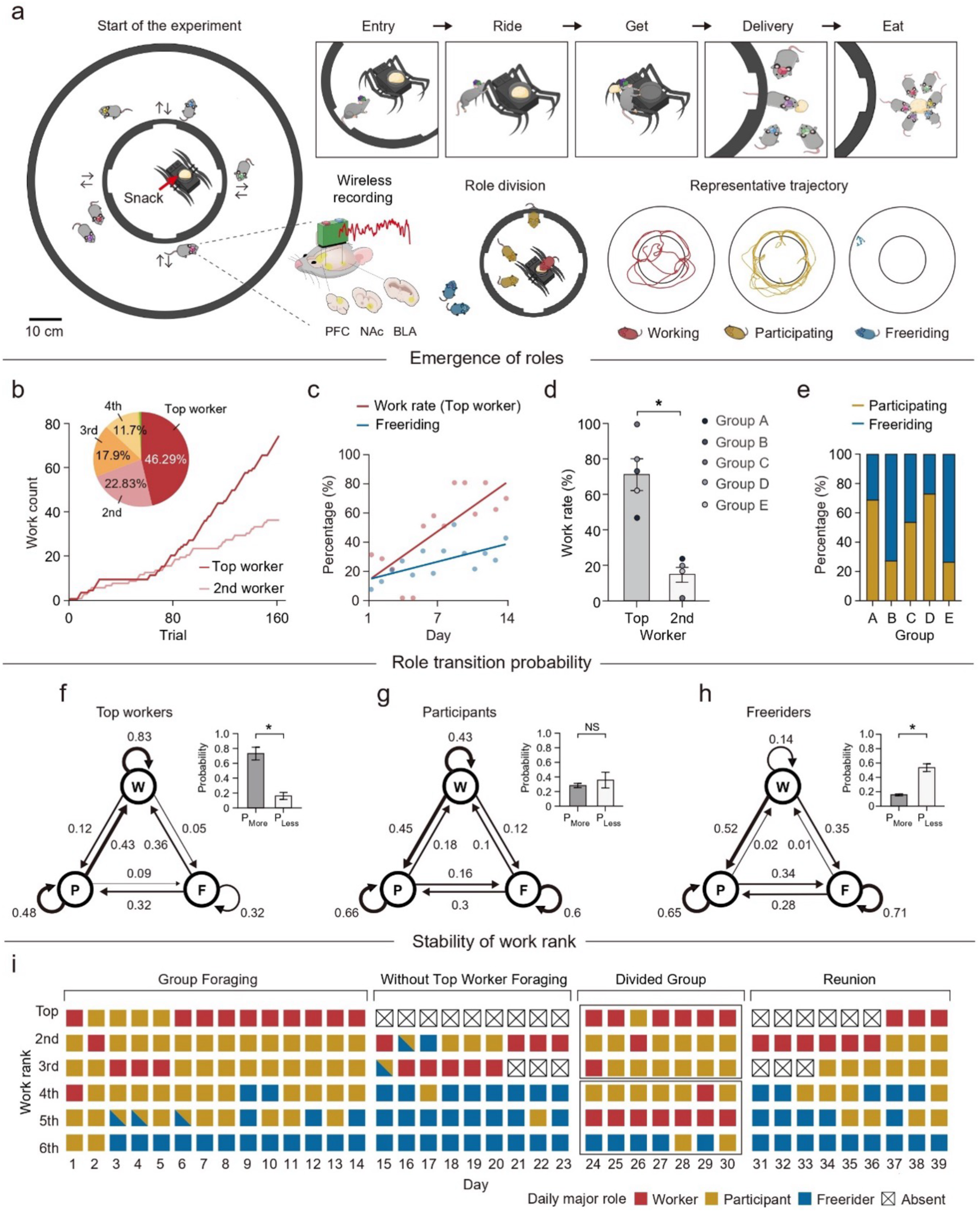
Emergence of roles and work rate consolidation. **a**, Group foraging paradigm and role differentiation. Mice cohabit with a predator-like spider robot in a colosseum-like cage (left). Five key events (Entry, Ride, Get, Delivery, Eat) define a foraging trial (top right), and each role—working, participating, or freeriding—exhibits distinct movement patterns (bottom right). **b**, Cumulative work contributions across 162 trials: the top worker steadily increases workload, while the second worker contributes less. The inset pie chart shows the proportional work distribution. **c**, The top worker’s work rate (slope = 5.09, R² = 0.587, *p* = 0.001) and individual freeriding rates (slope = 1.84, R² = 0.41, *p* = 0.013) both rose over time (daily averages shown). **d**, Participating (P) and freeriding (F) rates across five groups (A–E). Each rate is the ratio of total trials, excluding working trials. **e**, Top workers have a higher work rate than second workers (U = 25, *p* = 0.007; Mann– Whitney U test). Each dot represents a group value. **f–h**, Role transition probabilities (thicker arrows indicate higher probabilities) for top workers (**f**), participants (**g**), and freeriders (**h**). The inset bar graphs show the probability of transitioning to “more” or “less” work: **f**, t(4) = 4.681, *p* = 0.009; **g**, t(4) = 0.733, *p* = 0.503; **h**, t(25) = 6.49, *p* < 0.001; paired t-tests. **i**, Daily role assignments across experimental phases. Each day’s worker is the mouse with the highest work frequency; participants and freeriders are assigned based on their predominant non-work behavior. Diagonally divided squares indicate ties. Mice are ranked by their cumulative work rates. See Supplementary Table 1 for sample sizes. Error bars indicate mean ± s.e.m.; *p* < 0.05 is considered significant; NS, not significant.

Next, we tested the stability of work ranks by manipulating member composition (**Fig. 1i**). In group A, during “Without Top Worker Foraging” test (days 15–23) and after dividing into subgroups (days 24–30), the mice with the highest work rate within the subgroup worked the most, positioning itself as top worker. In “Reunion” test (days 31– 39), mice were reintroduced one by one in descending order of work rate until the group was fully reunited. In this process, the mouse with the lowest performance in the high work rate subgroup became the top worker in the low work rate subgroup, continued leading after the secondary worker was introduced, but stopped working when the original top worker returned. Other groups maintained their rank order within subgroups as well (**Extended Data 1**). These findings indicate that work roles and ranks are stable despite changed group compositions, with the highest work rate mice reliably assuming top worker roles.

## Traits of top workers

In human societies, voluntary action in group task has been observed to be either contingent upon individual abilities^9^ or conditional to the contributions of others^10^. To examine whether work rate correlates with individual capabilities, we assessed dominance rank, anxiety for height and open space, motor coordination, and working memory (**Supplementary Table 2**). None of these traits showed a statistically significant correlation with work rate (**Supplementary Fig. 3**). Dominance rank, assessed before and after the foraging experiments, remained stable (**Fig. 2a**). Across the five groups, top workers were primarily distributed within the middle ranks, whereas freeriders showed a polarized distribution, occupying either dominant or submissive positions (**Fig. 2b**). Top workers consistently demonstrated average motor coordination and height anxiety levels (**Fig. 2c**). In contrast, freeriders exhibited distinct patterns based on their dominance rank: dominant freeriders showed no clear trends, while submissive freeriders had reduced motor abilities and elevated height anxiety (**Fig. 2d**).

**Figure 2.**
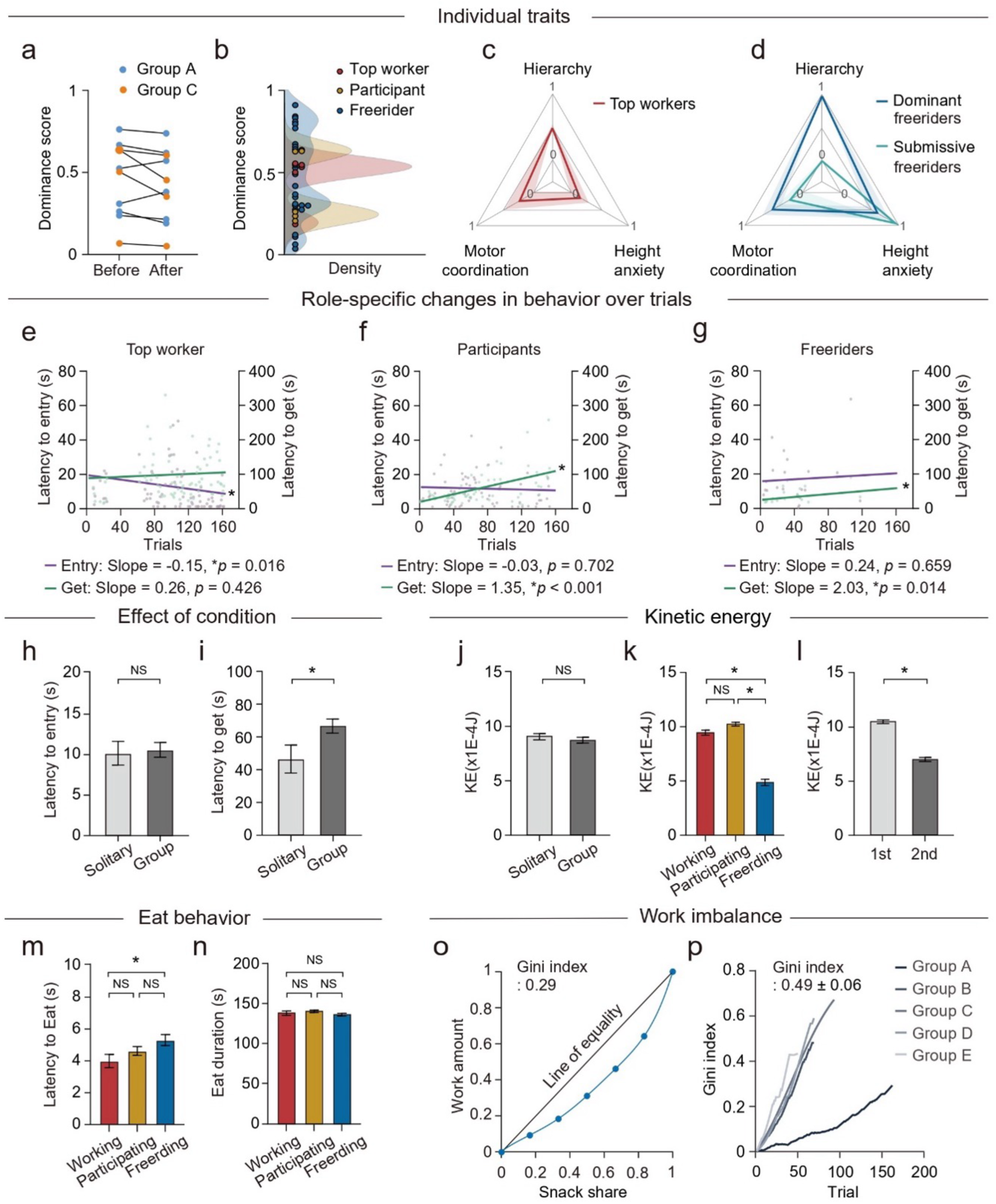
Role-dependent behavioral characteristics and work imbalance in group foraging. **a**, Normalized dominance scores before and after group foraging showed no significant changes in Groups A (six mice, W = 3, *p* = 0.843) or C (four mice, W = 10, *p* = 0.125; Wilcoxon matched-pairs signed-rank test). Higher scores indicate stronger dominance. **b**, Distribution of dominance for top workers, participants, and freeriders. Each dot represents one mouse; shaded regions show kernel density estimates. **c**, Top workers (three mice) exhibited moderate dominance, motor coordination, and height anxiety. **d**, Dominant freeriders (five mice) also showed moderate anxiety and motor coordination, whereas submissive freeriders (two mice) had lower motor ability and higher anxiety. **e**, As trials progressed, top workers (75 trials) showed decreased latency to entry (slope = – 0.15, R² = 0.076, *p* = 0.016) but no change in latency to get (slope = 0.26, R² = 0.008, *p* = 0.426). **f**, Participants (66 trials) showed stable entry latencies (slope = –0.03, R² = 0.002, *p* = 0.702) but increased latency to get (slope = 1.35, R² = 0.261, *p* < 0.001). **g**, Freeriders (21 trials) also showed no significant change in entry latency (slope = 0.24, R² = 0.01, *p* = 0.659) but increased latency to get (slope = 2.03, R² = 0.274, *p* = 0.014). Dots represent individual trials. **h**, Latency to entry did not differ between solitary (108 trials) and group (162 trials) conditions (U = 7561, *p* = 0.926). **i**, Latency to get was significantly longer in the group condition (U = 4423, *p* < 0.001). **j**, Mean instantaneous kinetic energy (KE) was similar in solitary vs. group conditions (U = 8488, *p* = 0.679). **k**, Working (W, n = 162) and participating (P, n = 554) mice showed higher KE than freeriding (F, n = 254) mice (*χ*²(2,972) = 237.5693, *p* < 0.001; W vs. P, *p* = 0.137; W vs. F, *p* < 0.001; P vs. F, *p* < 0.001). **l**, KE significantly decreased from Week 1 (87 trials) to Week 2 (75 trials; U = 885, *p* < 0.001). **m–n**, Latency to eat (**m**) and eating duration (**n**) by role (working, n = 159; participating, n = 539; freeriding, n = 251 for latency and n = 256 for duration). Freeriders took longer to begin eating (χ²(2,949) = 43.12, *p* < 0.001; W vs. P, *p* = 0.785; W vs. F, *p* < 0.001; P vs. F, *p* < 0.001), but total eating time did not differ significantly among roles (χ²(2,954) = 8.096, *p* = 0.017; W vs. P, *p* = 0.052; W vs. F, *p* = 0.904; P vs. F, *p* = 0.071). **o**, Lorenz curve illustrating work imbalance. The diagonal denotes perfect equity in normalized cumulative work and snack share; a 10^th^-degree polynomial fit (blue) indicates a Gini coefficient of 0.29 for Group A. **p**, The Gini coefficient rose over trials in all groups, averaging 0.49 ± 0.06 across trials. Error bars represent mean ± STD. unless otherwise stated. Mann–Whitney U tests were used for two-sample comparisons, and post hoc Dunn’s tests followed Kruskal–Wallis tests. \**p* < 0.05; NS, not significant.

## The presence of others changed the working aptitude

To investigate whether the presence of others influences foraging behavior, we analyzed the latency to enter the robot zone after snack placement (*1*_entry_) and the working time to retrieve the snack (*1*_get_) (Supplementary Table 3). Over repeated trials, the top worker showed a decreasing *1*_entry,_ while its *1*_get_ remained consistent (**Fig. 2e**). In contrast, both participants and freeriders maintained steady *1*_entry_ but exhibited longer *1*_get_ in trials where they acted as workers (**Fig. 2f-g**). Notably, these patterns observed during group trials were absent during solitary foraging (**Supplementary Fig. 4**). Compared to solitary condition, no mouse achieved *1*_entry_ or *1*_get_ faster than the solitary lower bound. *1*_entry_ was slower in 2.6–42.1% of trials but showed no significant difference on average (**Fig. 2h**). In contrast, *1*_get_ exceeded the solitary upper bound in 10.5–31.5% of trials and showed a significant overall increase (**Fig. 2i**).

We further examined the physical activity by calculating the kinetic energy (KE) from instantaneous speed. KE shows no significant difference between solitary and group settings (**Fig. 2j**). Although working and participating mice exhibited similarly high KE levels (**Fig. 2k**), their spatial behaviors diverged: working mice spent more time near the robot, while participating mice favored areas near the gates or outside the robot zone (**Extended Data 2**). Notably, physical activity decreased across all roles in the later trials compared to the earlier trials (**Fig. 2l**).

## Equal sharing and increase of imbalance between contribution and consumption

In this task, the common good is the snack. To estimate the snack consumption, we identified an eating cluster based on the behavior of mice gathering to eat, and calculated the latency to enter, *τ_eat_* and the total time spent within the cluster, *T_eat_* for each mouse (**Supplementary Table 4**). All mice began eating within 4.74 ± 5.79 s of the worker retrieving the snack out of the robot zone, with no significant individual differences (*p* = 0.0976). Excluding two mice, eating latency increased slightly over trials but remained negligible. Mice ate for 140.67 ± 31.72 s on average, with no significant individual differences (*p* = 0.3733). Eating duration increased by approximately 5 s per day, suggesting reduced urgency over time despite constant snack size. By roles, both *τ_eat_* and *T_eat_* did not show any significant difference (*p* = 0.630 and *p* = 0.2419, respectively, **Fig. 2m-n**), but a post-hoc test returned significant difference between working and freeriding mice in (*p* = 0.0315, **Fig. 2m**).

The surplus work, defined by the difference between cumulative normalized work amount and cumulative normalized snack share, highlighted a notable disparity across the roles. Furthermore, the Lorenz curve^11^, representing relative accumulation of normalized work amounts and normalized snack shared across animals, deviated significantly from the ‘line of equality’ (**Fig. 2o, Supplementary Fig. 5**). The Gini index (GI) for this group was 0.30, with an average of 0.50 ± 0.15 across five groups. The GI increased as trials progressed (**Fig. 2p**), indicating that the division of shares among mice significantly diverged from an equitable distribution based on contribution.

## Differential mPFC-NAc-BLA network dynamics by engaged roles

Mice inherently avoid the spider robot^12^ yet boldly enter its zone to retrieve snack placed on it, demonstrating distinct work behavior. This foraging task is similar to the approach-based foraging task described in study^13^, where food is obtained by walking toward a robot, but differs in requiring individuals to climb onto a moving robot to retrieve food. This process demands advanced cognitive abilities, including risk assessment, dynamic interaction, spatial awareness, and complex motor execution. To investigate the cognitive process underlying these behaviors, we analyzed mPFC-NAc-BLA network dynamics across various frequencies (**Supplementary Table 5**). During solitary foraging, distinct neural oscillations consistently associated with foraging behavior were observed across all mice: In the pre-Act period, high beta (*β*_high_, 24–32 Hz) activity was prominent in the mPFC. During the Act period, low beta (*β*_low_, 18–24 Hz) and *β*_high_ activity were observed across all regions, along with low gamma (*γ*_low_, 35–50 Hz) in the BLA (**Extended Data 3**). During the Eat period, low theta (*θ*_low_, 4–8 Hz) activity was consistently present across all regions.

Next, we analyzed the LFPs based on the roles shown during each trial. The ensemble-averaged spectrograms revealed distinct neural oscillation patterns associated with each role (**Fig. 3a**), particularly in the Act period (**Fig. 3b-d**). While working and participating mice exhibited similar spectral patterns, several differences were observed (**Extended Data 4a-l**, **Supplementary Table 6**). Compared to participating mice, working mice exhibited reduced high gamma (*γ*_high_, 70–90 Hz) in the mPFC during the pre-Act period. During the Act period, working mice showed decreased high theta (*θ*_high_, 8–12 Hz) and increased *β*_low_ in the BLA, elevated *β*_high_ across all regions, and reduced *γ*_high_ across all regions. Additionally, working mice displayed increased mPFC-NAc and mPFC-BLA coherence across all frequency bands except *γ*_high_ during both the pre-Act and Act periods, compared to participating mice (**Supplementary Table 7**).

**Figure 3.**
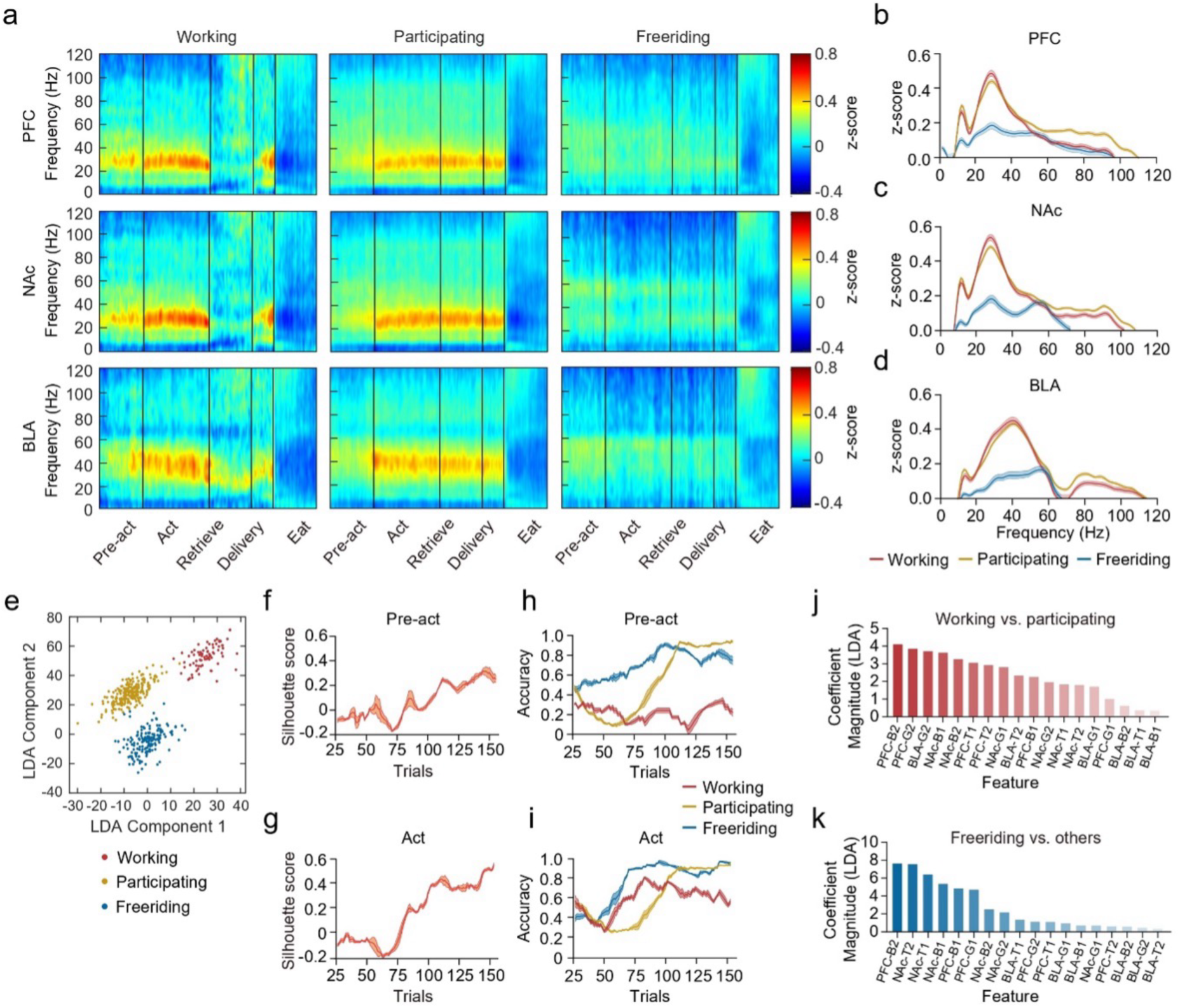
Neural activity across different brain regions during foraging task phases. **a**, Time-normalized, z-scored power spectrograms from the pre-Act through Eat periods in the mPFC, NAc, and BLA for working, participating, and freeriding mice. **b–d**, Z-scored spectral power distributions in the mPFC (**b**), NAc (**c**), and BLA (**d**) during the Act period for each role. Lines indicate means; shaded areas show s.e.m. See Supplementary Table 6 for statistical details. **e**, Projection of spectral patterns for individual trials onto the first two axes of a linear discriminant analysis (LDA), color-coded by role. **f–g**, Mean final silhouette scores over the last 20 trials in the pre-Act (**f**, 0.273 ± 0.005) and Act (**g**, 0.457 ± 0.015) periods. **h–i**, Trends in role prediction accuracy over trials. In the pre-Act period (**h**), mean accuracy over the last 20 trials is 0.292 ± 0.008 for working, 0.913 ± 0.003 for participating, and 0.79 ± 0.006 for freeriding. In the Act period (**i**), these accuracies rise to 0.607 ± 0.009, 0.925 ± 0.003, and 0.928 ± 0.009, respectively. **j–k**, LDA feature importance for distinguishing workers from participants (**j**) and freeriders from others (**k**). Bars represent coefficient magnitude, with higher values indicating greater discriminative power.

Compared to acting mice, freeriding mice exhibited no notable increases in neural activity, except for elevated *θ*_high_ and *γ*_high_ during the Eat period. Trajectory analysis revealed two distinct subtypes of freeriders: disengaged freeriders, who remained in the nest even when the snack was placed on the robot’s head, and engaged freeriders, who actively moved around the robot zone without entering it. Over time, the proportion of disengaged freeriders increased (**Extended Data 5**). During the pre-Act period, disengaged freeriders showed significantly higher *γ*_high_ in the mPFC compared to engaged freeriders. In contrast, engaged freeriders exhibited increased *β*_high_ in the BLA and *γ*_high_ in all regions during the Eat period compared to disengaged freeriders (**Supplementary Table 8**).

## Neural signatures of role consolidation in collective foraging

To determine whether neural dynamics underpin the progressive stabilization of roles across repeated trials, we applied a linear discriminant analysis (LDA)-based classification over a two-week period. The LDA model achieved a classification accuracy of 0.923, demonstrating robust separability of behavioral roles (**Fig. 3e**). Mahalanobis distances between roles were 1.6 ± 0.74 (workers vs. participants), 3.33 ± 1.43 (workers vs. freeriders), and 1.95 ± 1.21 (freeriders vs. participants), underscoring the distinct neural dynamics associated with each role. Clustering performance measured by silhouette score improved progressively, with gradual gains in clustering using pre-Act phase data (**Fig. 3f**) and a pronounced increase in the Act phase by the second week reflected in a rise of silhouette score (**Fig. 3g**).

Temporal development of role-specific predictability revealed distinct patterns. During the pre-Act period, predictability of working mice remained low, whereas freeriding mice exhibited a steady increase during early trials. Participating mice demonstrated a sharp rise in predictability after the first week (**Fig. 3h**). In the Act phase, both working and freeriding mice exhibited a marked increase in predictability around the fifth day, preceding a subsequent rise in participating mice predictability (**Fig. 3i**). These shifts suggest a dynamic adaptation of neural representation over time, with inflection point across roles showing distinctive alignments for different behavioral epochs (**Supplementary Fig. 6**).

Feature importance analysis of the LDA model identified *β*_high_ power in the mPFC as the primary discriminator between working and participating mice, as well as freeriding and other mice (**Fig. 3j-k**). Next, *γ*_high_ powers in the mPFC and BLA, along with *β*_low_ and *β*_high_ in the NAc, were critical for distinguishing working and participating roles (**Fig. 3j**). Slow oscillations (*θ*_low_, *θ*_high_, and *β*_low_) in the NAc were important for identifying freeriders (**Fig. 3k**).

## Critical role of prefrontal beta drive in determining working behavior

Next, we focused on critical moments, such as entering the robot zone or retrieving the snack and observed prominent *β*_high_ activity across all regions (**Extended data 6a-g**). Fast oscillations, including *β*_high_, primarily appeared as transient bursts rather than sustained oscillations^14^. Based on this, we detected transient oscillations in the *β*_high_ and *γ*_high_ bands^28^ (**Fig. 4a**), and revealed that burst occurrence density had notable role-specific differences (Extended Data 6h-s). Correlation analysis with working and freeriding rates revealed that *β*_high_ burst occurrence rates positively correlated with work rate across all regions (**Fig. 4b-d**) and negatively correlated with freeriding rate in the mPFC and NAc, but not in BLA (**Fig. 4e-g**). *γ*_high_ exhibited similar patterns in mPFC and NAc but was negatively correlated with work rate in the BLA (**Supplementary Fig. 7**). The role differentiation observed in *β*_high_ and burst occurrence densities (**Fig. 4h-i**) are generally align with spectral results (**Extended data 4e-p**).

**Figure 4.**
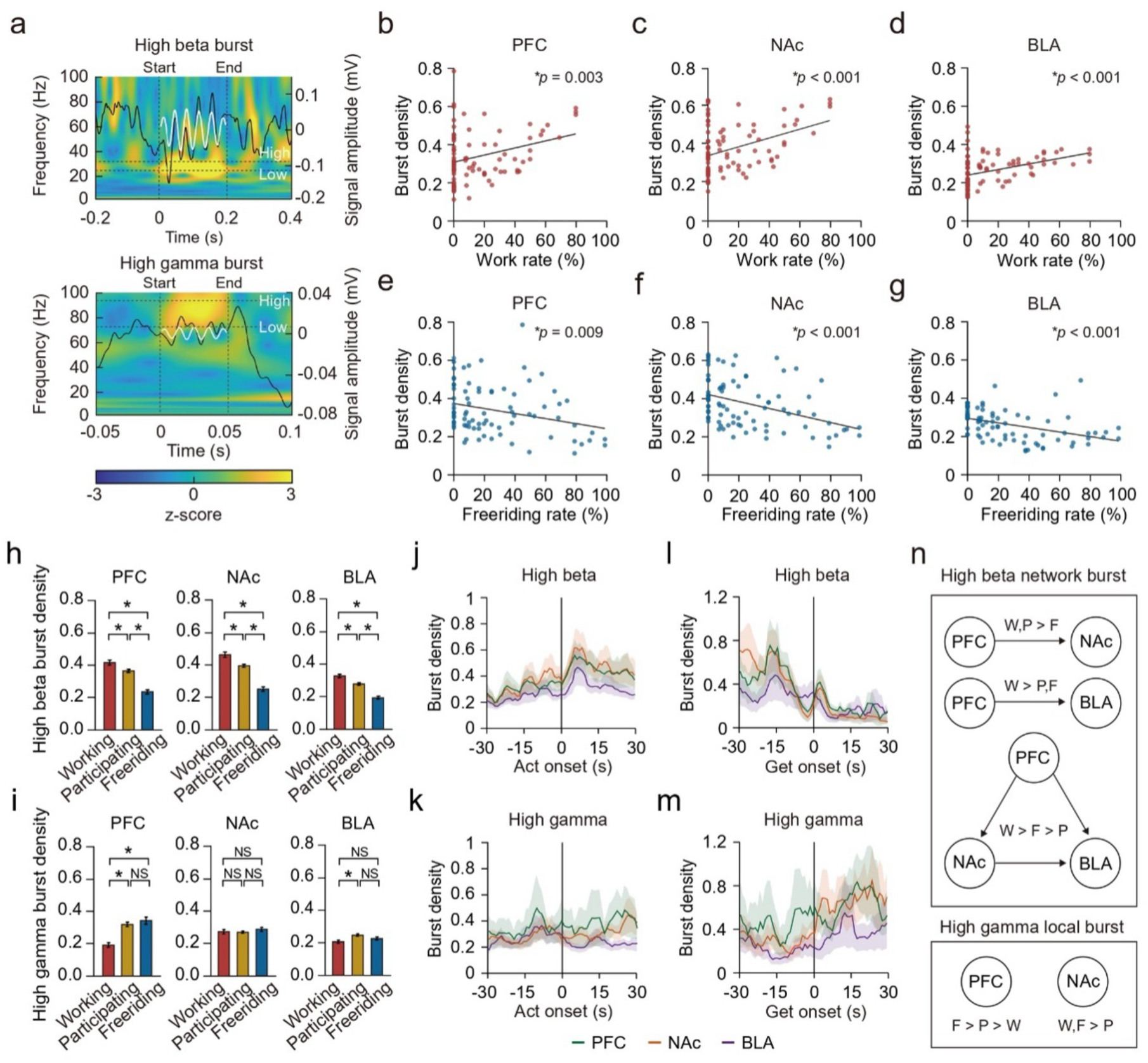
Beta and gamma burst dynamics in relation to behavior and network connectivity. **a**, Representative time-frequency plots of a high-beta burst (top) and a high-gamma burst (bottom). Black lines show LFP traces with burst amplitude (right *y*-axis), and white lines highlight band-pass-filtered burst events. **b–g**, High-beta burst density correlates positively with work rate in the mPFC (**b**, slope = 0.0018, R² = 0.097, *p* = 0.003), NAc (**c**, slope = 0.0023, R² = 0.158, *p* < 0.001), and BLA (**d**, slope = 0.0014, R² = 0.16, *p* < 0.001). High-gamma burst density correlates negatively with freeriding rate in the mPFC (**e**, slope = –0.0013, R² = 0.079, *p* = 0.009), NAc (**f**, slope = – 0.0018, R² = 0.162, *p* < 0.001), and BLA (**g**, slope = –0.0011, R² = 0.176, *p* < 0.001; linear regression analysis). Each dot represents daily averages for one mouse (n = 84). **j**, Comparison of high beta burst density among working, participating, and freeriding mice across the brain regions during the Act period: PFC (left, χ²(2,924) = 99.9987, p < 0.001; W vs. P, p = 0.0023; W vs. F, p < 0.001; P vs. F, p < 0.001), NAc (middle, χ²(2,924) = 117.2791, p < 0.001; W vs. P, p < 0.001; W vs. F, p < 0.001; P vs. F, p < 0.001), and BLA (right, χ²(2,924) = 106.5605, p < 0.001; W vs. P, p < 0.001; W vs. F, p < 0.001; P vs. F, p < 0.001). **k**, Comparison of high gamma burst density among working, participating, and freeriding mice across the brain regions the Act period: PFC (left, χ²(2,924) = 35.957, *p* < 0.001; W vs. P, *p* < 0.001; W vs. F, *p* < 0.001; P vs. F, *p* = 0.843), NAc (middle, χ²(2,924) = 2.0569, *p* = 0.357; W vs. P, *p* = 0.879; W vs. F, *p* = 0.385; P vs. F, *p* = 0.476), and BLA (right, χ²(2,927) = 12.7473, *p* = 0.0017; W vs. P, *p* = 0.001; W vs. F, *p* = 0.178; P vs. F, *p* = 0.188). **l**, High-beta burst density shows a sharp drop just before the Get moment and rebounds afterward in the mPFC and NAc (PFC: W = 0, *p* < 0.001; NAc: W = 0, *p* < 0.001; BLA: W = 127.5, *p* = 0.974). **m**, High-gamma power remains unchanged following the Get moment (PFC: W = 99.5, P = 0.837; NAc: W = 121, *p* = 0.166; BLA: W = 127.5, *p* = 0.19; Wilcoxon matched-pairs signed-rank tests). **n**, High-beta bursts (top) show stronger directed interactions from mPFC to BLA in workers (W > P, F) and from mPFC to NAc in both workers and participants (W, P > F). High-gamma bursts (bottom) exhibit more localized patterns, with elevated activity in the mPFC for freeriders (F > P > W) and in the NAc for workers and freeriders (W, F > P). Arrows denote significant directed interactions (see **Methods**).

Examining these dynamics in finer time scales, *β*_high_ burst density in both working and participating mice peaked after the Act onset across the regions (**Fig. 4j**), while *γ*_high_ burst occurrence decrease following the Act moment in the NAc (**Fig. 4k**). Right after getting the snack, *β*_high_ bursts in working mice decreased prior to retrieval but increased briefly but strongly right after getting the snack (**Fig. 4l**), accompanied by a quick increase and persistently elevated *γ*_high_ bursts occurrence rate (**Fig. 4m**).

We further identified networks of concurrent bursts across regions and evaluated the significance of their directionality using the directed phase lag index (dPLI). *β*_high_ bursts showed directionality towards the NAc and BLA from the mPFC, and from the NAc to the BLA. Notably, *β*_high_ bursts from the mPFC to the BLA showed pronounced increase in workers than in participants. In contrast, *γ*_high_ bursts showed locally confined pattern without a clear directionality, with a notable increase in the mPFC of freeriding mice. These results, encapsulated in Fig. 4n, emphasize the mPFC’s central role in modulating network dynamics during group task.

## Discussion

Our study provides novel insights into the real-time neural activities underlying voluntary participation in collective action, addressing a neurodynamic perspective on the variability of individual rationality conditions^2^. We found that distinct roles−workers, participants, and freeriders−emerge and stabilize over time, each exhibiting unique oscillatory signatures within the cost-benefit control system (mPFC-NAc-BLA network). Specifically, workers and participants displayed elevated beta activities, with workers exhibiting a pronounced fast beta drive from the mPFC to the BLA, suggesting a stronger top-down regulatory mechanism for commitment to action. Importantly, our study is the first to reveal that the neurodynamic properties of this decision-making circuit directly underpin role differentiation, linking oscillatory activity patterns to long-term behavioral specialization in mouse group dynamics.

Role predictability based on neural spectral features improved over repeated trials, reflecting the progressive consolidation of behavioral roles through network-level plasticity. While previous research has explored cooperation and freeriding through social norms or decision-making models^14–16^, few studies have examined the real-time neural mechanisms governing these social structures. By demonstrating how the dynamic interplay between risk assessment (BLA), reward valuation (NAc), and executive control (mPFC) determines voluntary working, our finding establish a neurodynamic framework for understanding role specialization in collective action under natural conditions.

In our groups of mice, behavioral patterns do not align with previous explanations of work imbalance based on greater endowments^16^, social rank^17,18^, reciprocity^19^, or prosocial motivation^20^. Food consumption remained consistent across roles, and top workers consistently emerged from the middle rank, opting out of rewards associated with higher endowments and submissiveness. Furthermore, we found no evidence of role-switching indicative of reciprocity. In a supplementary experiment where worker was given the opportunity to eat alone, most mice chose to do so, challenging altruistic interpretations (**Supplementary Fig. 8**). Attributing to a failure in establishing social rules in mice contradicts previous observation of turn-taking^21^ and altruistic behaviors^20^. Alternatively, our findings align with Olson’s logic of collective action^2^, where work imbalance arise from individual differences in perceived costs and expected benefits. In this cost-benefit framework, mice will act when expected benefit outweighs the perceived cost, and the variations in cost-benefit assessments will drive the working skew.

The neurodynamic correlates of perceived costs are likely proxied by BLA gamma activity, which has been associated with vigilance^13^ and peaks upon encountering the spider robot^22^. In the solitary condition, all mice exhibited increased BLA gamma. However, under group conditions, working mice showed a significantly smaller increase compared to participating mice, suggesting reduced threat sensitivity or a greater sense of control over the foraging task. This distinction sets them apart from participants, who entered the robot zone but refrained from retrieval.

Conversely, the benefit of working was unlikely to be the snack itself, as all mice had equal access to it. If the snack was the only benefit, both freeriders and workers would have made rational choices, whereas workers would not. However, when expanding the concept of benefit to include the internal perceptual reward associated with working, we can infer that behaviors exhibiting reward-related neural patterns likely correspond to moment of reward receipt. So far, increased beta activity in the frontal^23,24^ and ventral striatal^25,26^ regions have been well-documented as a neural signature of reward salience. Additionally, enhanced gamma activity in the NAc has been reported in mice at foraging site^27^. Interestingly, in our data, neither beta nor gamma activities were observed during eating. In contrast, during Act period, beta activity was significantly higher in working mice, whereas gamma activity was lower compared to participating mice showing anti-phasic relationship of beta and gamma. On the other hands, immediately after retrieving the food, we observed a sharp but transient increase in beta burst occurrence, accompanied by a rapid yet sustained increase in gamma burst occurrence. Taken together, these findings suggest that, for worker mice, the act of obtaining the snack itself−rather than the snack alone−would serve as the primary motivational reward.

Furthermore, the difference in beta-gamma relationship suggests distinct functional roles of beta activity. Notably, the stronger mPFC-to-BLA beta directionality in worker compared to participants would indicate top-down inhibition of bottom-up gamma activity^28^, as evidenced previously by the suppressed yet temporally nested gamma oscillations observed during flight from the robot^29^, supporting decisive action execution. In contrast, the simultaneous increase in beta and gamma activity following food retrieval may reflect mPFC encoding of expected benefits throughout the course of action^14^. This finding underscores the multifaceted nature of beta oscillations, aligning with recent studies on simultaneous dopamine release and beta oscillations at multiple sites, which have demonstrated spatially and temporally heterogeneous beta activity in relation to dopamine release^30^.

Lastly, we question whether worker behavior is driven by reward-based reinforcement learning. Faster task initiation and reduced kinetic energy with repeated trials might suggest reinforcement learning. However, the unchanging work durations and the progressive increase in beta power, contrary to typical reinforcement learning patterns^25^, raise the possibility of goal-directed behavior driven by innate motivation. Given the critical role of prefrontal cortex in foraging-related decision-making^31,32^, circuit-level interventions will be essential to distinguish between reward-driven reinforcement and innate motivation. Furthermore, simultaneous tracking of neuromodulators (dopamine, norepinephrine) and neural oscillations could provide deeper insight into the mechanisms underlying spontaneous role-taking in mice.

This study demonstrates that mice can be used as a model of high-level social processes, revealing distinct roles in group foraging that go beyond standard single-subject decision-making paradigms. By connecting social science perspectives on collective action with real-time neural analysis, we show how individual cost-benefit assessments manifest in distinct mPFC-NAc-BLA activity patterns, advancing our understanding of complex group behaviors.

## Acknowledgments

We thank the members of the Computational Cognitive and Systems Neuroscience Laboratory in KIST for their helpful discussions. This work was supported by the Basic Science Research Programs (RS-2022-NR070539, 2021R1A6A3A01087727) and the Global Cooperative Convergence Research Program (RS-2024-00460958) by National Research Foundation of Korea and the intramural grant of the Korea Institute of Science and Technology (2E32901).

## Author contributions

JHC and JHK conceptualized the study. JL designed the experimental paradigm. SYK, JL, GHL, and DJ acquired the data. JL, GHL, and JHC conducted the investigation. JL, JHC, and GHL performed the analysis. JL and DJ visualized the results. JHC obtained funding and supervised the project. JHC wrote the original draft. JHC, JL, and GHL reviewed and edited the manuscript.

## Data and materials availability

The primary analysis scripts and partial datasets used for this study are available at https://github.com/jeelabKIST/GroupForaging/. Due to repository size constraints, only the first trial from each day’s LFP and positional data for Group A have been deposited. Additional datasets, including the full LFP recordings, positional data, and video files, are available upon reasonable request from the corresponding author.

## Supplementary Materials

Methods

Extended Data 1-6

Supplementary Figures S1 to S9

Tables S1 to S9

Movies S1 to S2

## Methods

### Methods and Materials

#### Animals

Groups A–C consisted of 17 transgenic Thy1-COP4/EYFP male mice (B6.Cg-Tg(Thy1-COP4/EYFP)18Gfng/J, Stock number 007612, Jackson Laboratory, ME, USA). Groups D and E included 19 wild-type C57BL/6J male mice (Stock number 000664, Jackson Laboratory, ME, USA). The numbers of mice used in each group is specified in Supplementary Table 1. In all experiments, naive mice from at least three litters, aged 8–10 weeks, were randomly selected and combined to form a group to avoid potential behavioral confounds due to kinship^1^. Each experimental group was group-housed and allowed to interact freely for at least a week before being habituated to the experiment environment. Mice that exhibited abnormal aggression, defined as causing visible injuries to cage mates during the first 3-day observation period, were removed and replaced with other mice. Mice were co-housed under a 12:12 h light/dark cycle throughout all experiments. Mice had *ad libitum* access to food and water, except during the group foraging experiment period. All experimental procedures were approved by the Korea Institute of Science and Technology Animal Care and Use Committee (permit number: KIST-IACUC-2023-032-8) and complied with the National Institute of Health Guidelines for minimizing the pain and discomfort of animals.

#### Surgery

The CBRAIN surgery procedure was conducted as previously described^2^. Briefly, to obtain *in vivo* electrophysiological data from the medial prefrontal cortex (mPFC), nucleus accumbens (NAc), and basolateral amygdala (BLA) in freely behaving mice, all subjects underwent surgery for the chronic implantation of tungsten electrodes (0.005-inch shaft diameter, Cat# 577100, A-M Systems, WA, USA). All surgeries were performed under ketamine/xylazine anesthesia (120 and 6 mg/kg, respectively) using a stereotaxic apparatus (Model 900, David Kopf Instruments, Tujunga, CA, USA). The target coordinates were as follows: AP +1.4 mm, ML ±0.3 mm, DV −2.3 mm from bregma for the mPFC; AP +1.2 mm, ML ±0.55 mm, DV −4.3 mm from bregma for the NAc; and AP −1.5 mm, ML ±0.55 mm, DV −4.3 mm from skull for the BLA. To ensure accurate placement of electrodes in the BLA, an intraoperative microstimulation was conducted using Intan system to evoke observable neural responses. The ground and reference electrodes were implanted into the cerebellar skull to ensure signal stability. All electrodes were secured using dental cement (Vertex Self-Curing, Vertex Dental, Zeist, The Netherlands).

### Enriched environment

#### Colosseum cage

All experiments were conducted in a round-shaped cage named the “Colosseum cage.” The cage was constructed using plexiglass and consisted of two concentric cylinders, one placed within the other. The diameters of the inner and outer cylinders were 34 cm and 79 cm, respectively, while their heights were 18 cm and 50 cm, respectively. To prevent light reflection, the inner cylinder was painted with black water-based acrylic paint (Musou Black, Koyo Orient Japan Co., Ltd., Japan), and the outer cylinder was covered with waterproof black matte wallpaper. The inner cylinder had four gates, each measuring 6 cm × 10 cm (width × height). A water-dispenser was securely attached to one side of the outer cylinder. An ample amount of pulp chips (DooYeol Biotech Inc., South Korea) was provided as bedding material, along with cotton balls (DooYeol Biotech Inc., South Korea) as nesting material in the outer area. Environmental enrichments, including treadwheels, small huts, slides, and stairs made of white plexiglass, were included to encourage exploratory behavior and simulate a naturalistic environment. The Colosseum cage was placed in a soundproof, electromagnetic field-free room. All experiments were monitored from a separate room to minimize disturbance from the presence of experimenters.

#### Spider robot

A lightweight plastic spider robot with dimensions of 14 cm × 11 cm × 7 cm (length × width × height) was used in this study (Model No. 18143 Wired R/C Spider Robot, Academy Plastic Model Co., Ltd., South Korea). The robot’s body was painted with black paint to minimize light reflection. To serve as a fish snack tray, a small concave black plate was firmly attached to the top of the robot. Fish snack (Hanjin Foods, South Korea) is about 4 cm in diameter and weighs 0.8 g (3.08 kcal each). The robot’s motion was controlled by an Arduino chip, programmed to execute four types of movement (forward, backward, left turn, and right turn) for a randomly set duration between 1 and 2 seconds with 0.5-second intervals between each action. The amplitude of motor noise ranged from 75 to 85 dB. Since the spider robot is only placed in the inner cylinder, it is referred to as the robot zone, while the outer cylinder is referred to as the residence zone. To maintain threat in the robot zone, a different version of spider robot with no tray on the top, programmed to move for 3 minutes every 20 minutes, was placed in the colosseum cage during non-experimental period. The Arduino codes for motion control are available at https://github.com/jeelabKIST/GroupForaging/.

### Behavioral tasks

#### Foraging tasks

Prior to foraging tasks, mice were co-housed in the Colosseum cage for at least one week to habituate and remained together until all experiments were completed. Mice had free access to food and water ad libitum. When experiments such as solitary tasks or training or modified tasks required handling individual mice, they were temporarily placed in a cube-shaped cage made of plexiglass, featuring an open ceiling and a width of 40 cm. After the habituation period, food restriction was implemented to reduce their body weight to 85–90% of the initial value to enhance their motivation for food retrieval. The foraging tasks commenced once all mice had reached the target body weights. During the foraging tasks, food was provided at 50–80% of their regular daily intake to maintain their reduced body weight at a stable level.

Each trial began with a fish snack being placed on the spider robot by experimenter. using a fishing rode with 60 cm line. Since the robot zone and the residence zone are separated, we used another snack attached to the end of a 60 cm fishing line to bring it close to the mice as a signal to start the experiment. Approximately 10 trials were conducted per day, with a 5-minute interval between trials. The foraging task involves voluntary actions performed by the mice, with the behavioral events marked by their actions. Typical behavioral sequence includes the onsets of **Start** (placing the snack on the robot), **Entry** (mice entering the spider zone), **Ride** (a mouse riding on the spider robot), **Retrieval** (a mouse retrieves the snack from the spider robot), and **Eating**. The task concluded 5 and 2 minutes after Eating onset in group and solitary conditions, respectively, after which any remaining snacks were removed. The group tasks lasted between 5 and 14 days. Depending on the group, individual training or varied foraging tasks were conducted, resulting in a total duration of 10 to 49 days for the entire foraging experiments. Individual training involved attracting the mice into the robot zone by dangling a snack from a fishing line. Once inside, the snack was placed on the robot, allowing them to experience retrieving the food.

#### Modified foraging tasks

To examine group dynamics under different member compositions, we conducted three modified foraging tasks. First, in the *Without Top Worker Test*, we investigated the behavior of the remaining mice after removing the individual with the highest working rate in previous trials. The foraging task was repeated until one of the remaining mice consistently exhibited stable top worker behavior. Once identified, this mouse was also removed, and the process was repeated until the group size was reduced to half of its original size. Next, we conducted the *Divided Groups Test*, in which the remaining mice from the *Without Top Worker Test* were split into two subgroups: one consisting of the mice that remained after the removal of Top workers and the other consisting of the mice previously identified as top workers. Each subgroup then underwent the group foraging task independently. Finally, once stable top workers emerged in each subgroup, we performed the Reunion Test by systematically reintegrating the previously removed mice into the half-sized group from the *Without Top Worker Test* in reverse order. The experiment continued until a new stable top worker emerged and was repeated until the original group size was fully restored.

Another modified foraging task, the *Fixed Snack Test*, involved placing a clip on the robot instead of using a fish rod to position the snack. The clip secured the snack in place, preventing the mice from easily removing it, so they could only eat it while staying on the robot.

#### Prosociality Test

To examine whether the act of bring out food reflected prosocial intentions, we introduced an “eating chamber” that allowed the mouse to make a choice to eat alone or share. The eating chamber is made with transparent plexiglass box measured 14 cm × 14 cm × 30 cm (length × width × height) and featured two one-way doors: one providing entry from the robot zone into the eating chamber and the other allowing exit towards the residence zone. Each door measured 10 cm × 10 cm (length × width). To provide olfactory cues regarding food consumption, several 0.5 cm holes were drilled into the chamber walls. In this test, the mouse was placed directly into the robot zone at the start, while the other three doors were kept closed to prevent exit through alternate routes. If the mouse failed to retrieve the snack or fled into the chamber, it was removed and reintroduced into the zone after at least 5 minutes. The tet continued until the snack was fully consumed, regardless of whether it was eaten inside or outside the chamber.

#### Tube test

To assess the impact of social hierarchy on individuality in collective action, we conducted the tube test ^3^. Briefly, two mice were placed in separate chambers (11 cm × 8.5 cm × 8.5 cm in length × width × height) connected to a transparent plexiglass tube (30 cm × 3 cm × 9.5 cm in length × width × height), but initially blocked by transparent doors. A second transparent door was positioned at the center of the tube. After habituation, the chamber doors were opened, allowing the mice to enter the tube. Once both mice reached the center and faced each other, the central door was also opened. A winner was determined as the mouse that successfully pushed its opponent back into the opposite chamber. Trials lasting over 1 minute were considered ties. Each pair was tested once per day for four to eight days.

#### Y-Maze test

To assess spatial working memory, we conducted the Y-maze test^4^. A mouse was placed at the center of a Y-shaped maze, where three identical arms (30 cm × 4.5 cm × 12 cm in length × width × height) extended at equal angles in a dimly lit environment (10 Lux). The sequence of arm entries was recorded, and the number of times the mouse entered a different arm from the one it previously visited was counted. The arm alternation rate (%) was then calculated by dividing the number of successive entries into three different arms by the total number of arm entries minus two, to account for the minimum entries required for alternation. The test was conducted

#### Open field test

To monitor the exploratory and anxiety-like behaviors of individual mice, we performed the open field test (OFT)^5^. A mouse was initially placed at the periphery of a white plexiglass box (40 cm x 40 cm) in a dimly lit environment (10 Lux). The mouse’s movements were recorded for 20 minutes, and its position was tracked using EthoVision XT 16.0 (Noldus Information Technology, Wageningen, The Netherlands). The times spent in the central and peripheral zones were measured as indicators of exploratory and anxiety-like behavior, respectively.

#### Elevated plus maze test

To evaluate anxiety-like behavior, we conducted the elevated plus maze (EPM) test^6^. A mouse was placed at the center of a cross-shaped maze, elevated 40 cm above the ground, with two open arms and two enclosed arms surrounded by 15 cm high walls. Each arm measured 60 cm × 8 cm (length × width). The test was conducted under the dim light (10 Lux). The mouse’s movements were recorded for 10 minutes, and its position was tracked using EthoVision XT 16.0. The time spent in the open arm was measured as an inverse indicator of height-related anxiety.

#### Rotarod test

To assess motor coordination and balance, we conducted the rotarod test^7^ using a commercial rotarod apparatus (Catalogue number: 99-0061, LAIYUE Biotech Co., Ltd., China). Each mouse was placed in a separate lane, starting at 4 rpm, with the speed gradually increased to 40 rpm over 5 minutes. The latency to fall was recorded as a measure of motor performance. Each mouse was tested four times a day for four consecutive days, with at least 15-minutes intervals between trials to prevent fatigue.

### Recording

#### LFP recording

We used the CBRAIN headstage, a telemetry for collective brain research^2^, featuring an embedded 16-channel differential amplifier (RHD2216, 16-bit, INTAN Technologies LLC, Los Angeles, CA, USA) for recording local field potentials (LFPs). Data were transmitted via a Bluetooth System-on-Chip (nRF52832, Nordic Semiconductor) at 8 Mbit/s after band-pass filtering (1 Hz−4 kHz). CBRAIN also features onboard computing for detecting transient oscillations, indicated in real time by an LED light on the headstage. LFP recording was conducted at a sampling rate of 1024 Hz, and the data were processed using CBRAIN Studio, a custom MATLAB-based software (MATLAB 2021a, Mathworks Inc., Natick, MA, USA). The LED was activated when the power of beta or gamma oscillations exceeded the baseline by 3 standard deviations, with the baseline defined as the average power recorded during the pre-experiment period. This real-time indicator provided insight into which neural oscillations were associated with foraging behavior.

#### Video recording

A spectral camera (Lt225, Teledyne Lumenera, Ottawa, ON, Canada) was mounted 1.5 m above the center of the cage, capturing a top-down view of the entire area. Videos were recorded at 30 frames per second with a 1000 × 1000 pixel resolution using using StreamPix 8 software (Norpix Inc., Montreal, Quebec, Canada). The camera settings were optimized to capture detailed behavioral movements without motion blur, while the resolution provided sufficient spatial detail for precise analysis. Lighting conditions (70 Lux) were controlled to provide uniform illumination, minimizing shadows that could affect video quality or behavior. The recorded videos were stored directly on a dedicated storage device for subsequent analyses.

### Data analysis

#### Mouse position tracking

We used the color marker tracking function in the social interaction module (NSE-EV-SIM) of EthoVision XT 16.0, calibrated for precise mouse tracking under experimental conditions. Each mouse was uniquely identified by a 1.3 cm cube-shaped color marker, made of hardboard paper, featuring a distinct color and a printed identification number, and securely attached to the CBRAIN headstage. EthoVision XT 16.0 recorded spatial coordinates (cm) at each timepoint (s), allowing detailed analysis of spatial and temporal activity patterns during foraging experiments.

#### Space occupancy

The Colosseum cage was broadly divided into the Robot zone and the Residence zone. To further analysis, the Residence zone was further subdivided into three zones: (1) the Near zone, the region extending 10 cm from the Spider zone wall, (2) the Gate zone, which spanned 30° around each gate, and (3) the Nest zone, occupying approximately 35° of the Residence zone. Space occupancy was determined by measuring the time each mouse spent in these defined areas. Mouse positions, recorded in Cartesian coordinates, were converted to polar coordinates, and the time spent in each region was used to calculate space occupancy.

#### Kinetic energy

Kinetic energy at each time was calculated as *KE* = *mv*^2^⁄2, where *m* is the mouse’s weight on a given day, and *v* is the instantaneous speed, determined by the distance traveled between tracked positions divided by the time interval.

#### Eating cluster and snack consumption estimation

Estimating snack consumption is challenging due to the irregular shape of the fish snack. However, we observed a consistent behavioral pattern – mice always gathered around the snack while eating. Based on this observation, we estimated snack consumption by measuring the time each mouse spent within the eating cluster. This approach was based on two key behavioral tendencies: (1) when a mouse attempted to take the snack and run away alone, it consumed very little, and (2) once a mouse began eating alone, other mice quickly converged, forming a cluster and sharing the snack. Given this strong tendency for collective eating, we used the time spent within the eating cluster as a proxy for snack consumption, referred as 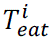.

To systematically identify eating cluster, we applied Density-Based Spatial Clustering of Applications with Noise (DBSCAN)^8^ using MATLAB’s built-in *dbscan.m* function. A search radius of 6 cm(approximately the body size of a mouse) and a minimum neighbor count of four were set as clustering parameters. Clusters containing at least two core points and peristing for more than three seconds were classified as potential eating clusters. To ensure accuracy, all detected eating clusters underwent manual verification through video analysis, confirming that the aggregation reflected actual eating behavior.

#### Latency analysis

Latency analysis was conducted to quantify the temporal dynamics of key behaivoral events during the experiment. Event timing was defined based on critical moments captured in the video recordings. The **Start of the experiment**, *t_start_* was defined as the moment the experimenter disappeared from the video frame after placing a snack on the robot. The **entry moment**, 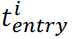 was identified as the first instance when the *i*^th^ mouse inserted its head and forelimb into the robot zone. The **ride moment**, 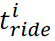 was identified as the first instance when the *i*^th^ mouse grasped the spider robot and maintained its hold without releasing. The **get moment**, *t_get_* was defined as the moment when the worker mouse obtained the snack, identified by the moment when the snack was no longer overlapping with the robot in the video. The **delivery moment**, *t_deliver_* corresponded to the point at which a mouse holding the snack completely exited the robot zone. The **eat moment**, 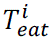 was determined as the first instance when the *i*^th^ mouse entered the eating cluster. Finally, the **end moment**, *t_end_* was defined as the moment when all mcie had finished eating and dispersed from the eating cluster.

Event durations were calculated to assess latency and response times for different phases of the experiment. The **latency of action**, 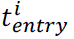 was computed as the time elapsed between the start of the experiment and the entry moment (i.e., 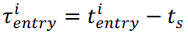). If the *i*^th^ mouse did not enter the robot zone, its latency was considered infinite (i.e., 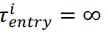). The **latency to get the snack**, 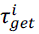 was measured as the time between the entry moment and the get moment (i.e., 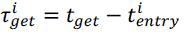), with infinite latency assigned to mice that never entered the robot zone. The **latency for eating**, 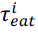 was defined as the time gap between delivery moment and the eat moment (i.e., 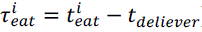).

Behavior periods are defined as follows: the period from the start of the experiment to entry was defined as **Pre-act**, the period between entry and ride as **Act**, the period between ride and get as **Retrieve**, the period between get and eat as **Delivery**, and the period from the start to the end of eating as **Eat**.

#### Role assignment

To classify individual mice based on their behavior in group foraging tasks, we assigned roles according to both behavioral actions and individual performance metrics. Each mouse was categorized into one of three behavioral roles based on its actions during foraging. **Working mice** were defined as individuals that entered the robot zone and successfully retrieved the snack from the robot. **Participating mice** entered the robot zone but did not retrieve the snack. **Freeriding mice** did not enter the robot zone until the snack had been retrieved by another mouse. Freeriding behavior was further divided into **engaged freeriding**, where the mouse remained active in the proximal robot zone or gate zones while waiting for the snack, and **disengaged freeriding**, where the mouse was either stationary or remained in the nest or residence zones. Additionally, **ride behavior**, defined as a mouse riding on the moving robot while holding onto it, was noted but not used as a primary criterion for role assignment. **Daily working, participating, and freeriding rates** were assessed for each mouse by calculating the proportion of trials per day in which they exhibited the respective behaviors.

For role assignment, the **accumulated work rate** was a key metric, calculated as the number of trials in which a mouse exhibited working behavior divided by the total number of group foraging trials. Mice were classified based on their individual performance across trials. The **Top worker** was the mouse with the highest accumulated work rate for at least three consecutive days. **Participants** exceeded the work chance level, the expected contribution under equal workload distribution, while **Freeriders** did not.

#### Role transition

To analyze individual’ role transition among working (W), participating (P), and freeriding (F) role, we employed a Markov chain approach. First, a role array was constructed for each mouse, representing its assigned role at each trial, and then we generated a discrete-time Markov chain (DTMC) with MATLAB’s *dtmc.m* function, which produced a normalized transition probability matrix, quantifying the standardized probabilities of role transitions. To examine group-level role transitions for Top Workers, Participants, and Freeriders, we averaged the transition probabilities of individuals within each category and applied the same DTMC-based procedure to derive the normalized transition probability matrix for each role group.

As trials progressed, we defined the probability of transitioning toward increased work engagement as the probability of more work and the probability of transitioning toward reduced work engagement as the probability of less work. These were computed as the sum of transition probabilities from F → W, F → P, P → W, and W → W for more work, and the sum of transition probabilities from W → P, W → F, P → F, and F → F for less work, respectively.

#### Lorentz curve and Gini index

To analyze the fairness of resource allocation in relation to common resource (i.e., snack) acquisition and distribution, we incorporated behavioral economic variable. Specifically, we applied the Boltzmann fair division for distributive justice (PMID: 36171244), assuming mice receive compensation proportional to their work. Under this framework, the payroll of *i*^th^ mouse (*P_i_*) was calculated by

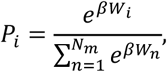

where *W_i_* represents the cumulative number of trials in which the *i*^th^ mouse engaged in working behavior, β is a scaling factor, *N_m_* is the mouse group size. Here, we determined the value of β based on the assumption that if only one worker contributed to foraging, a fair resource distribution would result in an 80:20 split between the worker and non-workers:

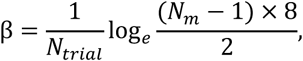

where *N_trial_* is the total number of trials.

Likewise, the exact value of resource share, *E_i_* can be estimated by the duration in the eating time 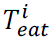,

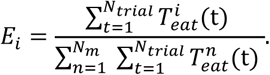

However, here, as 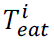 did not show statistically significant differences among mice, we used 1⁄*N_m_* for simplicity. Then the Lorenz curve^9^ was plotted with the cumulative *P_i_* values, sorted in ascending order, on the y-axis, and the cumulative *E_i_* on the x-axis. The Gini coefficient^10^, by definition, the area between the line of equality and the Lorenz curve divided by 0.5.

#### Delineation of social dominance rank

Social dominance rank assessed using the tube test, was determined based on the win rate. However, this method can lead to tied ranks, where multiple mice share the same ranking. To resolve this issue, we implemented the ELO scoring system^11^, which assign greater weight to wins against higher-ranked opponents. First, ranks were assigned based on the total number of wins. In cases of ties, mice were further ranked in ascending order of the cumulative ranks of their defeated opponents, calculated as sum of the ranks of all defeated opponents. If cumulative opponent ranks were also identical, precedence was given to mice that had defeated the highest-ranked opponent. The code used is available at: https://github.com/jeelabKIST/GroupForaging/.

#### LFP analysis

For preprocessing, the LFP signals collected using CBRAIN telemetry were high-pass filtered at 1 Hz and low-pass filtered at 300 Hz with a band-stop filter at 60 Hz. Movement artifacts were detected using amplitude thresholding and manually verified for accuracy before being rejected. The preprocessed LFP signals were transformed into spectral powers using MATLAB’s spectrogram function with a 0.25 s window and 0.125 s overlap. Spectral powers for each trial were z-score normalized by subtracting the mean and dividing by the standard deviation of the spectral powers at each frequency across the entire trial. For the time-normalized spectrogram, an equal number of time points were sampled for each behavioral period (e.g., Pre-act, Act). The time between the start and end points of a period in a given trial was divided into equidistant intervals, and the spectral powers closest to these intervals points were collected. After repeating this process for each trial, the samples were stacked and averaged to produce a time-normalized spectral representation of the foraging trials. The frequencies of interests are set as low theta (4–8 Hz), high theta (8–12 Hz), low beta (18–24 Hz), high beta (24–32 Hz), low gamma (35–50 Hz), and high gamma (70–90 Hz) frequency bands. The coherence of two signals were calculated with *cohere.m* function in MATLAB.

To extract transient oscillatory bursts, we followed the burst detection method previously reported in Cho et al., 2022. The time series-based burst detection algorithm was used to extract burst periods from narrow band-passed LFP traces (*detect_burst_timeseries.m*). The algorithm was run by setting amplitude threshold factor to 2.3 for all frequency bands. In burst classification, a burst detected at only one site was classified as a local burst, whereas bursts detected simultaneously at two or more sites were categorized as a network burst. The occurrence rate of local or network burst was estimated as the reciprocal of the inter-burst interval.

For analyzing connectivity of network bursts, we calculated the phase locking value (PLV)^12^. First, the instantaneous phase of LFPs were calculated via Hilbert transform using MATLAB built-in hilbert.m function after bandpass filtering and then unwrapped. Within the time of interest, PLV was calculated by 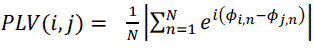, where ϕ*_i,n_* and ϕ*_j,n_* are the instantaneous phases of *i*^th^ and *j*^th^ LFP signals at time n. PLV of 1 indicates perfect phase locking, whereas PLV of 0 indicates absence of phase locking. Only the time intervals containing co-occurred bursts were used for calculating PLV.

To determine the directionality of network bursts, we calculated the directed Phase Lag Index (dPLI)^13^, defined as 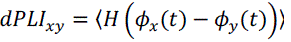, where *H* is the Heaviside step function, and 〈·〉 denotes the expectation over time. A *dPLI_xy_* of 0.5 indicates that neither *x* or *y* leads or lags on average. A *dPLI_xy_* > 0.5 indicates signal *x* is phase-leading compared to signal *y*, whereas *dPLI_xy_* < 0.5 indicates *x* is phase-lagging relative to *y*.

#### Statistics

Statistical analyses were performed with MATLAB. Prior to statistical tests, the normality of data was tested with the Shapiro-Wilk and Kolmogorov-Smirnov tests. In the case of highly skewed data, the best-fitting distribution function for each dataset was determined by testing its goodness-of-fit. The statistical power was validated by comparing the test results of the original data with those generated from the fitted distribution. If a significant discrepancy was found between the two, the statistical results from this study were rejected. For statistical analysis, the Mann-Whitney U test was used for pairwise comparisons, and the Kruskal-Wallis test with post hoc multiple comparisons was applied for comparisons involving more than three groups. A significance level of 0.05 was used to determine statistical significance, and adjustments for multiple comparisons were made using the Bonferroni correction. For regression analyses, Pearson’s correlation was used for continuous data, while Spearman’s correlation was applied to ordinal data. The statistical distance between two distributions, *p*(*x*) and *q*(*x*) was assessed with Kullback-Leibler divergence, *D_KL_* defined as 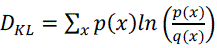. Data are presented as mean ± SEM.

### Histology

After completing all behavior experiments, electrode implantation sites were verified through electrolytic lesions (2 mA, 5 s), to confirm the precise location of the electrodes. Perfused brains were fixed overnight (12–16 hours) at 4℃ in 4% paraformaldehyde (PFA) in PBS and then cryoprotected in 30% sucrose in PBS for 48 hours at 4℃. After freezing, the brains were sectioned into 50 μm slices using a cryostat and mounted using Fluoroshield with DAPI mounting medium (Catalogue number: 0000288392, SIGMA-ALDRICH Co., USA). Images of the PFC, NAc, and BLA regions were captured using a confocal microscope at 2.5x magnification (AxioCam MRc, ZEISS, Germany) and acquired using Stereo Investigator software (MicroBrightField, Inc., Colchester, VT, USA).

## Supplementary Information

Extended Data 1–6

Movie S1–S2

Supplementary Figures 1–9

Supplementary Tables 1–8

**Extended Data 1.**
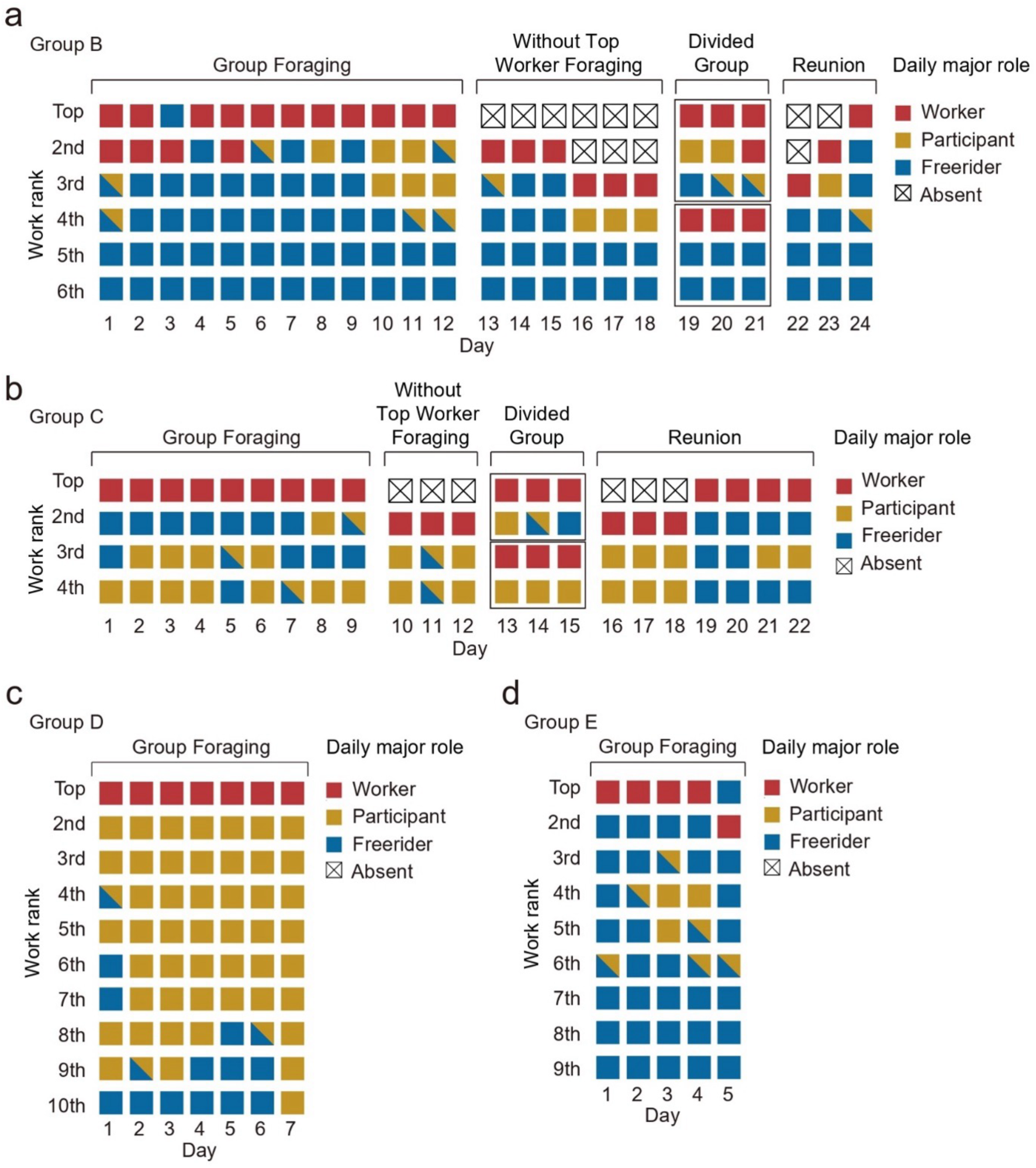
Daily role assignments in group B-E. **a-d**, Daily role assignments across experimental phases. Daily roles were tracked throughout different experimental phases, including group foraging in groups B–E, foraging without the top worker, subgrouping, and reunion in groups B and C.

**Extended Data 2.**
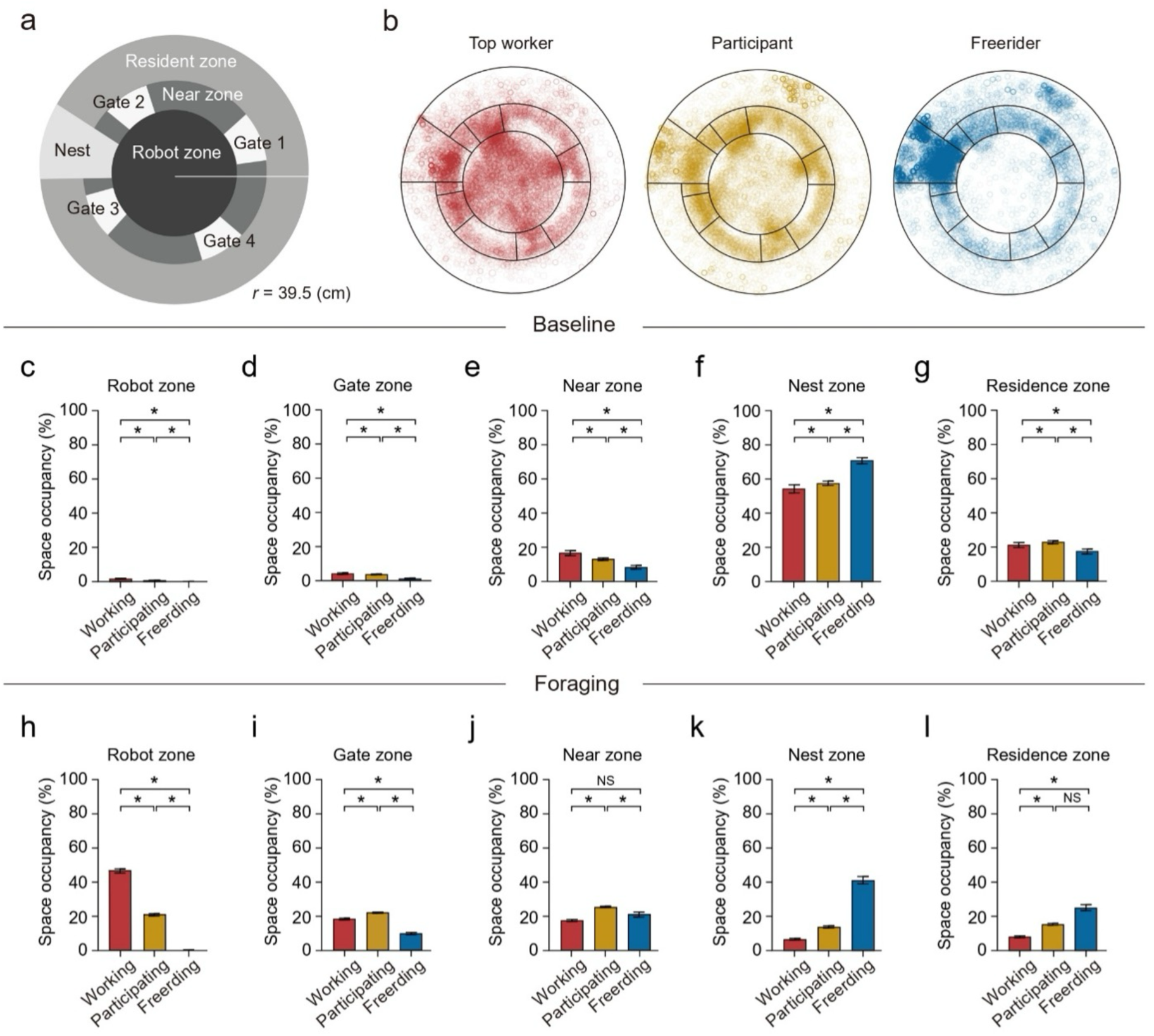
Spatial distribution and zone occupancy across behavioral roles. **a**, Spatial map of the colosseum cage, divided into Robot, Near, Gate, Resident, and Nest zones (see **Methods**). **b**, Representative spatial occupancy heat maps for top worker (red), participant (yellow), and freerider (blue) across all trials (159 trials) during group foraging. **c–g**, Percentages of time spent in each zone during the baseline phase (working, W: n = 159; participating, P: n = 542; freeriding, F: n = 253). Kruskal–Wallis tests with post hoc Dunn’s comparisons were used throughout. **c**, Robot zone: χ²(2,954) = 86.989, *p* < 0.001 for all pairwise comparisons. **d**, Gate zone: χ²(2,954) = 96.337, *p* < 0.001; P vs. F and W vs. F, *p* < 0.001; W vs. P, *p* = 0.024. **e**, Near zone: χ²(2,954) = 69.777, *p* < 0.001; W vs. F and P vs. F, *p* < 0.001; W vs. P, *p* = 0.029. **f**, Nest zone: χ²(2,954) = 49.107, *p* < 0.001; P vs. F and W vs. F, *p* < 0.001; W vs. P, *p* = 0.596. **g**, Residence zone: χ²(2,954) = 27.597, *p* < 0.001; P vs. F and W vs. F, *p* < 0.001; W vs. P, *p* > 0.999. **h–l**, Percentages of time spent in each zone during the foraging phase. **h**, Robot zone: χ²(2,954) = 632.152, *p* < 0.001; *p* < 0.001 for all pairwise comparisons. **i**, Gate zone: χ²(2,954) = 213.891, *p* < 0.001; *p* < 0.001 for all pairwise comparisons. **j**, Near zone: χ²(2,954) = 90.285, *p* < 0.001; W vs. P and P vs. F, *p* < 0.001; W vs. F, *p* = 0.176. **k**, Nest zone: χ²(2,954) = 156.232, *p* < 0.001; *p* < 0.001 for all pairwise comparisons. **l**, Residence zone: χ²(2,954) = 55.462, *p* < 0.001; W vs. P and W vs. F, *p* < 0.001; P vs. F, *p* = 0.729. Error bars represent mean ± s.e.m. \**p* < 0.05; NS, not significant.

**Extended Data 3.**
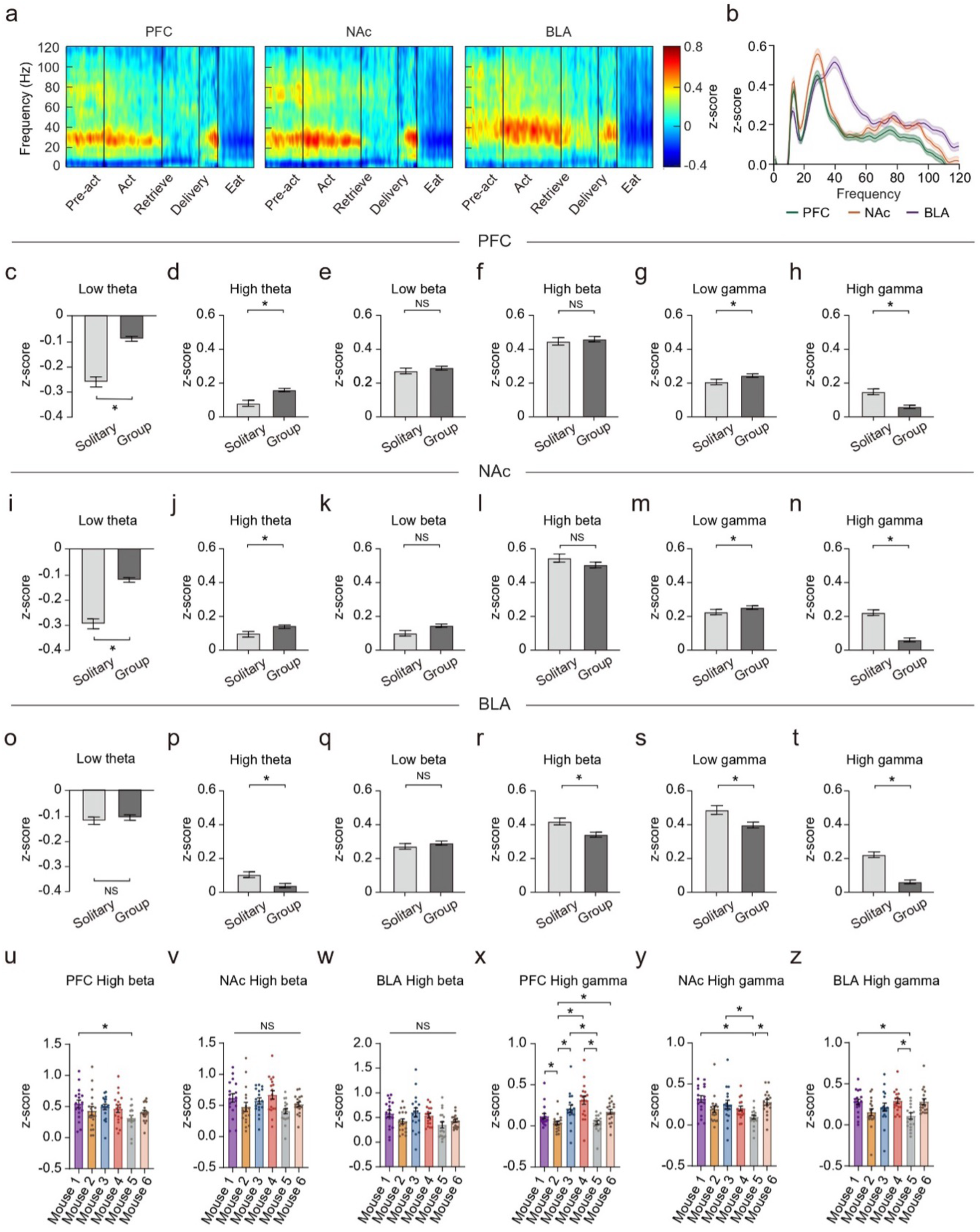
Neural oscillatory power during solitary foraging. **a**, Time-normalized, z-scored power spectrograms in the mPFC, NAc, and BLA from pre-Act to Eat phases during solitary foraging. **b**, Z-scored spectral power distributions across these regions during the Act period. **c–t**, Comparison of mean z-scored power in solitary versus group foraging across frequency bands and brain regions during Act. In the mPFC, low theta (**c**, U = 4434, *p* < 0.001), high theta (**d**, U = 5978, *p* < 0.001), and low gamma (**g**, U = 6697, *p* = 0.001) are significantly higher in group, whereas high gamma (**h**, W = 5857, *p* < 0.001) is elevated in solitary. Low beta (**e**, W = 8015, *p* = 0.243) and high beta (**f**, U = 8135, *p* = 0.329) do not differ. In the NAc, low theta (**i**, U = 4061, *p* < 0.001), high theta (**j**, U = 7187, *p* = 0.013), and low gamma (**m**, U = 7211, *p* = 0.014) are higher in group, whereas high gamma (**n**, U = 4090, *p* < 0.001) is higher in solitary. Low beta (**k**, W = 8717, *p* = 0.9613) and high beta (**l**, U = 8115, *p* = 0.314) do not differ. In the BLA, high theta (**p**, W = 7052, *p* = 0.007), high beta (**r**, U = 6869, *p* = 0.002), low gamma (**s**, U = 7392, *p* = 0.031), and high gamma (**t**, U = 3528, *p* < 0.001) are significantly higher in solitary. Low theta (**o**, U = 8532, *p* = 0.731) and low beta (**q**, U=8554, *p* = 0.758) do not differ. **u–z**, Comparison of high-beta and high-gamma power across individual mice (n = 18 solitary foraging trials per mouse). In the mPFC (**u**, χ²(5,108) = 12.15, *p* = 0.032), high-beta differs only between Mouse 1 and Mouse 5 (*p* < 0.041), whereas no significant difference is observed in the NAc (**v**, χ²(5,108) = 14.4, *p* = 0.013) or BLA (**w**, χ²(5,108) = 5.855, *p* = 0.32). High gamma (**x–z**) shows significant between-mouse differences, though the pattern varies by region (see text for pairwise comparisons). Error bars represent mean ± s.e.m. Statistical significance is based on Mann–Whitney U tests for two-sample comparisons, with post hoc Dunn’s tests following Kruskal–Wallis. \**p* < 0.05; NS, not significant.

**Extended Data 4.**
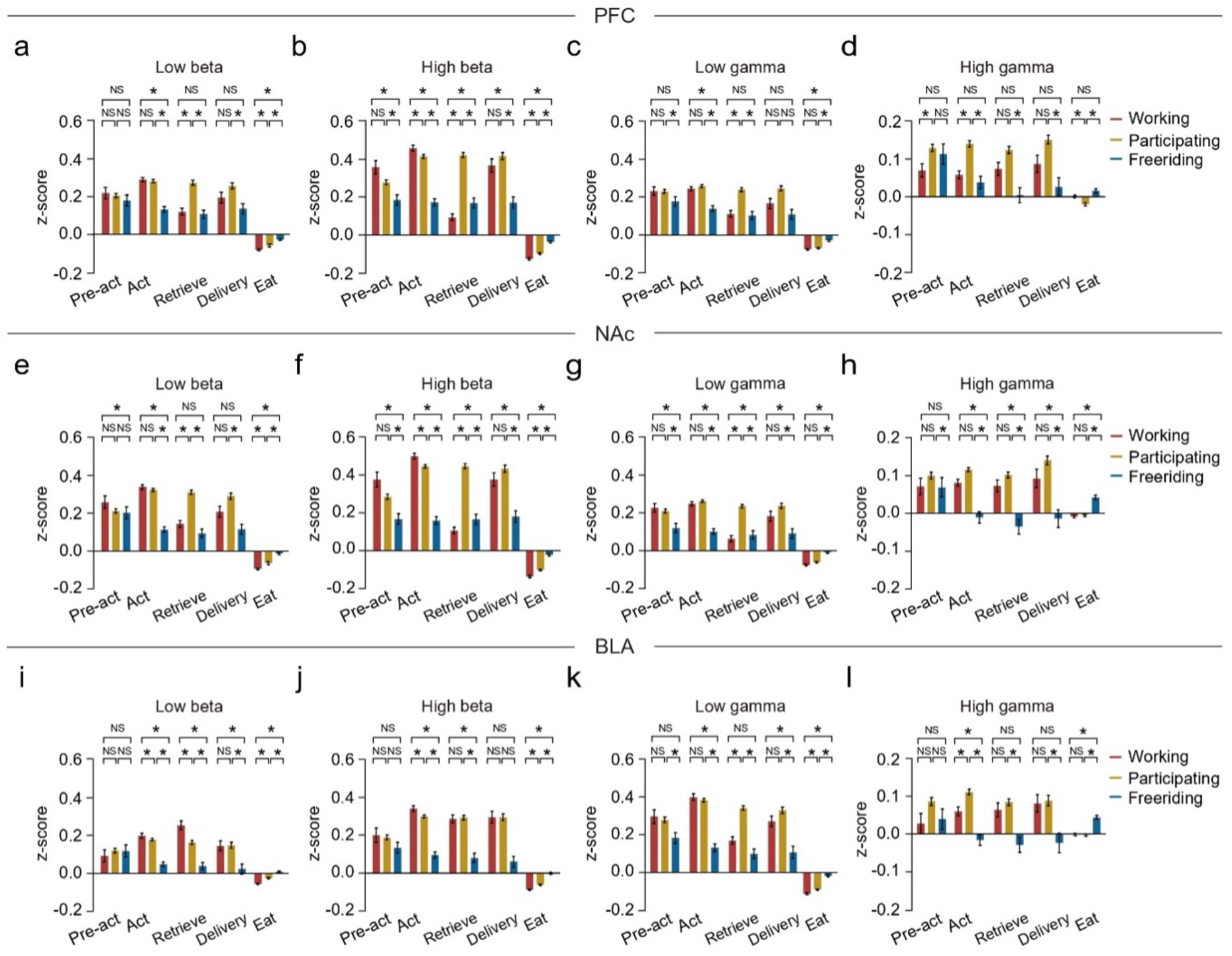
Neural oscillatory power across behavioral periods. **a–l**, Mean z-scored power for low beta, high beta, low gamma, and high gamma in the mPFC (**a–d**), NAc (**e–h**), and BLA (**i–l**) during specific behavioral periods. Error bars represent mean ± s.e.m. \**p* < 0.05; NS, not significant. See Supplementary Table 6 for full statistical details.

**Extended Data 5.**
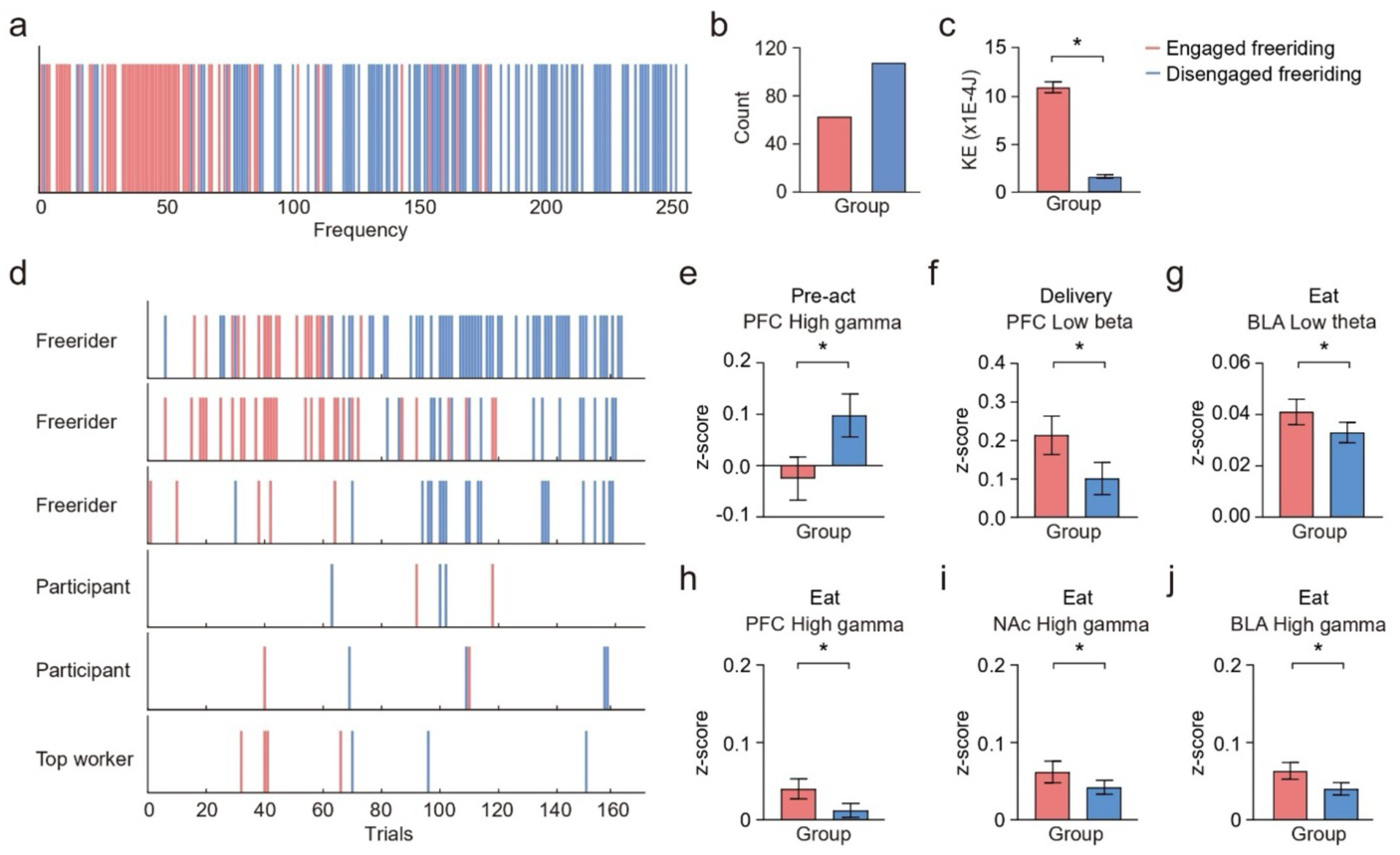
Behavioral and neural differences between engaged and disengaged freeriding. **a**, Temporal distribution of engaged (light red) and disengaged (light blue) freeriding behaviors across mice (see **Methods**). **b**, Total trials of engaged (62 trials) vs. disengaged (107 trials) freeriding. **c**, Kinetic energy is higher in engaged vs. disengaged freeriding (U = 193, *p* < 0.001; Mann– Whitney U). **d**, Role-based individual freeriding patterns over trials. **e–g**, Z-scored spectral activity during different behavioral periods for engaged (61 trials) and disengaged (106 trials) freeriding, except e (disengaged trials). Disengaged freeriders have higher high-gamma power in the mPFC during pre-Act (**e**, U = 2348, *p* = 0.005). In contrast, engaged freeriders show higher low beta power in the mPFC during Delivery (**f**, U = 2637, *p* = 0.047) and higher low theta power in the BLA during Eat (**g**, U = 2607, *p* = 0.037). **h–j**, Engaged freeriders also exhibit higher high gamma power in the mPFC (**h**, U = 2424, *p* = 0.007), NAc (**i**, U = 2571, *p* = 0.028), and BLA (**j**, U = 2404, *p* = 0.006). Error bars represent mean ± s.e.m. \**p* < 0.05.

**Extended Data 6.**
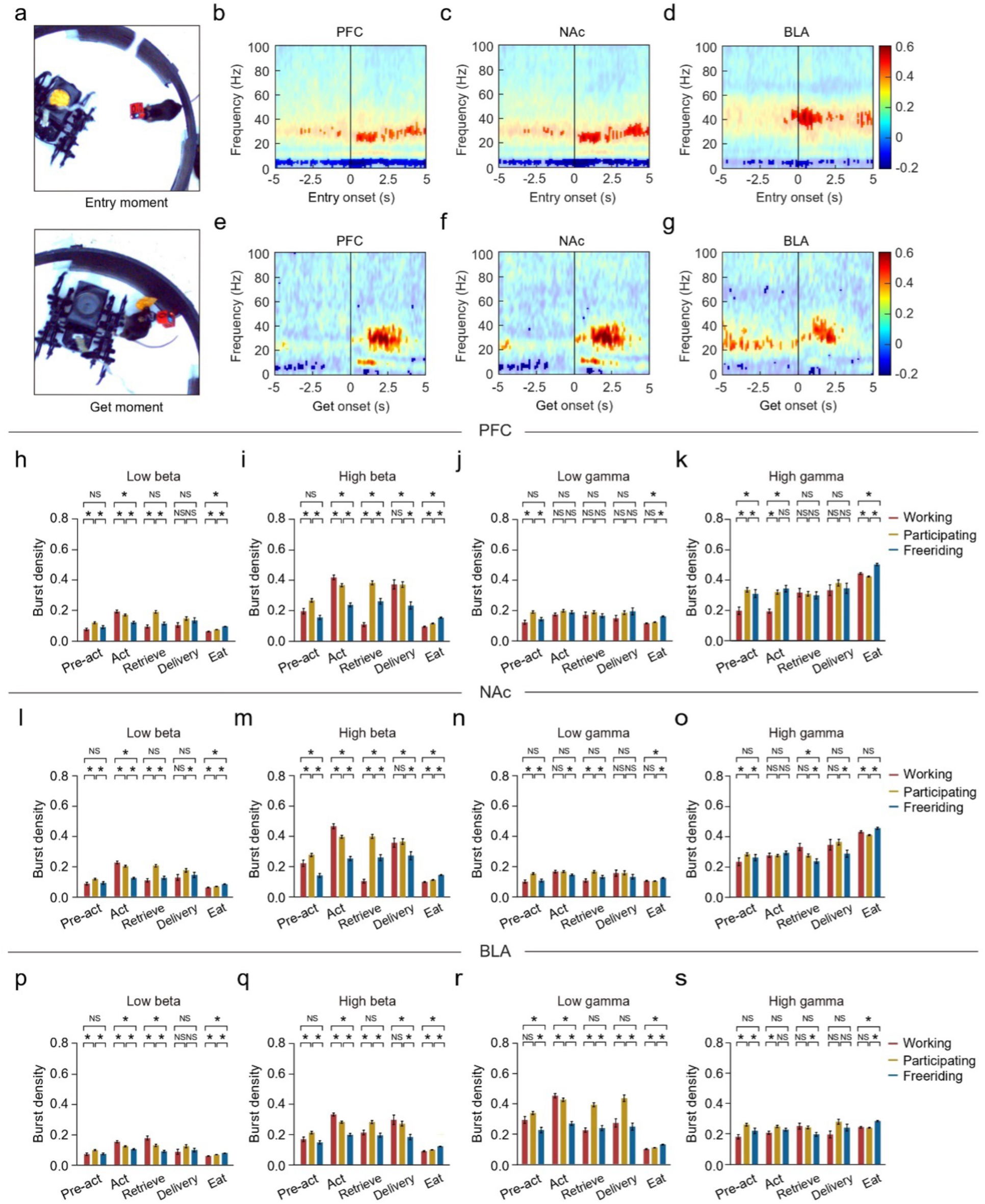
Burst density across behavioral periods. **a**, Schematic of the Entry point (top) and Get point (bottom). **b–g**, Time-frequency spectrograms (z-scored) aligned to behavioral events, where red shading indicates significant increases and blue shading indicates decreases (*p* < 0.05; ± 1.96σ). After the Entry point, beta power rises in the PFC (**b**) and NAc (**c**), while low gamma power increases in the BLA (**d**). After the Get point, beta power increases in the PFC (**e**) and NAc (**f**), and low gamma power increases in the BLA (**g**). **h–s**, Mean burst density for low beta, high beta, low gamma, and high gamma in the PFC (**h– k**), NAc (**l–o**), and BLA (**p–s**) during specific behavioral periods. Error bars represent mean ± s.e.m. *p < 0.05; NS, not significant. See **Supplementary Table 9** for full statistical details.

**Movie S1.**
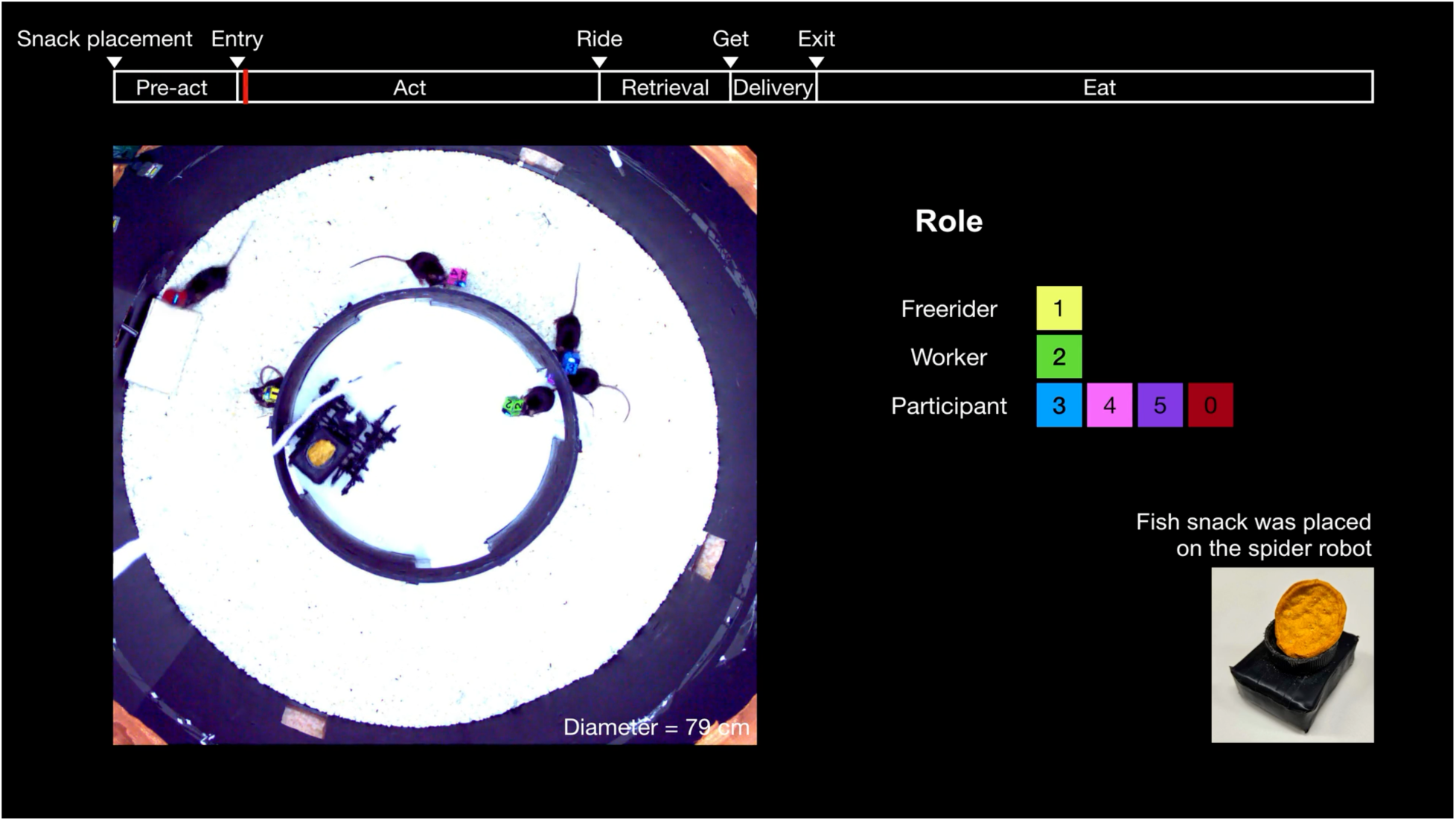
Group foraging. Example of a group foraging trial. Bar on the top describes the progression of time across defined behavioral moments and periods. In the video, six mice have distinct color blocks attached aside the CBRAIN headstage. The exmaple trial involves one worker, four participants and one freerider.

**Movie S2.**
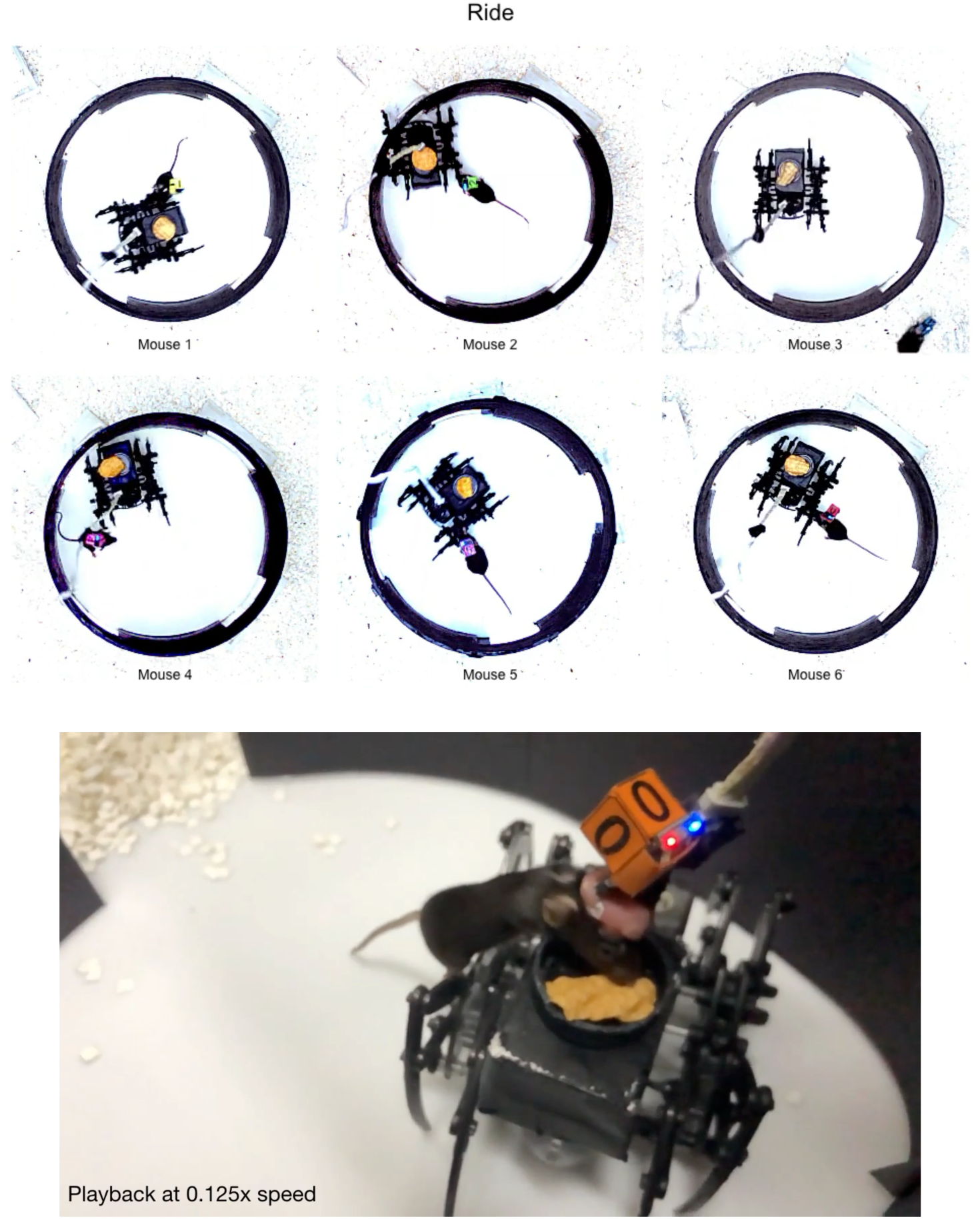
Solitary foraging. Example of solitary foraging trials performed by individual mice from group A. Videos are played side-by-side, showing one behavioral period at a time. Appended to the solitary foraging videos is a side-view video of a mouse climbing to a spider robot and retrieving the fish snack.

**Supplementary Figure. 1.**
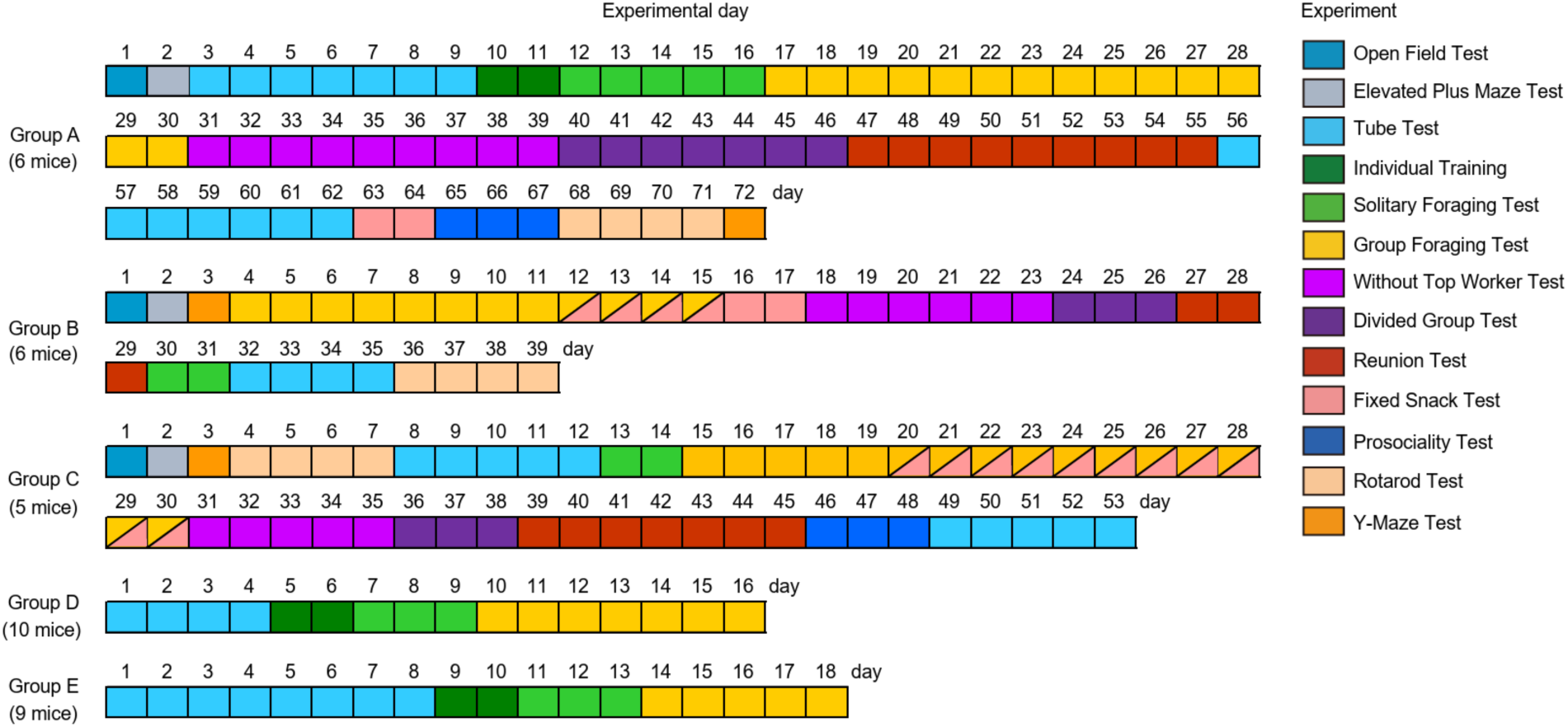
Experimental timelines for groups A–E. Visualization of comprehensive summary of the experimental flow and the progression of tests conducted over consecutive days. Schedules for five experimental groups (Group A – E) across days. Each block represents a day, with colors indicating the type of experiment conducted on that day. Experimental days are sequentially numbered for each group. For different types of experiments, refer to the methods.

**Supplementary Figure 2.**
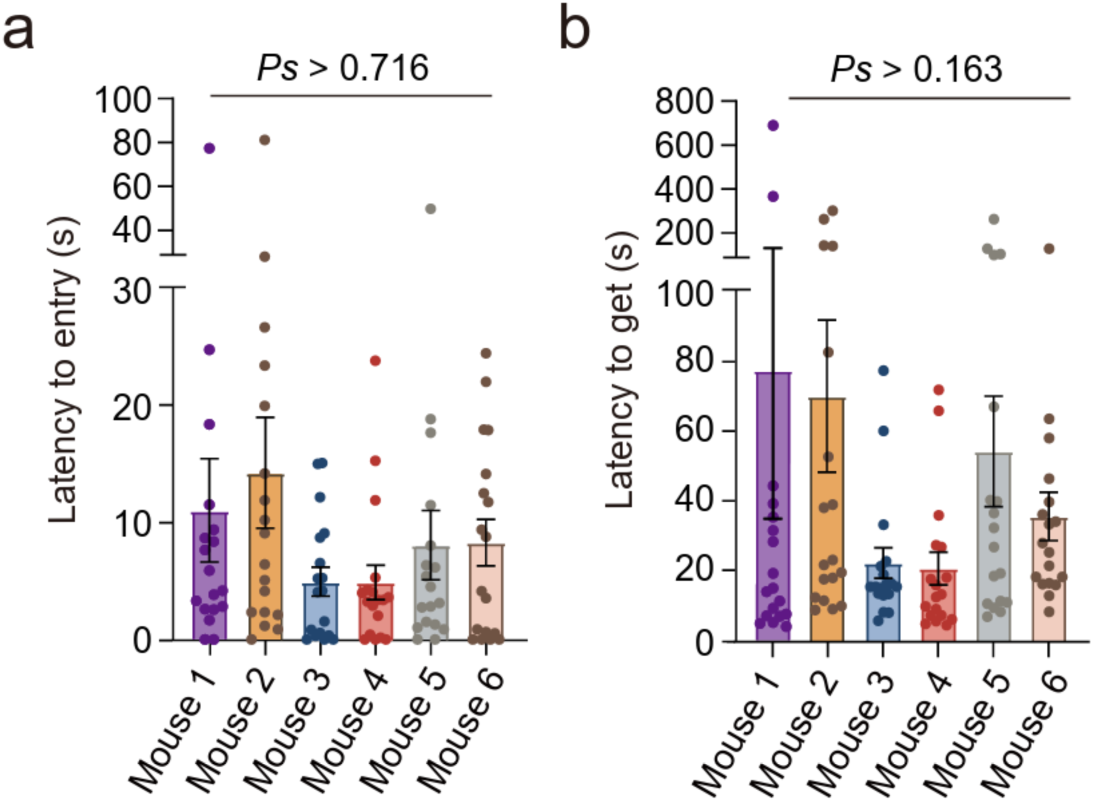
Latency to entry and get the snack during solitary foraging. Latency to entry and get were measured to assess each individual’s foraging ability. All mice from group A completed 18 solitary foraging trials before the group foraging experiment began. **a**, Latency to entry did not differ significantly across individuals (χ²(5, 108) = 5.364, *p* = 0.373, *ps* > 0.716). **b**, Latency to get showed no significant difference across individuals (χ²(5, 108) = 12.47, *p* = 0.028, *ps* > 0.163). Error bars represent the mean ± s.e.m., and dots indicate individual data points. There were no significant differences in entry and retrieval latency among individuals during solitary foraging.

**Supplementary Figure 3.**
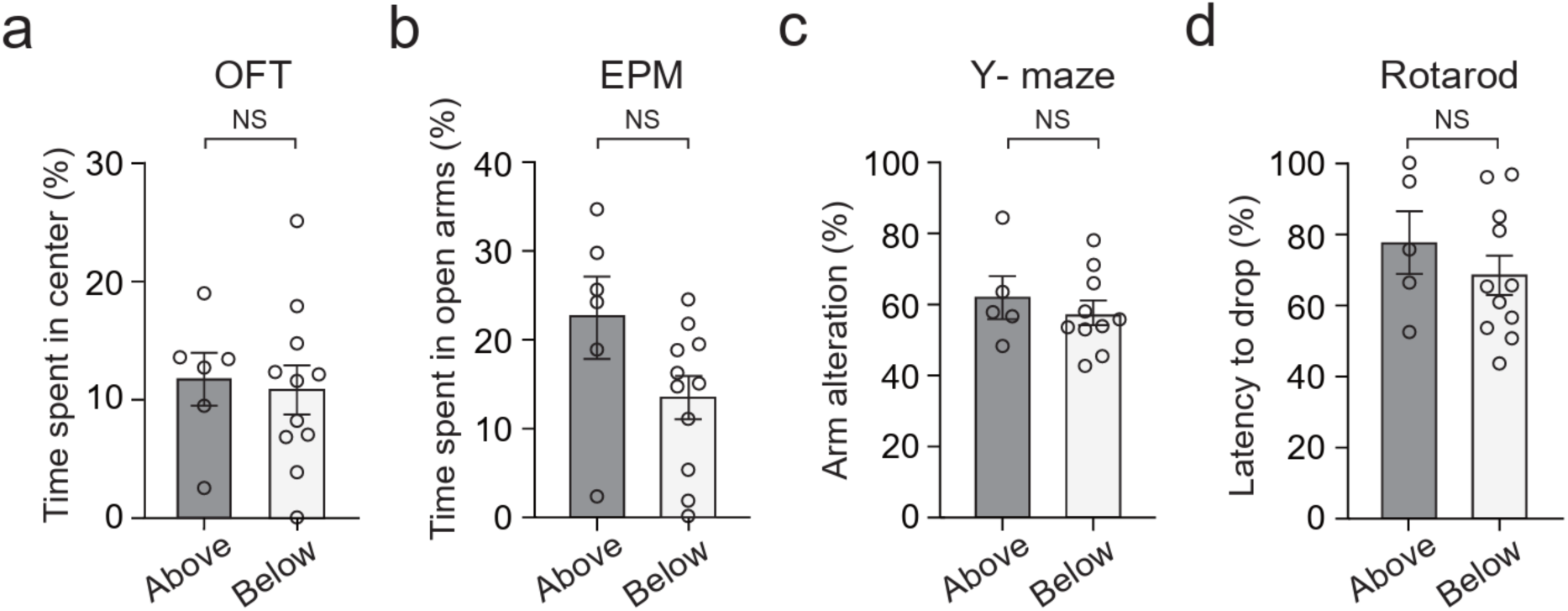
No significant association found between work rate and individual traits. The Open Field Test (OFT), Elevated Plus Maze (EPM), Y-maze, and Rotarod tests were performed to assess the relation between work rates and individual traits. The Above group (dark gray) includes individuals with a work rate exceeding the expected chance level, while the Below group (light gray) consists of individuals with a work rate lower than the expected chance level (methods). There were no significant differences between the Above and Below groups overall individual traits. **a–b**, Anxiety-like behavior (Above: *n =* 6 mice, Below: *n =* 11 mice) estimated by OFT (**a**, *t*(15) = 0.288, *p* = 0.777) and EPM (**b**, *t*(15) =1.98, *p* = 0.065). **c**, Working memory estimated by Y-maze (Above: *n =* 5 mice, Below: *n =* 10 mice); *t*(13) = 0.805, *p* = 0.434), and **d**, Motor coordination estimated by Rotarod test (Above: *n =* 5 mice, Below: *n =* 11 mice); *t*(14) = 0.921, *p* = 0.372, Unpaired two-tailed *t*-test). Bar graphs represent the mean ± s.e.m and dots indicate individual data. NS denotes that no significant statistical difference was found.

**Supplementary Figure 4.**
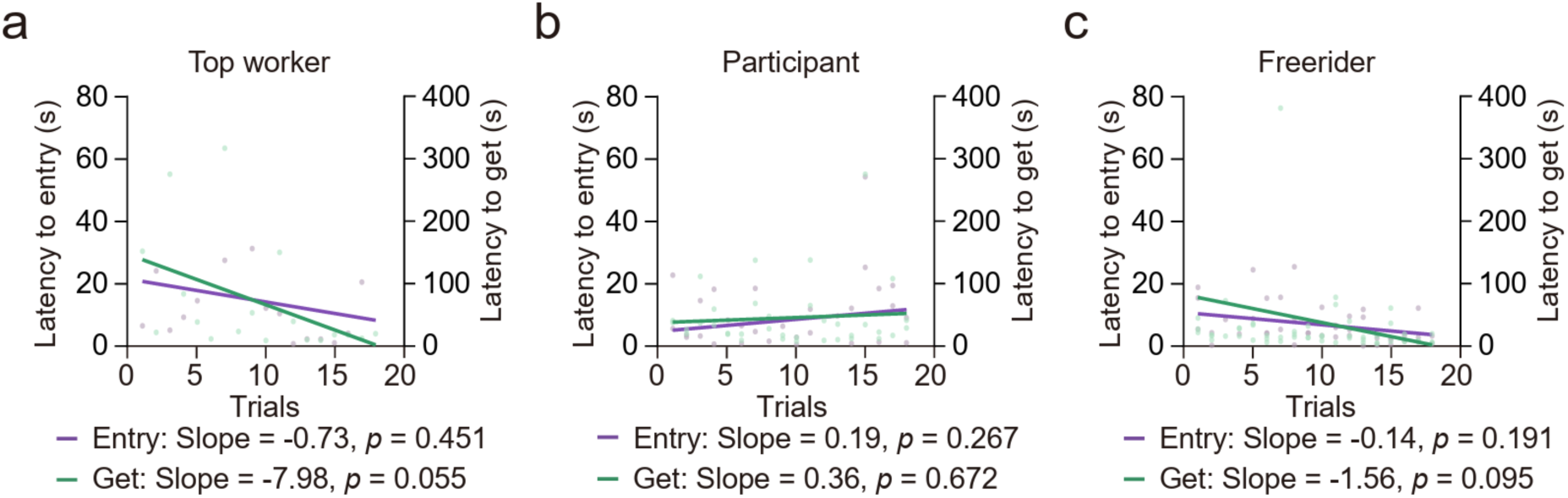
No significant trends in entry and retrieval latency across trials in solitary conditions. To confirm whether individual foraging behavior changed with repeated trials in the solitary condition, we performed a linear regression analysis on entry latency and retrieval time. No significant trends were observed in the latency to Entry and Get among **a**, Top worker (*n* = 1 mouse, 18 trials, Entry: slope = –0.73, R² = 0.035, *p* = 0.451, Get: slope = –7.98, R² = 0.209, *p* = 0.055), **b**, Participant (*n* = 2 mice, 36 trials, Entry: slope = 0.19, R² = 0.036, *p* = 0.267, Get: slope = 0.36, R² = 0.005, *p* = 0.672) and **c**, Freeriders (*n* = 3 mice, 54 trials, Entry: slope = –0.14, R² = 0.032, *p* = 0.191, Get: slope = –1.56, R² = 0.052, *p* = 0.095). The lines represent linear regression trends, while the dots indicate individual trial data points.

**Supplementary Figure 5.**
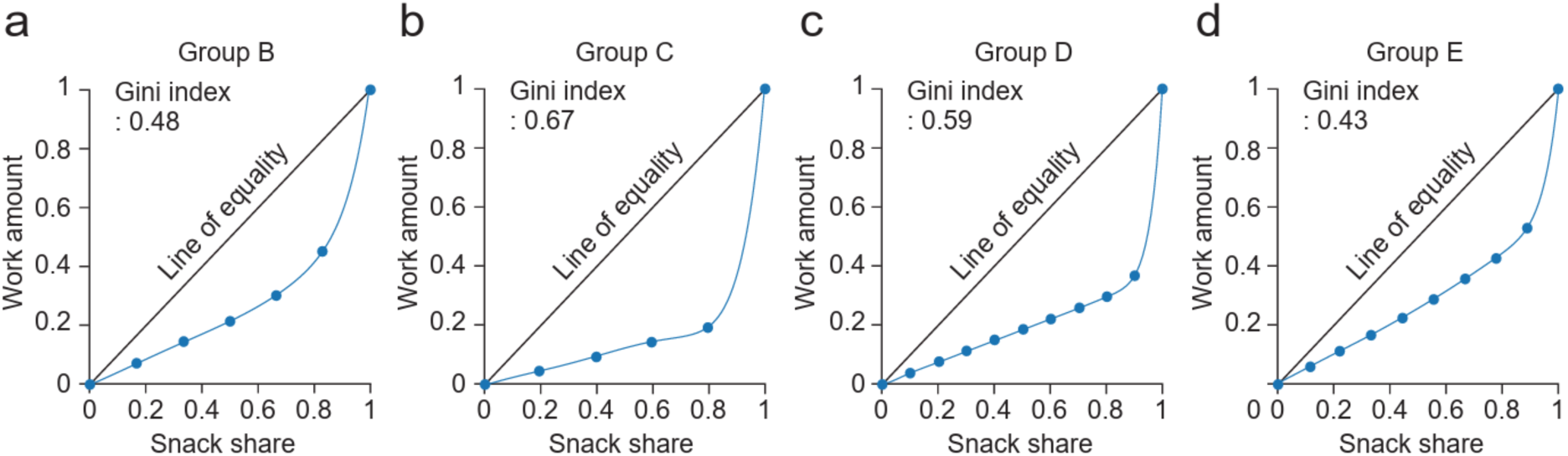
Work imbalance in different groups. Lorenz curve-based analysis of work imbalance: the diagonal line indicates perfect equity between normalized cumulative work contribution and snack share, with deviations from this ideal balance captured by a 10th-degree polynomial fit (blue). **a**, Group B (Gini index: 0.48), **b**, Group C (Gini index: 0.67), **c**, Group D (Gini index: 0.59), and **d**, Group E (Gini index: 0.43). While the degree of work imbalance varies across groups, all exhibit deviations from perfect equity, reflecting unequal distribution in relation to contribution.

**Supplementary Figure 6.**
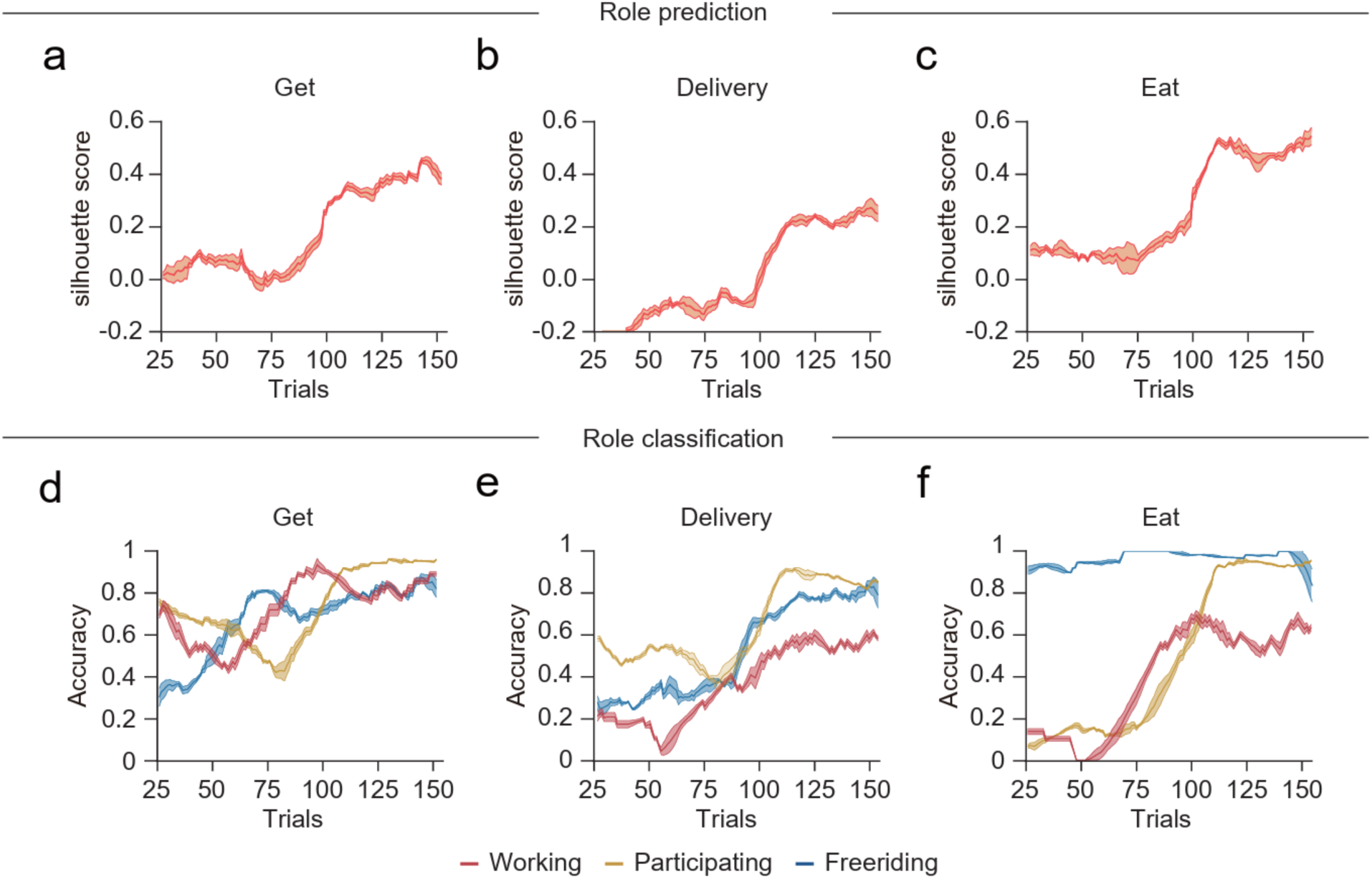
Role classification and prediction accuracy during Get, Delivery, and Eat period. As in Figure 3f-g, role classification accuracy was assessed using silhouette scores, with higher values indicating better separation. Role classification accuracy increases over trials. **a**, Get period: mean final silhouette score over the last 20 trials was 0.412 ± 0.006. **b**, Delivery period (0.243 ± 0.005) and **c**, Eat period (0.478 ± 0.007). As in Figure.3 h-i, accuracy trend shows changes in role prediction performance across trials. **d**, Get period: Mean final scores over the last 20 trials increased for working (0.846 ± 0.008), participating (0.958 ± 0.001), and freeriding (0.833 ± 0.007). **e**, Delivery period: Role prediction gradually improved, with final scores for working (0.563 ± 0.005), participating (0.848 ± 0.004), and freeriding (0.799 ± 0.004). **f**, Eat period: Working (0.608 ± 0.010) and participating (0.934 ± 0.002) mice show gradual increases, whereas freeriding mice (0.961 ± 0.011) showed no change. The line shows the mean, and the shaded area represents the s.e.m.

**Supplementary Figure 7.**
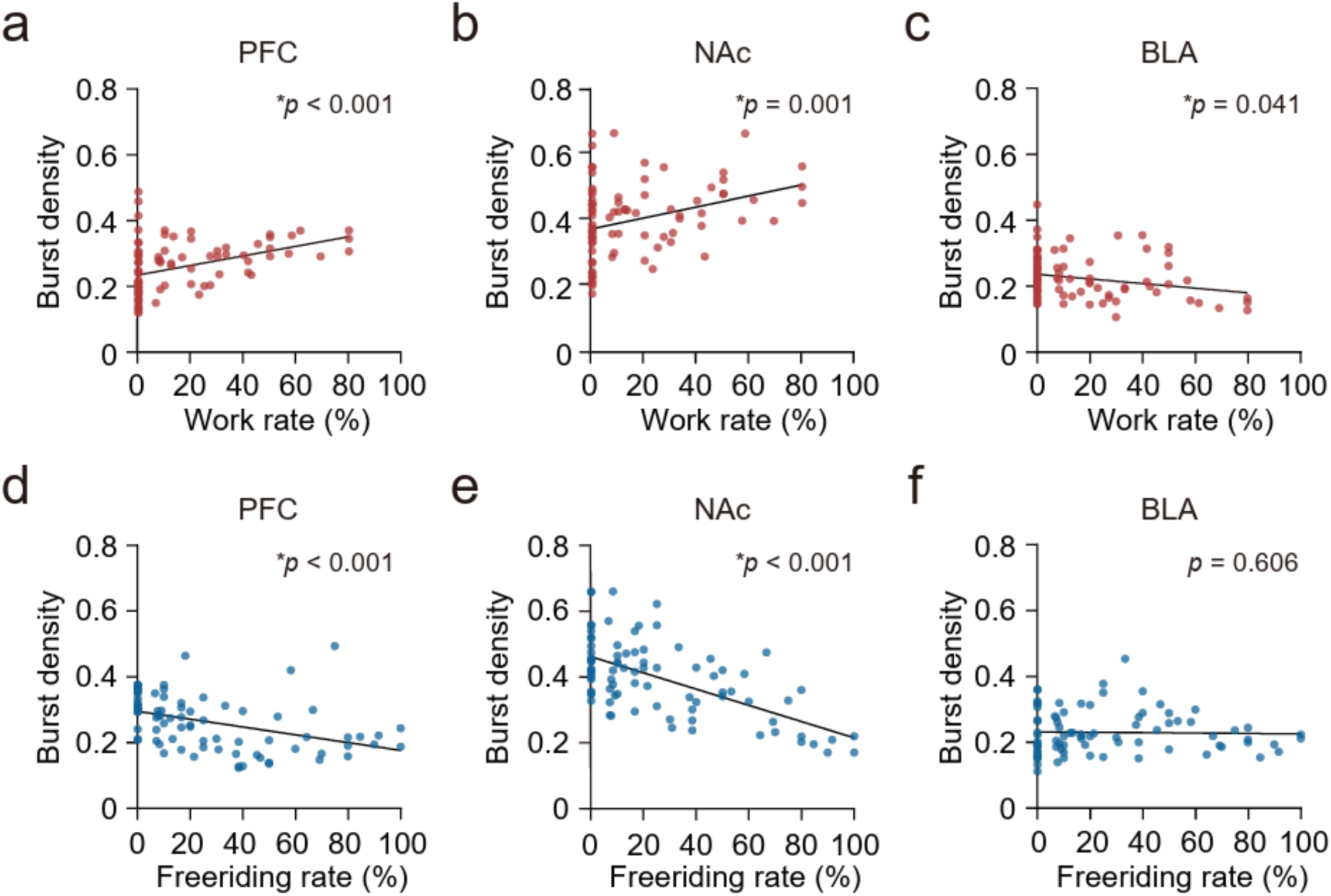
Correlation between gamma burst density and behavioral engagement in work and freeriding. As in Figure. 4b-d, Positive correlations between gamma burst density and work rate were found in **a**, PFC (slope = 0.00146, R² = 0.1608, *p* < 0.001) and **b**, NAc (slope = 0.00168, R² = 0.1064, *p* = 0.002), but a negative correlation was found in the **c**, BLA (slope = –0.0007, R² = 0.05607, *p* = 0.03). As in Figure. 4e-g, Negative correlations were found between gamma burst density and freeriding rate in the **d**, PFC (slope = –0.00118, R² = 0.1761, *p* < 0.001) and **e**, NAc (slope = – 0.00246, R² = 0.3814, *p* < 0.001), but no significant relationship in the **f**, BLA (slope = – 0.00006, R² = 0.00069, *p* = 0.81265). Each dot represents data from te daily average values of each individual (*n* = 84). The lines represent linear regression trends. *p* < 0.05 indicates statistical significance.

**Supplementary Figure. 8.**
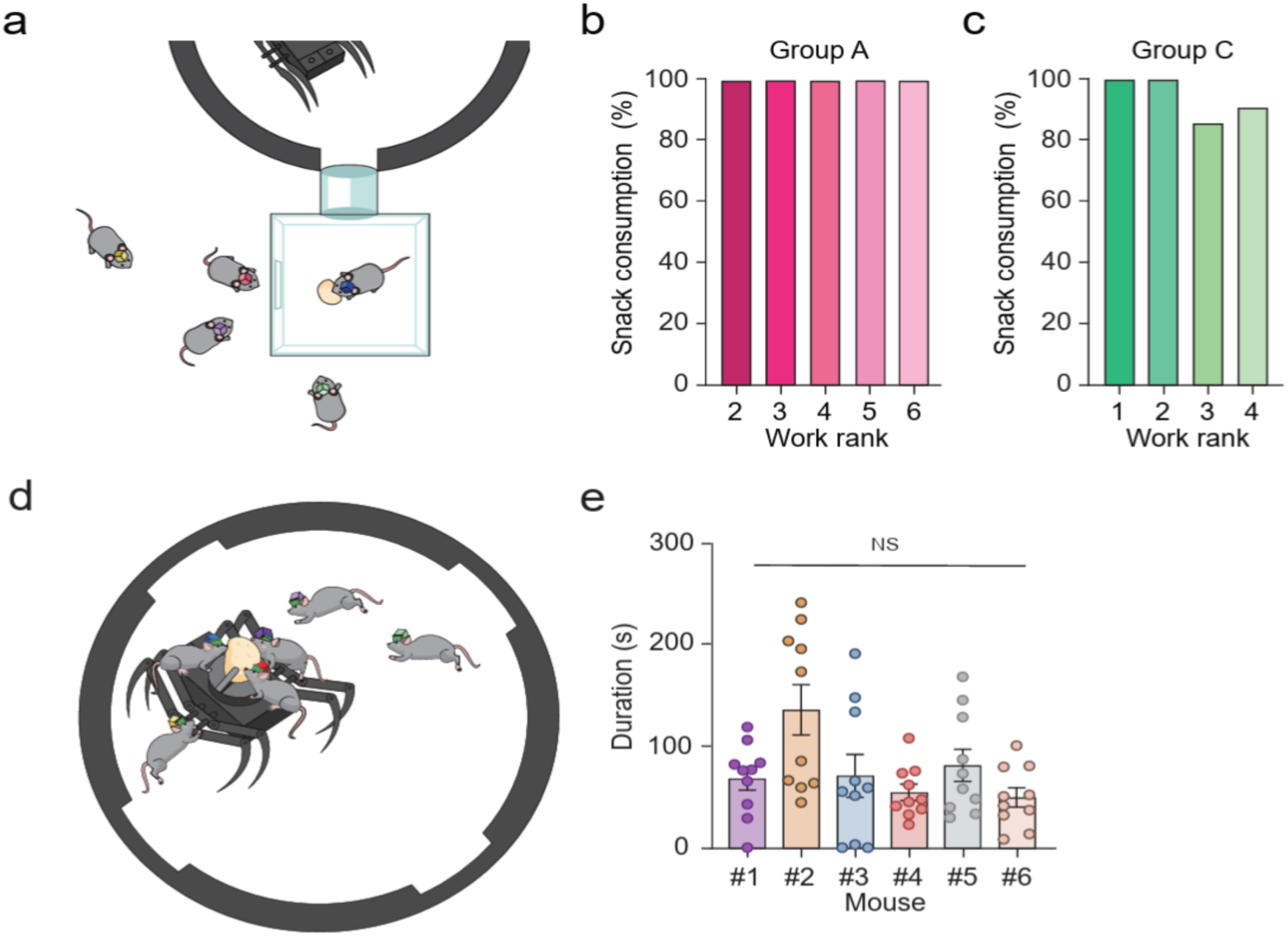
Behavioral motivation of working behavior. To verify whether the motivation behind working behavior is prosocial, we conducted a modified foraging experiment (methods). **a**, Schematic illustration of the prosociality test, featuring an “eating chamber” that allows mice to choose between eating alone or leaving it to share the food. Percentage of snacks consumed inside the box across work ranks showed working behavior is not driven by prosocial motives in **b**, Group A and in **c**, Group C. Next, we examined whether mice independently attempt to acquire and successfully consume food when a worker fails to obtain and retrieve it. **d**, Schematic illustration of the fixed-snack foraging test, where a clip securing the snack, requiring mice to stay on the robot to eat. **e**, There were no significant differences in the time spent consuming the snack on the spider robot between individuals in Group A (*n =* 6 mice, 10 trials per mouse; χ²(5, 60) = 8.662, *p* = 0.123, Kruskal-Wallis test). Each dot represents a single trial, and bars indicate mean ± s.e.m. NS indicates no statistically significant difference.

**Supplementary Figure. 9.**
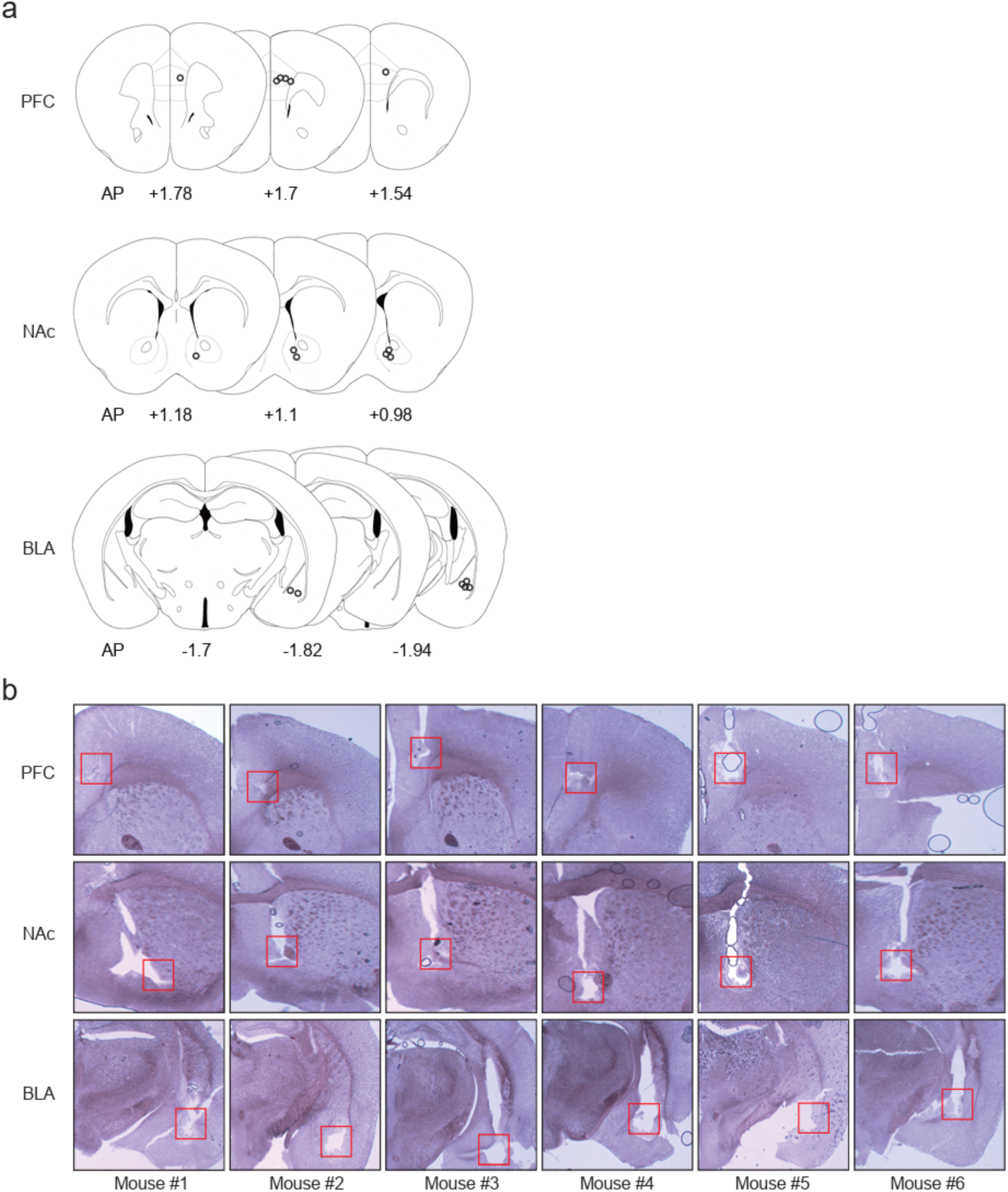
Histological verification of electrode implantation. Electrode placements were confirmed in the PFC, NAc, and BLA through histological verification and brain atlas mapping. All animals included in the study had accurately implanted electrodes, with precise targeting to the intended brain regions. **a**, Coronal sections of brain atlas plates showing the locations of the electrode tips (circles): left PFC (top), NAc (middle) and BLA (right). Each dot represents the position for an individual mouse. **b**, Histological verification of electrode placements in target brain regions of individual mice. The red square indicates the position of the electrode tip. Only Group A mice are plotted, while all mice included in the experiments had correctly placed electrodes. Atlas images redrawn from ‘The Mouse Brain in Stereotaxic Coordinates, 2nd ed.,’ by G. Paxinos and K. Franklin.

**Supplementary Table 1.**
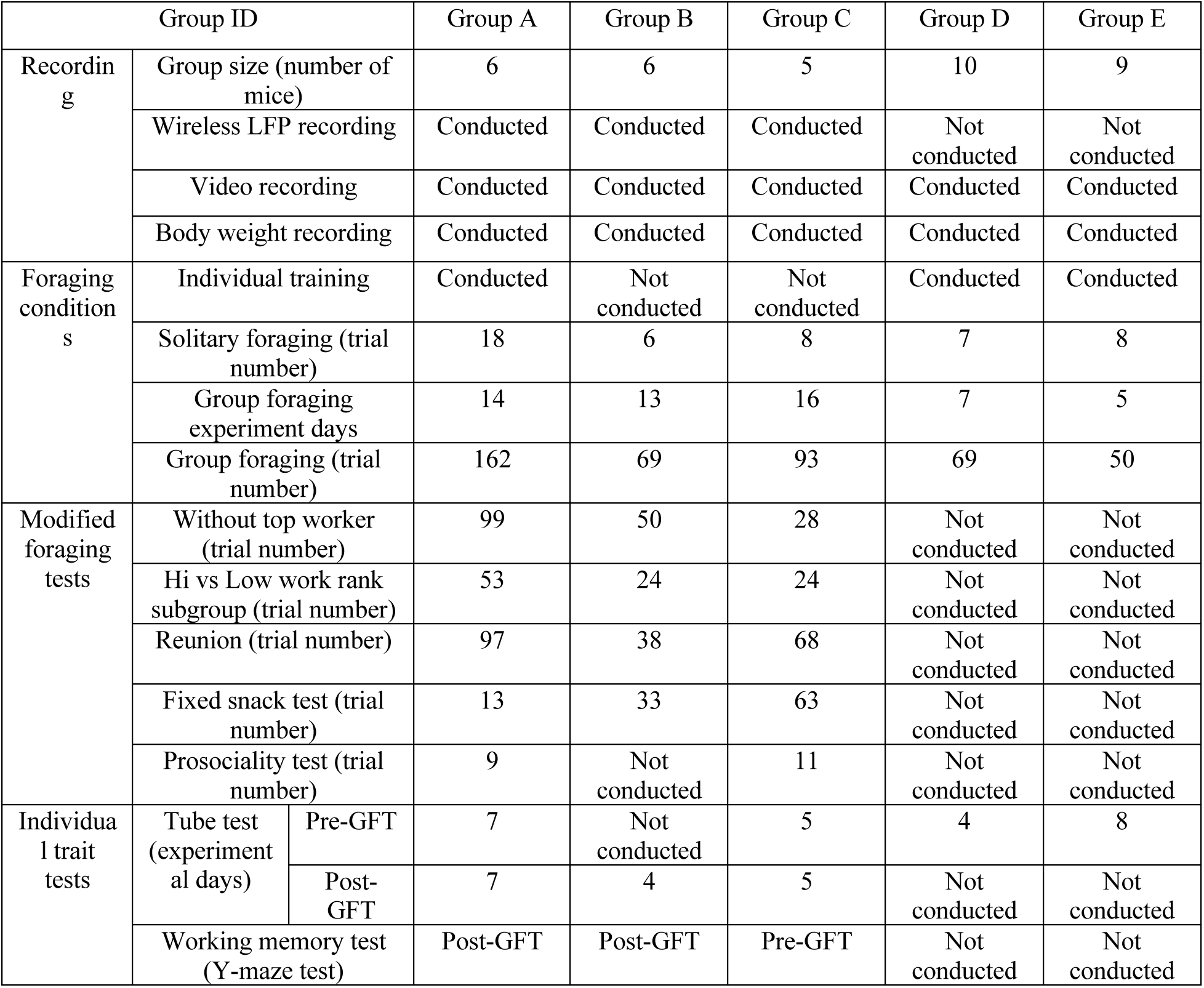
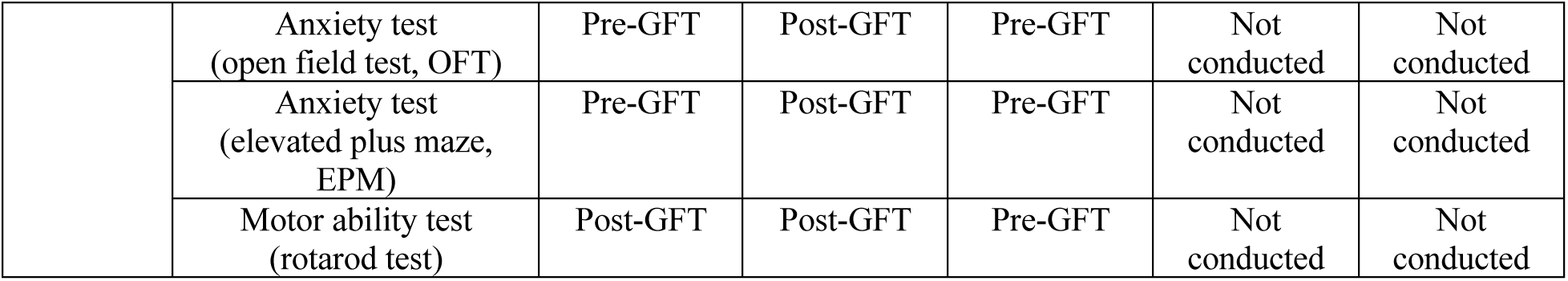
Summary of experimental design, foraging protocols, and behavioral assessments across mouse groups. Experimental setup and protocol summary for groups of mice in the study. Groups A–E represent distinct experimental cohorts with varying group sizes and experimental conditions. The setup includes parameters for wireless LFP recording, video monitoring, and body weight tracking. Foraging experiments consisted of individual training, solitary and group foraging sessions, with additional modified tasks like exclusion of top workers, rank-based subgroup trials, and reunion tests. Individual behavioral traits were evaluated using multiple assays, including the tube test, Y-maze, open field test, elevated plus maze, and rotarod test. Trials and experimental days are indicated for each group where applicable. “Pre-GFT” or “Post-GFT” denote the trials conducted prior or after the whole group foraging task (GFT) sessions, respectively.

| Group ID |  |  | Group A | Group B | Group C | Group D | Group E |
| --- | --- | --- | --- | --- | --- | --- | --- |
| Recording | Group size (number of mice) |  | 6 | 6 | 5 | 10 | 9 |
|  | Wireless LFP recording |  | Conducted | Conducted | Conducted | Not conducted | Not conducted |
|  | Video recording |  | Conducted | Conducted | Conducted | Conducted | Conducted |
|  | Body weight recording |  | Conducted | Conducted | Conducted | Conducted | Conducted |
| Foraging conditions | Individual training |  | Conducted | Not conducted | Not conducted | Conducted | Conducted |
|  | Solitary foraging (trial number) |  | 18 | 6 | 8 | 7 | 8 |
|  | Group foraging experiment days |  | 14 | 13 | 16 | 7 | 5 |
|  | Group foraging (trial number) |  | 162 | 69 | 93 | 69 | 50 |
| Modified foraging tests | Without top worker (trial number) |  | 99 | 50 | 28 | Not conducted | Not conducted |
|  | Hi vs Low work rank subgroup (trial number) |  | 53 | 24 | 24 | Not conducted | Not conducted |
|  | Reunion (trial number) |  | 97 | 38 | 68 | Not conducted | Not conducted |
|  | Fixed snack test (trial number) |  | 13 | 33 | 63 | Not conducted | Not conducted |
|  | Prosociality test (trial number) |  | 9 | Not conducted | 11 | Not conducted | Not conducted |
| Individual trait tests | Tube test (experimental days) | Pre-GFT | 7 | Not conducted | 5 | 4 | 8 |
|  |  | Post-GFT | 7 | 4 | 5 | Not conducted | Not conducted |
|  | Working memory test (Y-maze test) |  | Post-GFT | Post-GFT | Pre-GFT | Not conducted | Not conducted |

|  |  |  |  |  |  |  |
| --- | --- | --- | --- | --- | --- | --- |
|  | Anxiety test<br>(open field test, OFT) | Pre-GFT | Post-GFT | Pre-GFT | Not<br>conducted | Not<br>conducted |
|  | Anxiety test<br>(elevated plus maze,<br>EPM) | Pre-GFT | Post-GFT | Pre-GFT | Not<br>conducted | Not<br>conducted |
|  | Motor ability test<br>(rotarod test) | Post-GFT | Post-GFT | Pre-GFT | Not<br>conducted | Not<br>conducted |

**Supplementary Table 2.** Summary of individual trait scores in relation to work rate. This table presents the relationship between work rank, work rate, and individual trait scores for mice across the groups. Work rank categories (Worker, Participant, Freerider) are defined based on the percentage of working trials (%). Individual trait assessments include social dominance (ELO score and normalized score), height anxiety (elevated plus maze: time spent in the open arm and normalized score), motor coordination (rotarod: drop latency), open space anxiety (open field test: time spent in the center and normalized score), and working memory (Y-maze: percentage of arm alternation). “N/A“ indicates data unavailable due to mortality or other factors.

| Group | Work rank | Dominant role across trials | Percentage of working trials (%) | Tube test (Social dominance) |  | Elevated plus maze (Height anxiety) |  | Rotor rod (Motor coordination) |  | Open field test (Open space anxiety) |  | Y-maze (Working memory) |
| --- | --- | --- | --- | --- | --- | --- | --- | --- | --- | --- | --- | --- |
|  |  |  |  | ELO score (average) | Norm. score (%) | Time spent in open arm (sec) | Norm. score (%) | Drop latency (sec) | Norm. score (%) | Time spent in center (sec) | Norm. score (%) | Arm alternation (%) |
| A | 1 | Top worker | 46.30 | 2.86 | 47.62 | 145.57 | 24.26 | N/A | N/A | 228.68 | 19.06 | N/A |
|  | 2 | Participant | 22.84 | 4.57 | 76.19 | 178.84 | 29.81 | Max (300) | 100.00 | 163.49 | 13.62 | 48.28 |
|  | 3 | Participant | 17.90 | 4.43 | 73.81 | 12.39 | 2.07 | 284.30 | 94.77 | 152.79 | 12.73 | 56.67 |
|  | 4 | Freerider | 11.73 | 4.14 | 69.05 | 110.61 | 18.43 | 290.08 | 96.69 | 145.37 | 12.11 | 41.67 |
|  | 5 | Freerider | 0.62 | 1.43 | 23.81 | 88.56 | 14.76 | 287.81 | 95.94 | 98.13 | 8.18 | 65.12 |
|  | 6 | Freerider | 0.62 | 2.00 | 33.33 | 86.53 | 14.42 | 242.68 | 80.89 | 84.07 | 7.01 | 44.44 |
| B | 1 | Top worker | 73.13 | 2.75 | 45.83 | 113.31 | 18.88 | 157.20 | 52.40 | 113.98 | 9.50 | 57.89 |
|  | 2 | Participant | 22.39 | 4.75 | 79.17 | 153.87 | 25.65 | 198.94 | 66.31 | 29.92 | 2.49 | 63.64 |
|  | 3 | Freerider | 4.48 | 3.50 | 58.33 | 0.00 | 0.00 | 130.36 | 43.45 | 301.12 | 25.09 | 53.85 |
|  | 4 | Freerider | 0.00 | 1.00 | 16.67 | 114.60 | 19.10 | 195.33 | 65.11 | 81.83 | 6.82 | 57.14 |
|  | 5 | Freerider | 0.00 | 5.75 | 95.83 | 30.97 | 5.16 | 151.55 | 50.52 | 0.00 | 0.00 | Nan |
|  | 6 | Freerider | 0.00 | 2.75 | 45.83 | 95.52 | 15.92 | 169.08 | 56.36 | 138.67 | 11.56 | 52.94 |
| C | 1 | Top worker | 100.00 | 3.00 | 50.00 | 208.53 | 34.75 | 226.91 | 75.64 | 161.64 | 13.47 | 84.62 |
|  | 2 | Freerider | 0.00 | 2.20 | 36.67 | 144.4<br>8 | 24.08 | 254.4<br>8 | 84.83 | 214.5<br>5 | 17.88 | 52.00 |
|  | 3 | Freerider | 0.00 | 5.60 | 93.33 | 10.39 | 1.73 | 196.3<br>8 | 65.46 | 147.6<br>8 | 12.31 | 70.27 |
|  | 4 | Freerider | 0.00 | 2.20 | 36.67 | 64.85 | 10.81 | 160.3<br>0 | 53.43 | 176.9<br>6 | 14.75 | 54.84 |
|  | 5 | Freerider | 0.00 | 3.80 | 63.33 | 128.2<br>9 | 21.38 | 182.0<br>8 | 60.69 | 45.91 | 3.83 | 77.27 |
| D | 1 | Top<br>worker | 80.00 | 4.50 | 45.00 |  |  |  |  |  |  |  |
|  | 2 | Participa<br>nt | 18.57 | 3.75 | 37.50 |  |  |  |  |  |  |  |
|  | 3 | Freerider | 1.43 | 1.00 | 10.00 |  |  |  |  |  |  |  |
|  | 4 | Freerider | 0.00 | 4.75 | 47.50 |  |  |  |  |  |  |  |
|  | 5 | Freerider | 0.00 | 8.75 | 87.50 |  |  |  |  |  |  |  |
|  | 6 | Freerider | 0.00 | 7.25 | 72.50 |  |  |  |  |  |  |  |
|  | 7 | Freerider | 0.00 | 7.00 | 70.00 |  |  |  |  |  |  |  |
|  | 8 | Freerider | 0.00 | 7.25 | 72.50 |  |  |  |  |  |  |  |
|  | 9 | Freerider | 0.00 | 7.00 | 70.00 |  |  |  |  |  |  |  |
|  | 10 | Freerider | 0.00 | 2.00 | 20.00 |  |  |  |  |  |  |  |
| E | 1 | Top<br>worker | 62.00 | 8.13 | 81.25 |  |  |  |  |  |  |  |
|  | 2 | Participa<br>nt | 16.00 | 3.75 | 37.50 |  |  |  |  |  |  |  |
|  | 3 | Freerider | 10.00 | 1.75 | 17.50 |  |  |  |  |  |  |  |
|  | 4 | Freerider | 6.00 | 3.50 | 35.00 |  |  |  |  |  |  |  |
|  | 5 | Freerider | 4.00 | 2.13 | 21.25 |  |  |  |  |  |  |  |
|  | 6 | Freerider | 2.00 | 8.88 | 88.75 |  |  |  |  |  |  |  |
|  | 7 | Freerider | 0.00 | 7.00 | 70.00 |  |  |  |  |  |  |  |
|  | 8 | Freerider | 0.00 | 5.13 | 51.25 |  |  |  |  |  |  |  |
|  | 9 | Freerider | 0.00 | 6.50 | 65.00 |  |  |  |  |  |  |  |

**Supplementary Table 3.**
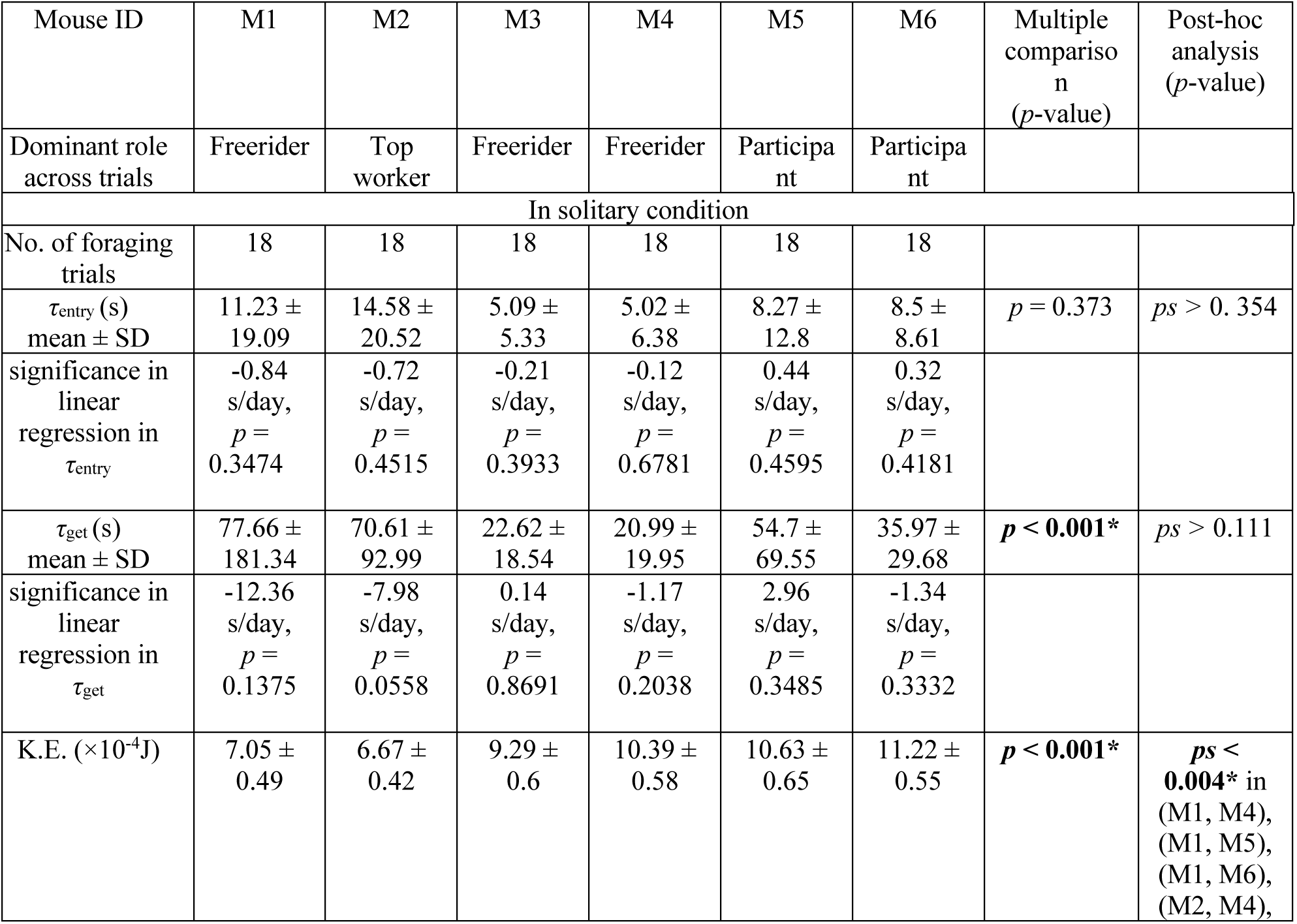
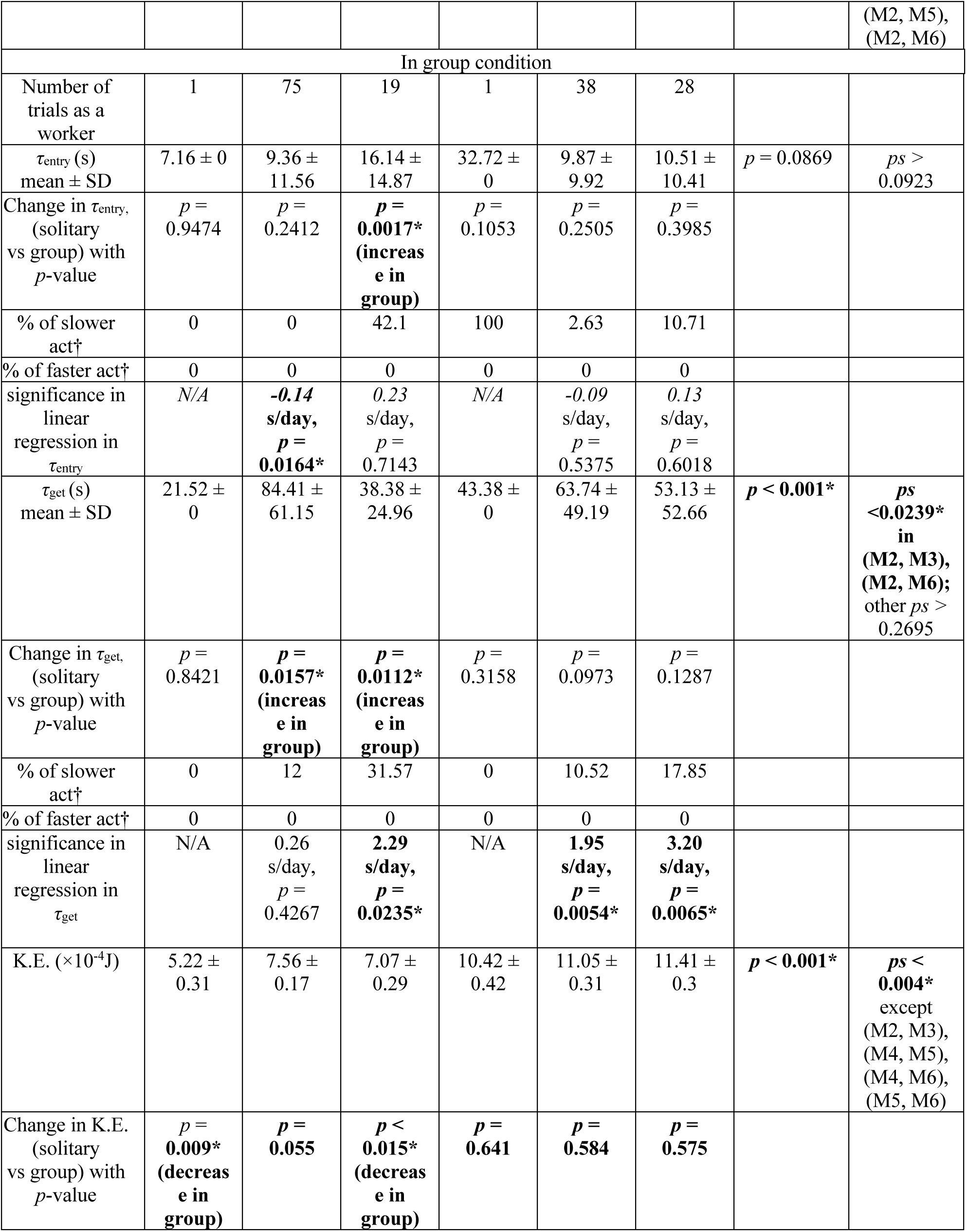
Foraging time parameters and kinetic energy of individual mice across solitary and group conditions. This table summarizes the foraging time parameters and kinetic energy (K.E.) of individual mice acting as workers in each trial under solitary and group conditions. The parameters include *τ*_entry_, which is the entry latency or the time taken to enter the robot zone after the snack is placed on the robot, and *τ*_get_, which is the get duration or the time taken to retrieve the snack after entering the robot zone. K.E. represents the kinetic energy 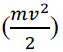, where *m* is the mouse’s weight and *v* is the instantaneous speed extracted from the trajectory. The significance of slower or faster activity † indicates the ratio of values exceeding the confidence interval of the solitary condition. *p*-values are presented for statistical comparisons, with values less than 0.05 considered significant and marked with an asterisk (*). Multiple comparisons were performed using Kruskal-Wallis test. N/A denotes not available due to lack of data for fitting linear regression.

**Supplementary Table 4.** Eating latency, duration, and related metrics of individual mice. This table summarizes the eating latency (*τ*_eat_) and eating duration (*T*_eat_) for individual mice during foraging tasks in Group A. *τ*_eat_ represents the time taken to start eating after the worker retrieves the snack from the robot zone, and *T*_eat_ refers to the total eating duration. Statistical analyses include linear regression and comparisons between solitary and group conditions, with *p*-values reported for each parameter (α=0.05). Asterisks (*) indicate statistically significant differences. To account for cases where eating occurred too quickly, food consumption was calculated based on trials where the eating duration exceeded 4 minutes. Additionally, trials in which Mouse 3 and Mouse 4 failed to participate in eating (twice and once, respectively) were excluded from the analysis. In total, 151 trials were included in the calculations.

| | M1 | M2 | M3 | M4 | M5 | M6 | Multiple<br>comparis<br>on<br>($p$-value) | Post-hoc<br>analysis<br>($p$-value) |
| --- | --- | --- | --- | --- | --- | --- | --- | --- |
| Dominant role<br>the mouse<br>played | Freerider | Worker | Freerider | Freerider | Participa<br>nt | Participa<br>nt |  |  |
| $\tau_{\text{eat}}$ (s)<br>mean ± SD | 5.87 ±<br>6.69 | 3,99 ±<br>5.14 | 5.00 ±<br>6.02 | 4.73 ±<br>5.26 | 5.05 ±<br>6.62 | 3.82 ±<br>4.99 | $p =$<br>0.0976 | $ps > 0.0842$ |
| significance in<br>linear<br>regression in<br>$\tau_{\text{eat}}$ | **0.001**<br>s/trial,<br>$p =$<br>**0.0084\*** | **0.0006**<br>s/trial,<br>$p =$<br>**0.049\*** | 0.0004<br>s/trial<br>$p =$<br>0.2554 | **0.0006**<br>s/trial,<br>$p =$<br>**0.0484\*** | **0.0008**<br>s/trial,<br>$p =$<br>**0.0312\*** | 0.0005<br>s/trial,<br>$p =$<br>0.0587 | | |
| $T_{\text{eat}}$ (s)<br>mean ± SD | 136.67 ±<br>30.49 | 140.31 ±<br>33.43 | 140.65 ±<br>29.95 | 139.94 ±<br>31.41 | 141.12 ±<br>32.12 | 145.34 ±<br>32.94 | $p =$<br>0.3733 | $ps > 0.4131$ |
| significance<br>in linear<br>regression in<br>$T_{\text{eat}}$ | **0.34**<br>s/trial,<br>$p <$<br>**0.001\*** | **0.49**<br>s/trial,<br>$p <$<br>**0.001\*** | **0.39 s/**<br>**trial,**<br>$p <$<br>**0.001\*** | **0.39 s/**<br>**trial,**<br>$p <$<br>**0.001\*** | **0.49 s/**<br>**trial,**<br>$p <$<br>**0.001\*** | **0.48 s/**<br>**trial,**<br>$p <$<br>**0.001\*** | | |

**Supplementary Table 5.**
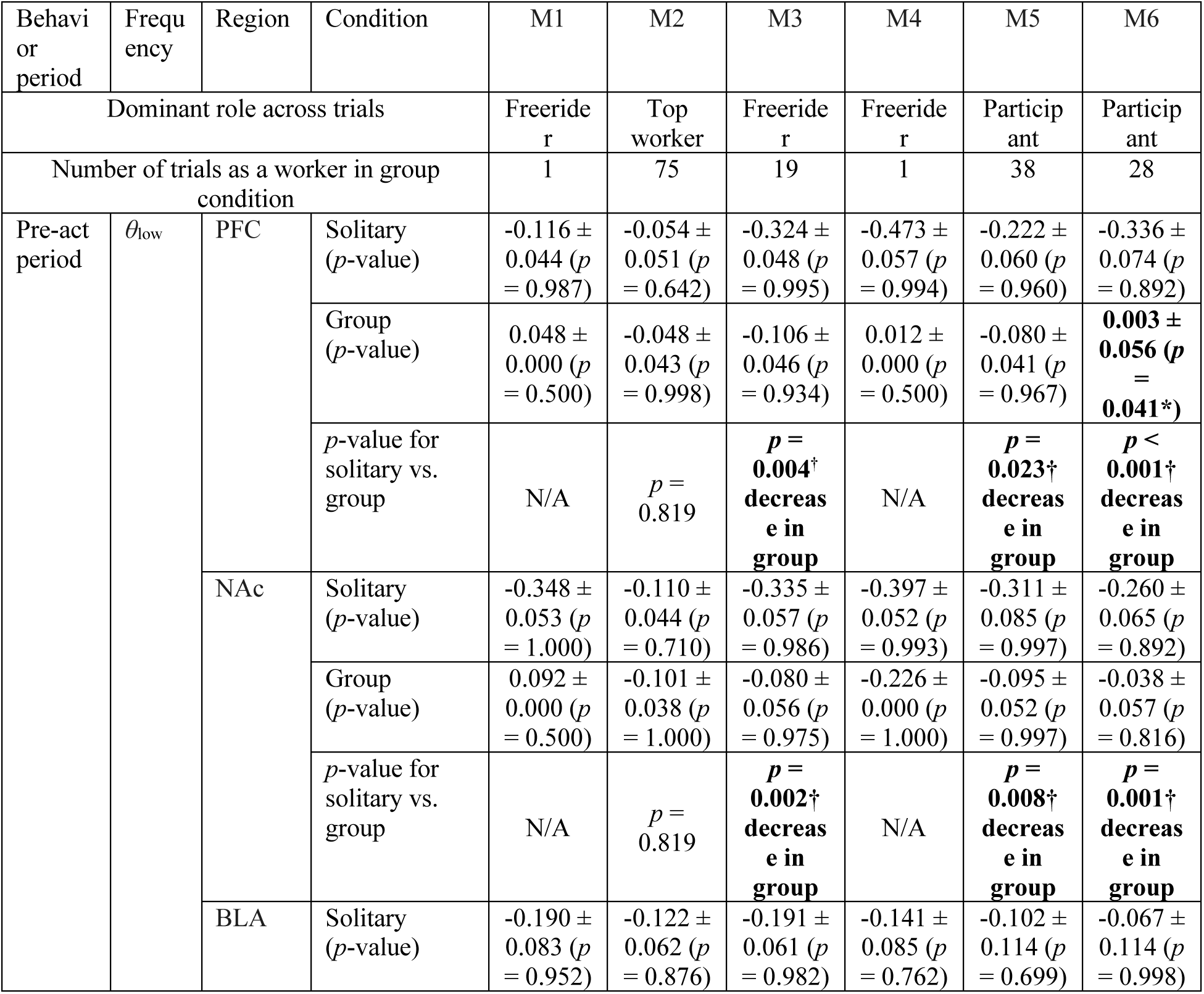
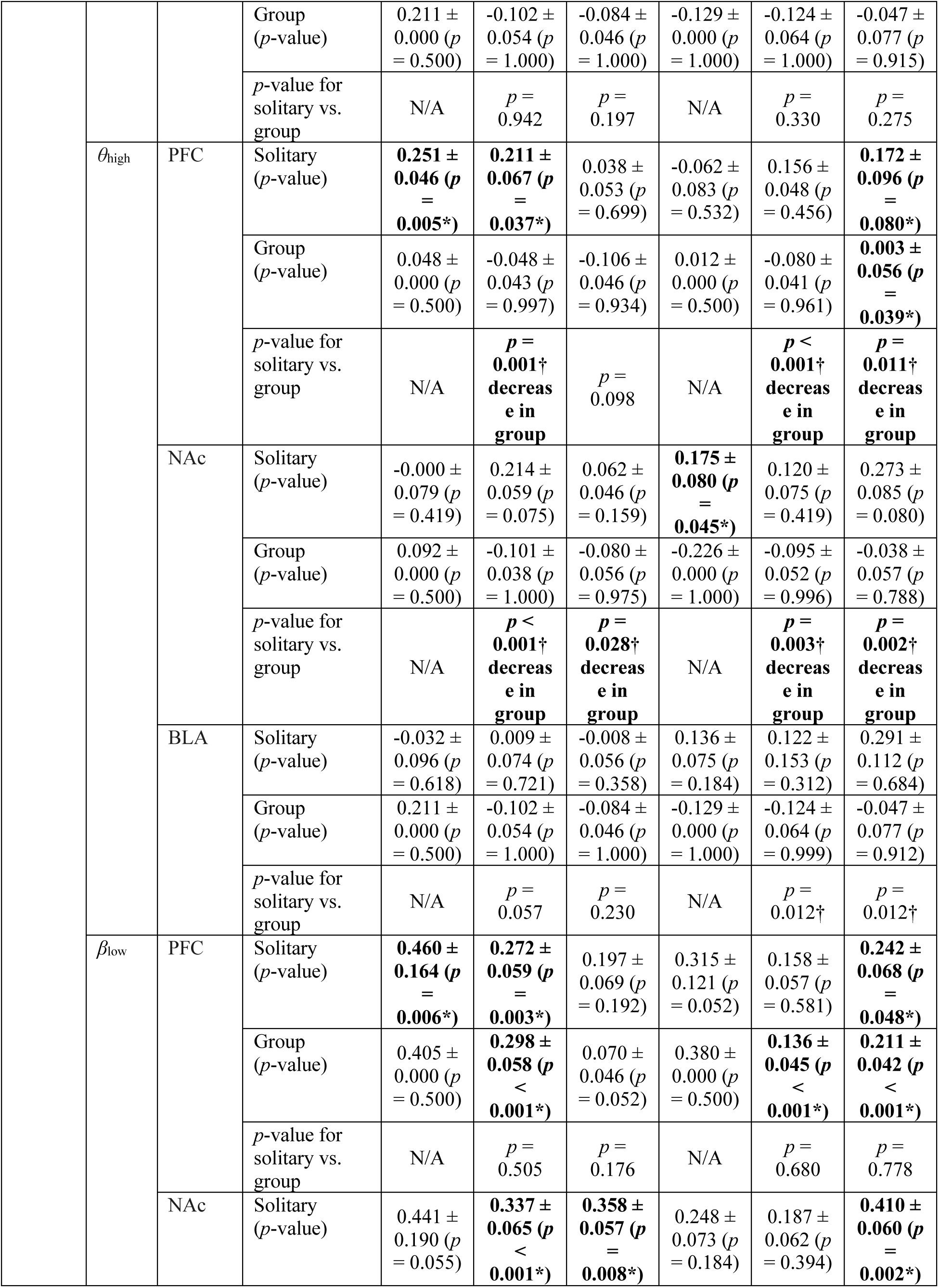
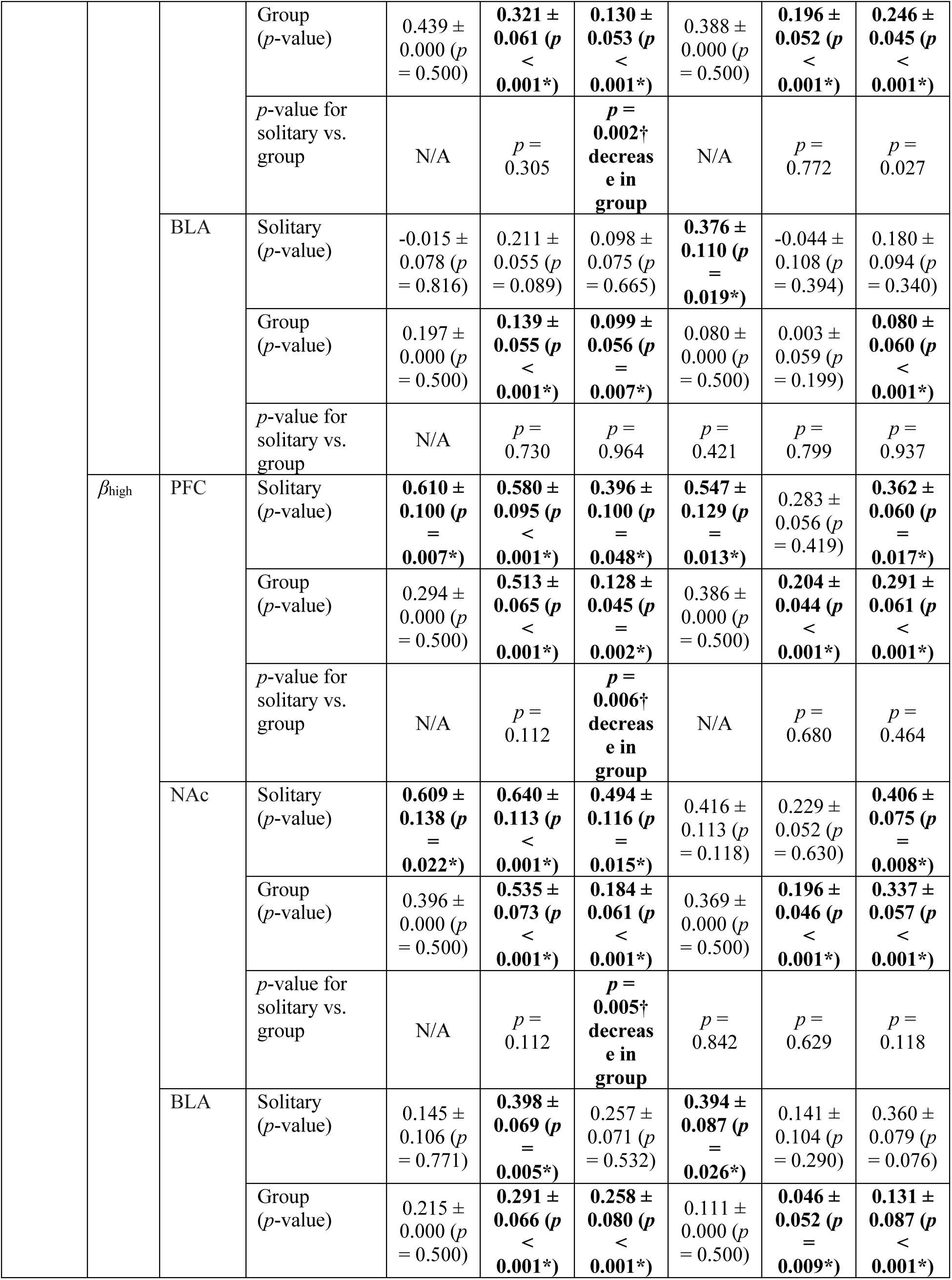
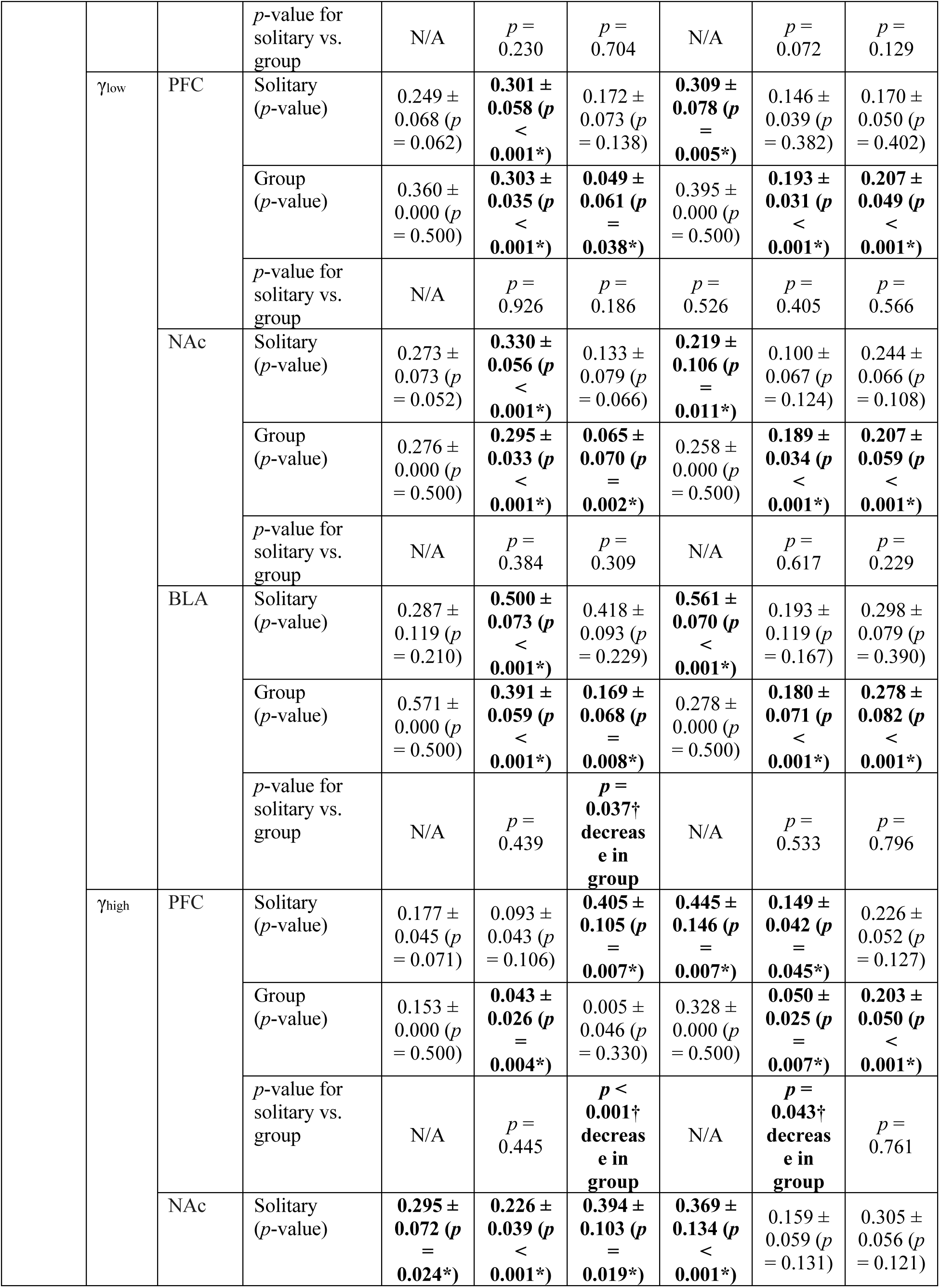
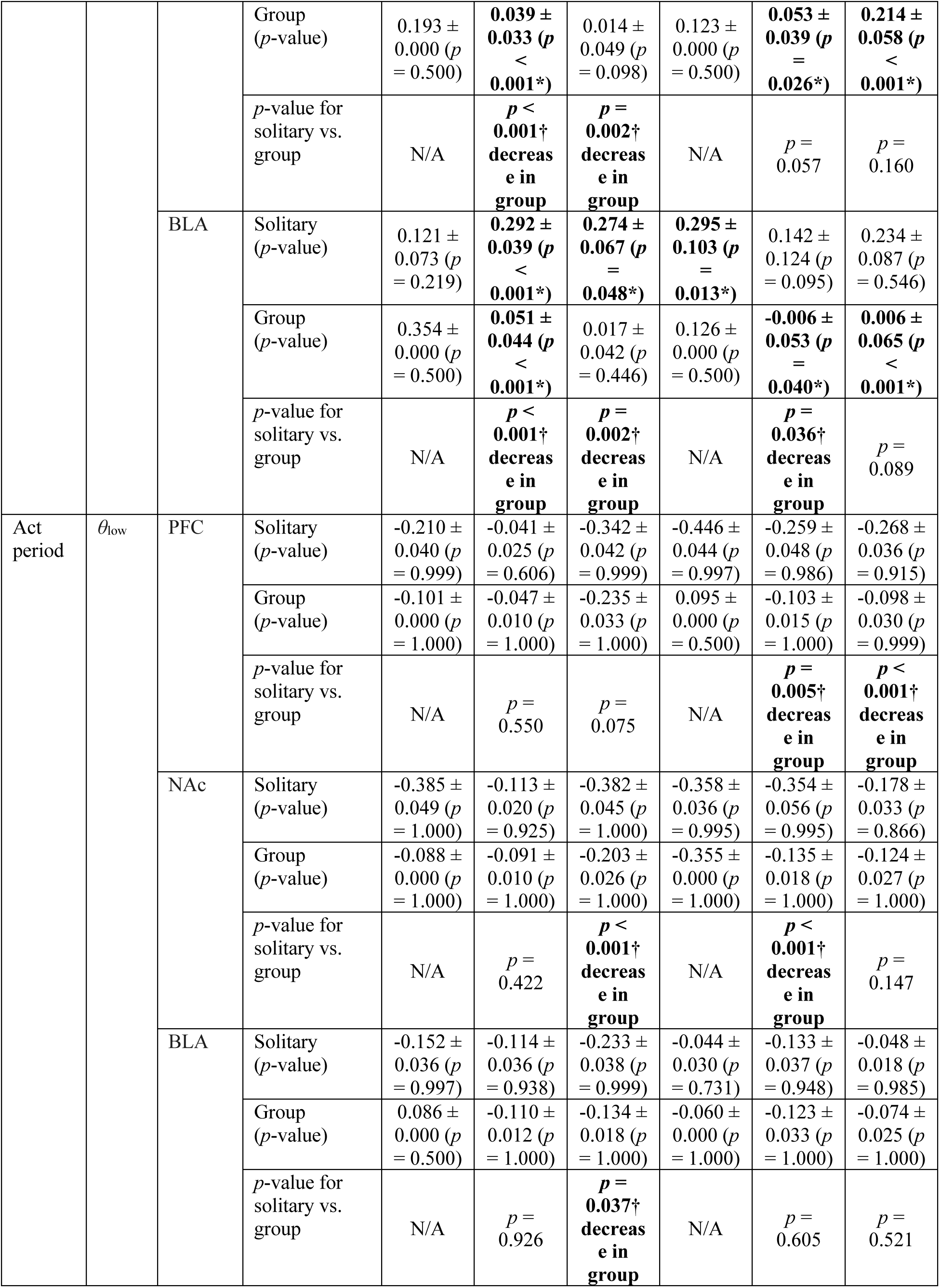
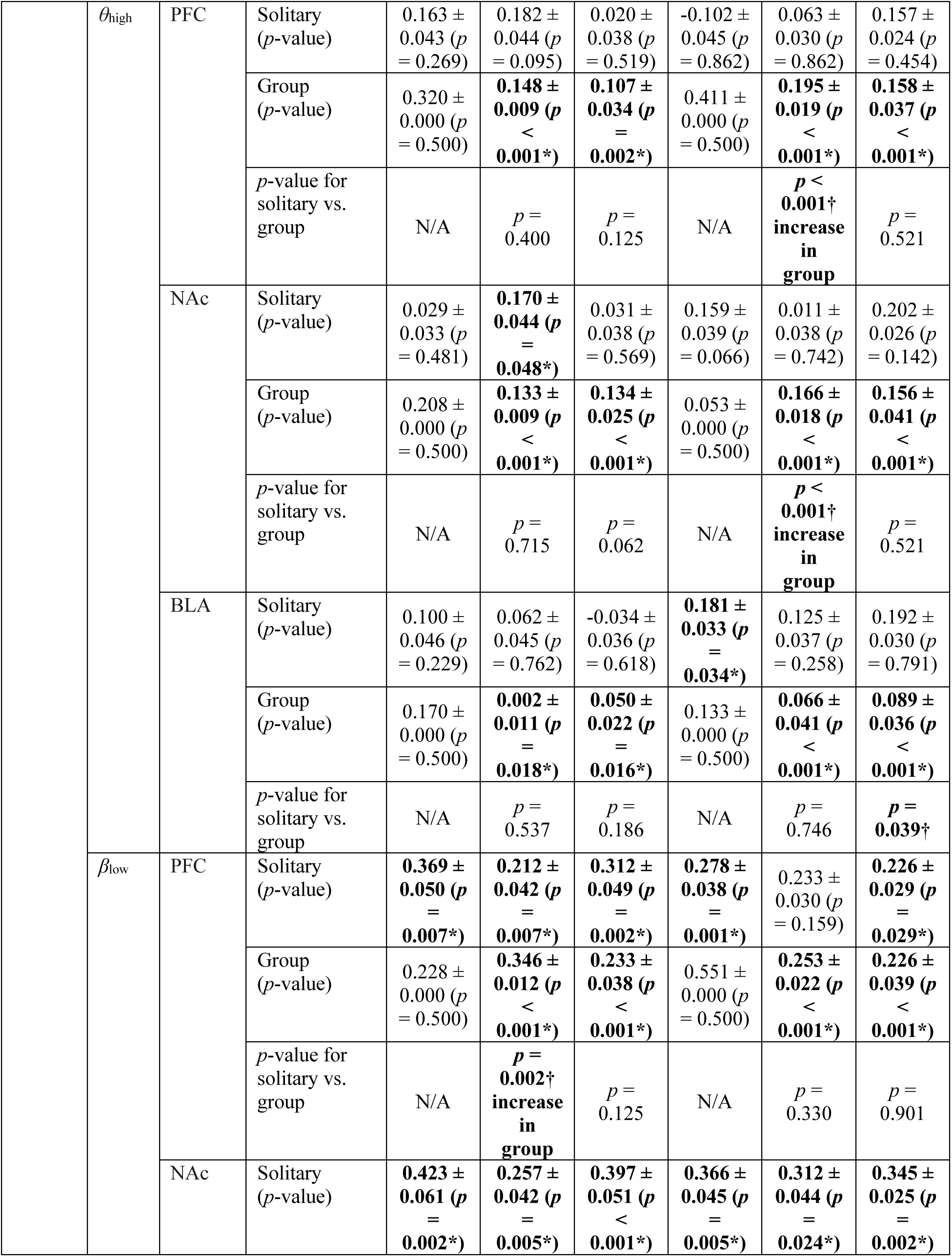
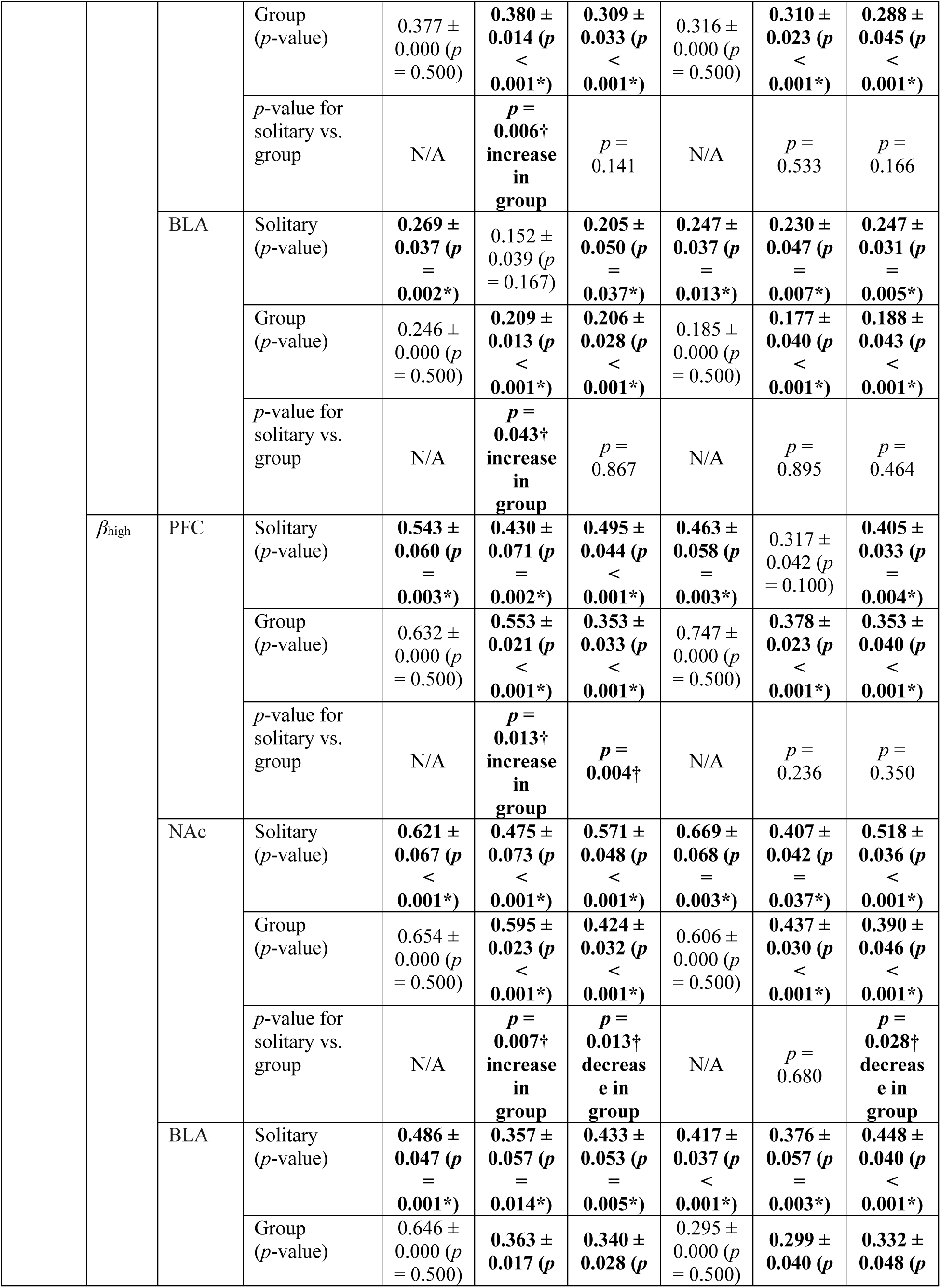
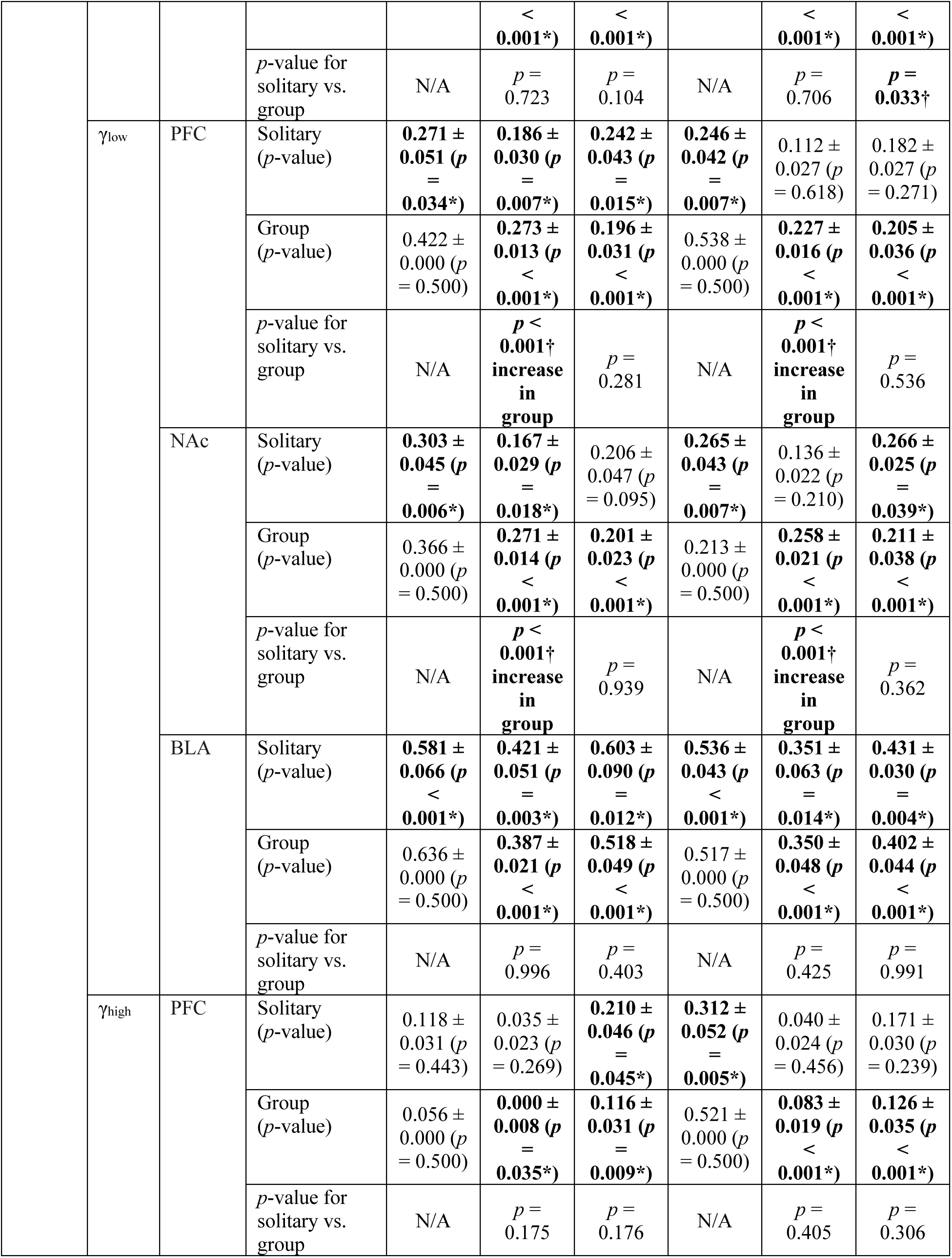
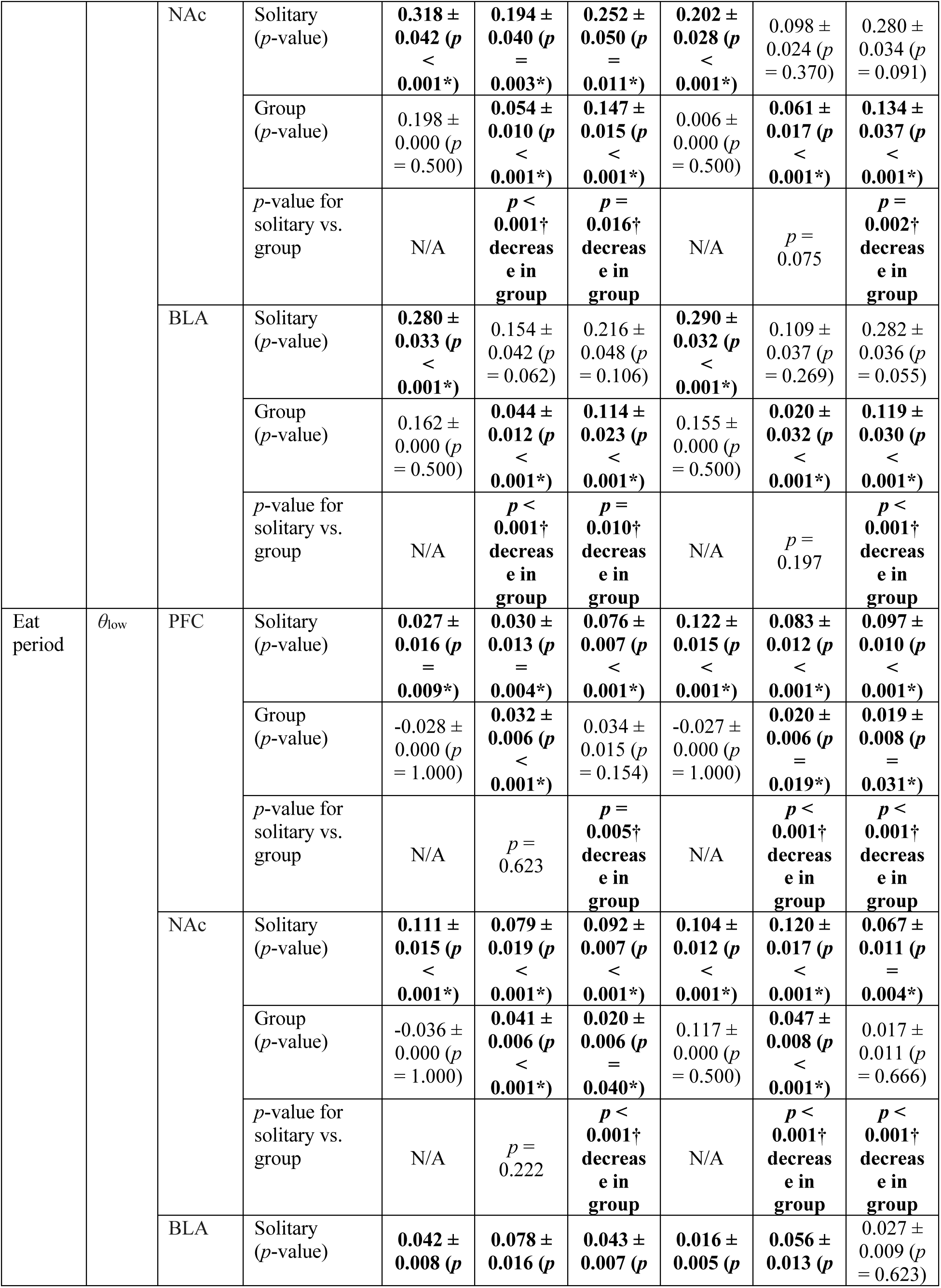
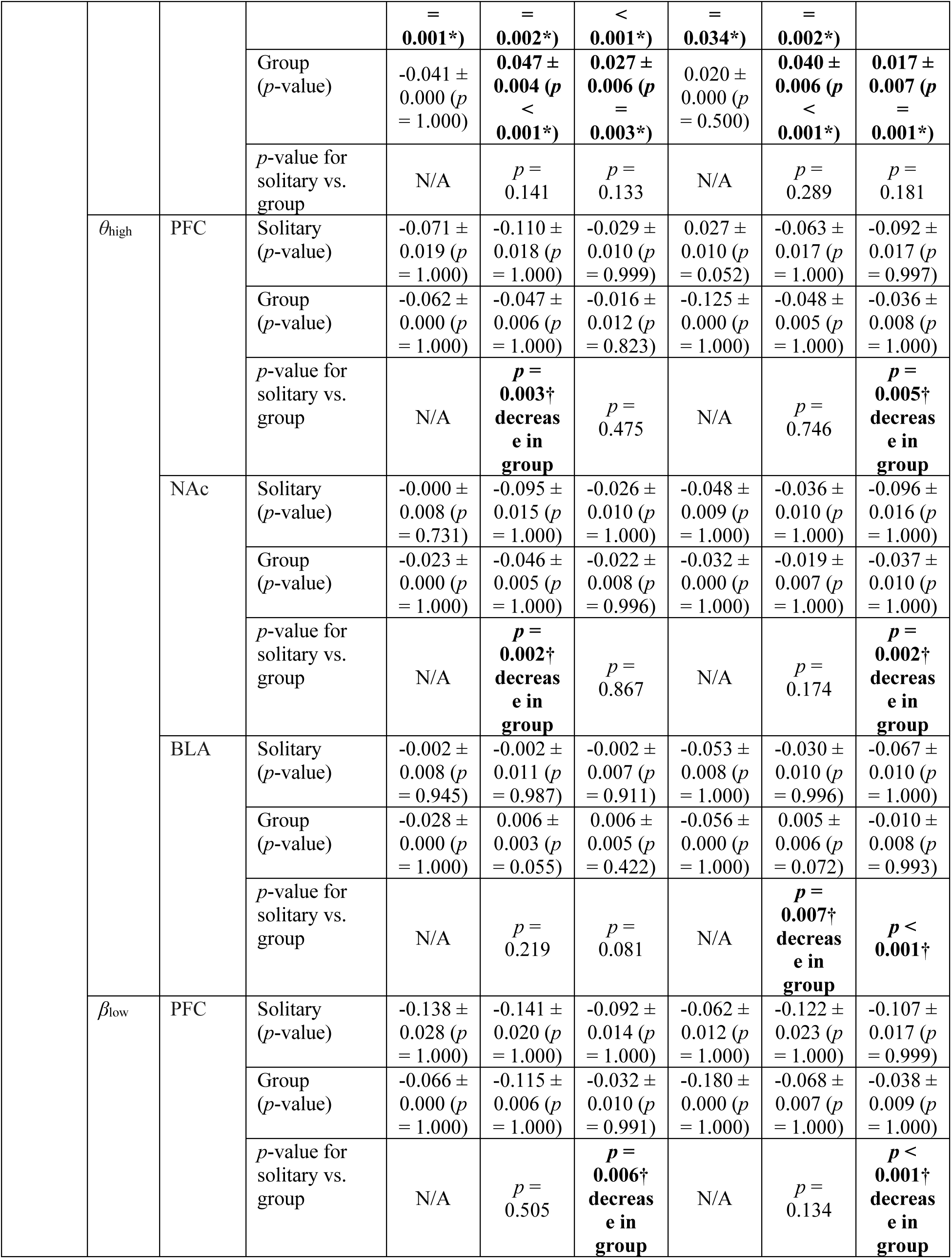
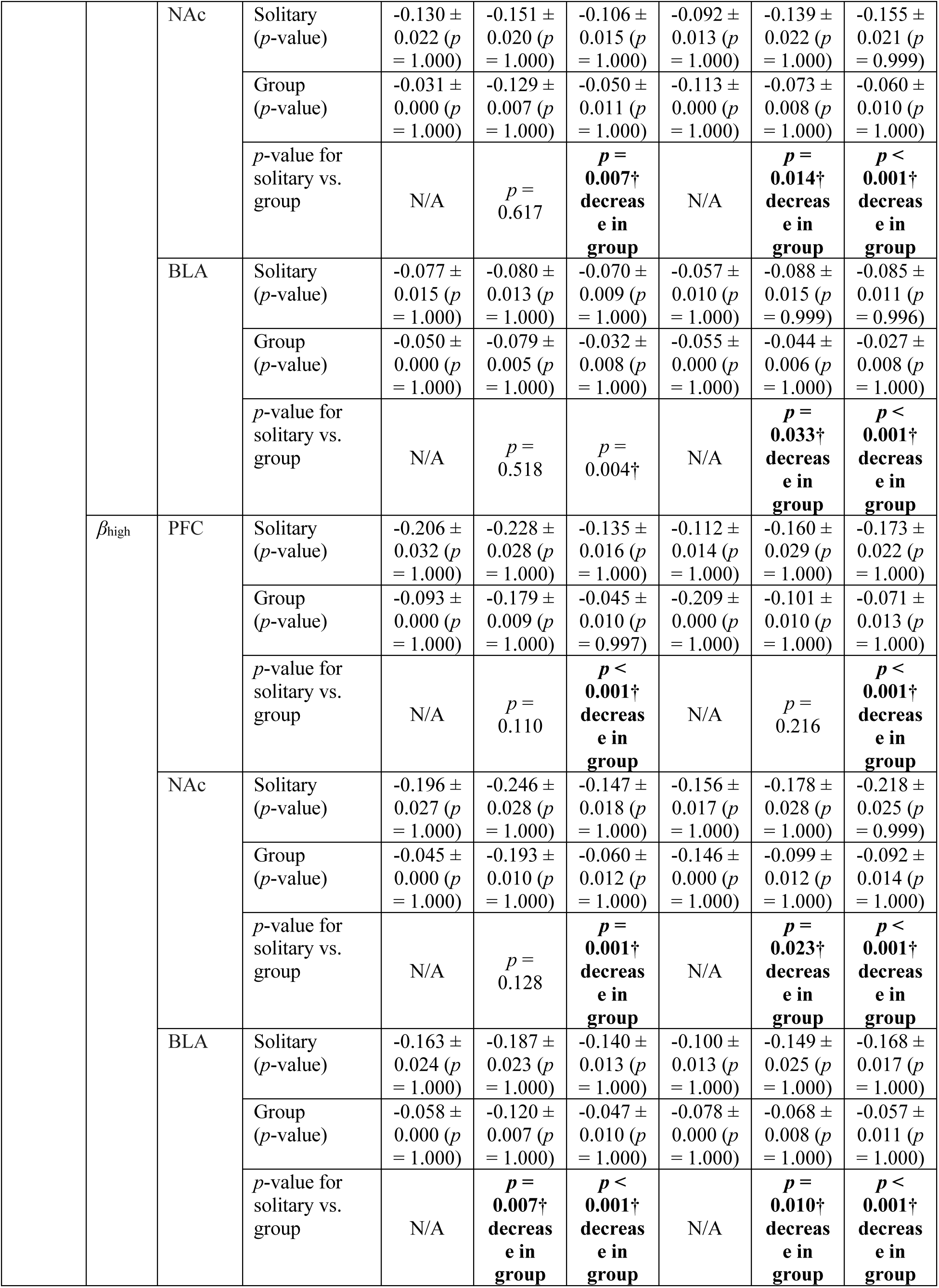
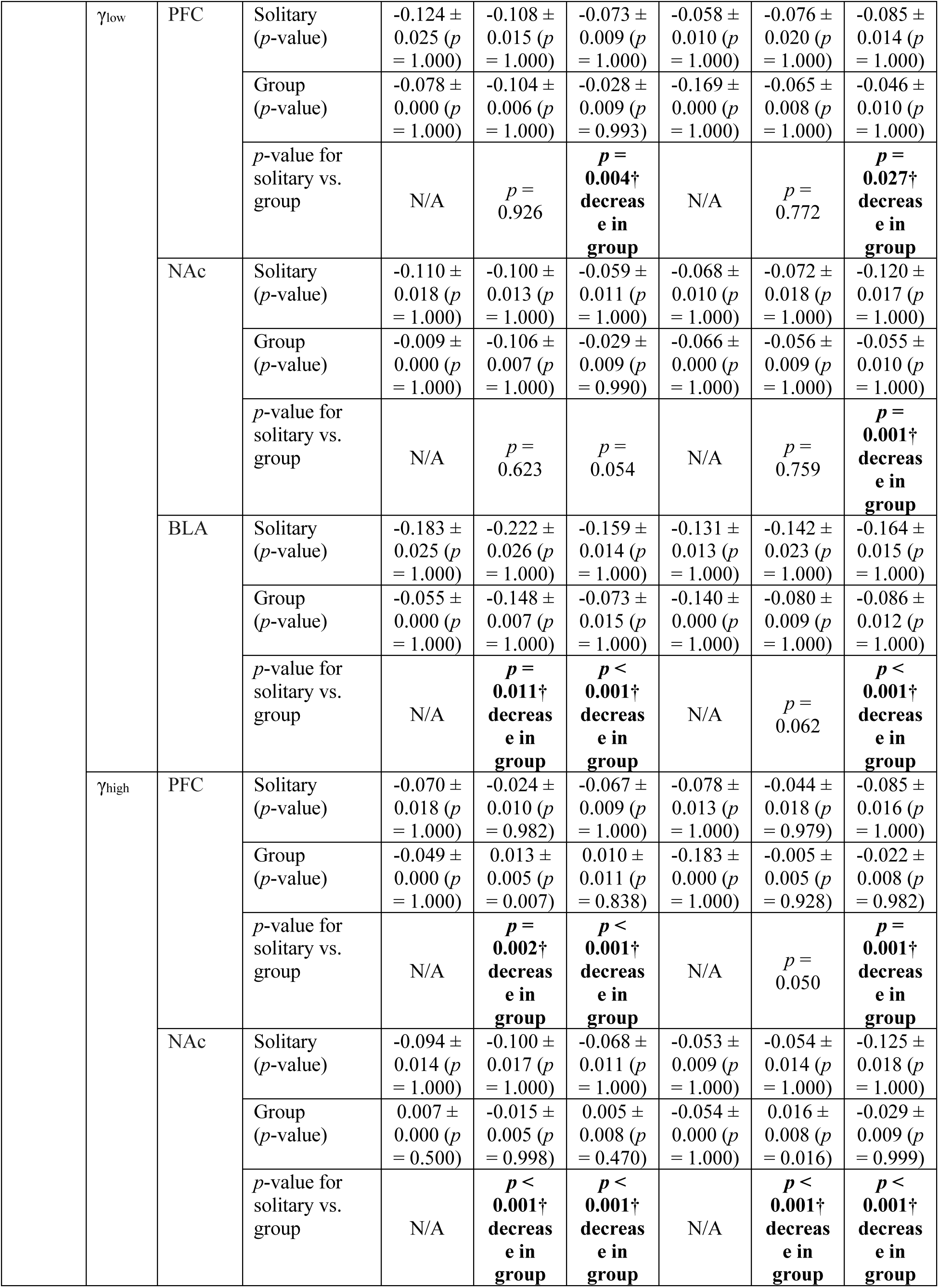
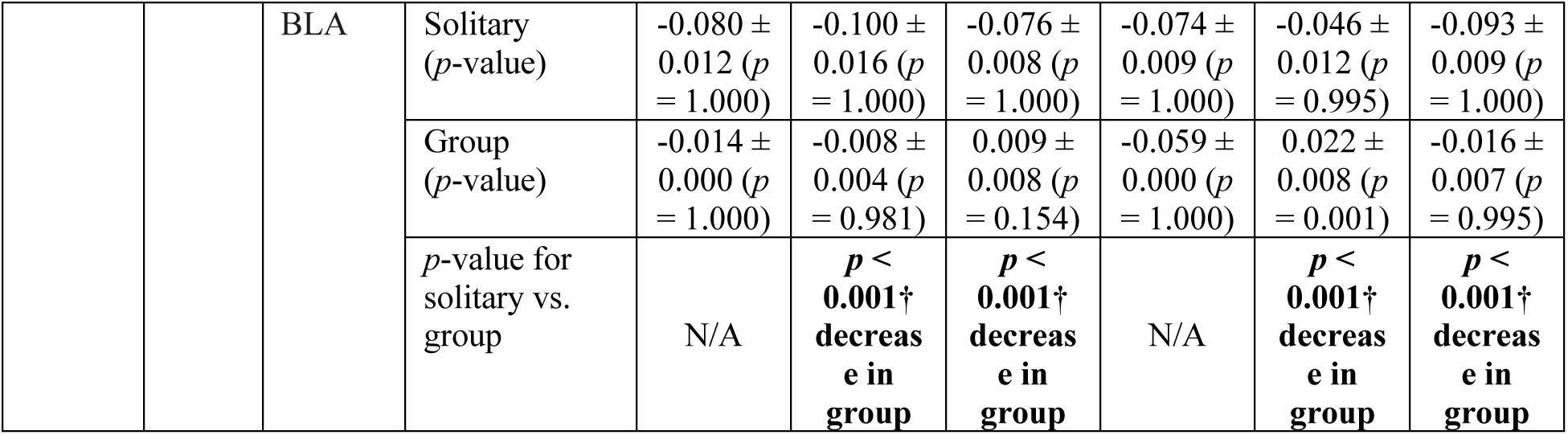
z-scored spectral power differences between solitary and group foraging tasks. This table summarizes the z-scored spectral power differences in local field potentials (LFPs) observed during solitary and group foraging tasks across brain regions, focusing on mice acting as workers. Each value represents the z-score of LFP power changes during distinct behavioral states of the foraging task under solitary (18 trials) and group (162 trials) conditions in Group A. Asterisks (*) indicate statistically significant differences from the baseline (null hypothesis, *α* = 0.05), while daggers † denote significant differences between group and solitary conditions (Mann-Whitney U test, *α* = 0.05). “increase” or “decrease” is noted on the based on the absolute values of the z-score. N/A denotes not available due to lack of data for statistical tests. Frequency bands are defined as follows: *θ*_low_ (4–8 Hz), *θ*_high_ (8–12 Hz), *β*_low_ (18–24 Hz), *β*_high_ (24–32 Hz), *γ*_low_ (35–50 Hz), and *γ*_high_ (70–90 Hz).

**Supplementary Table 6.**
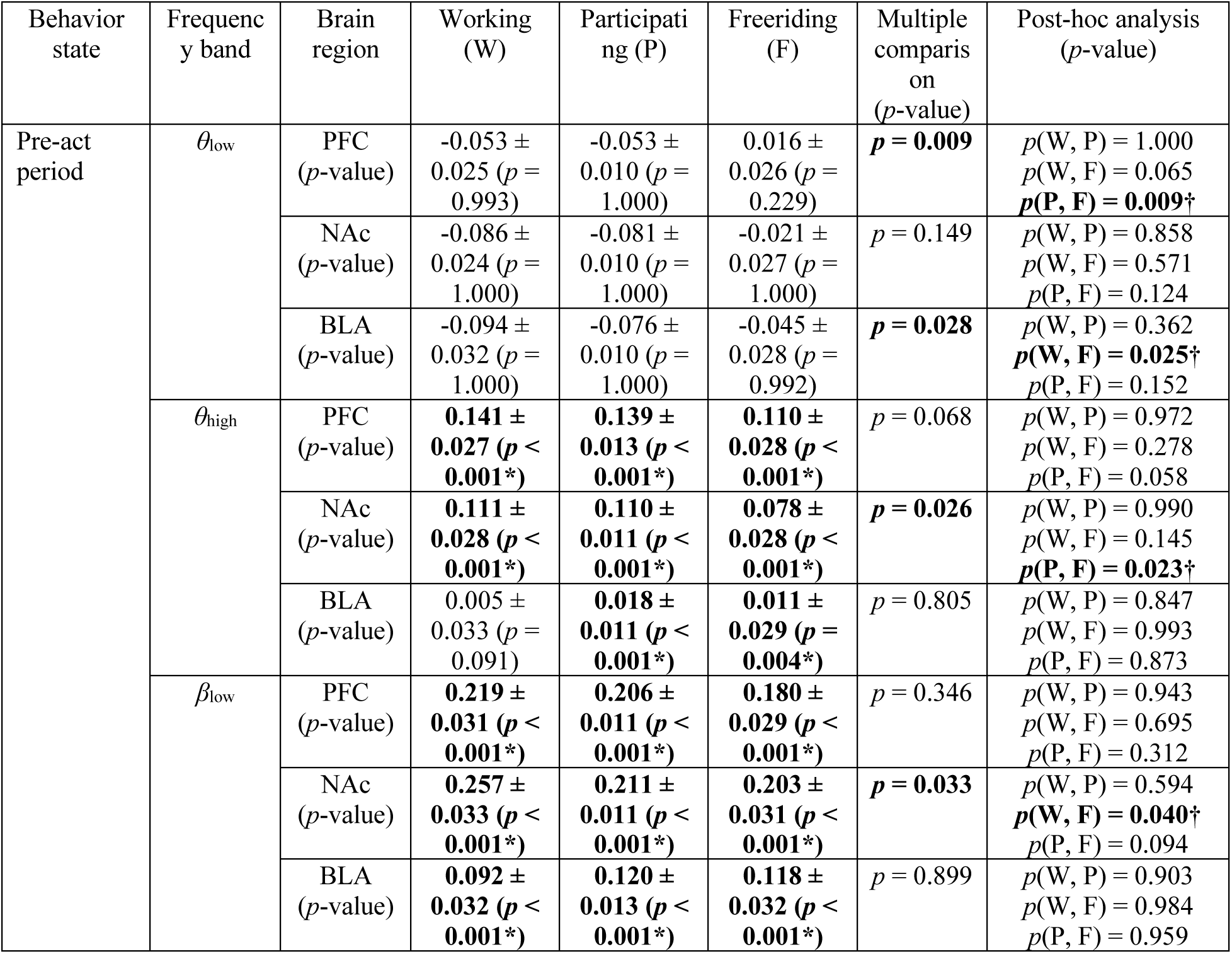
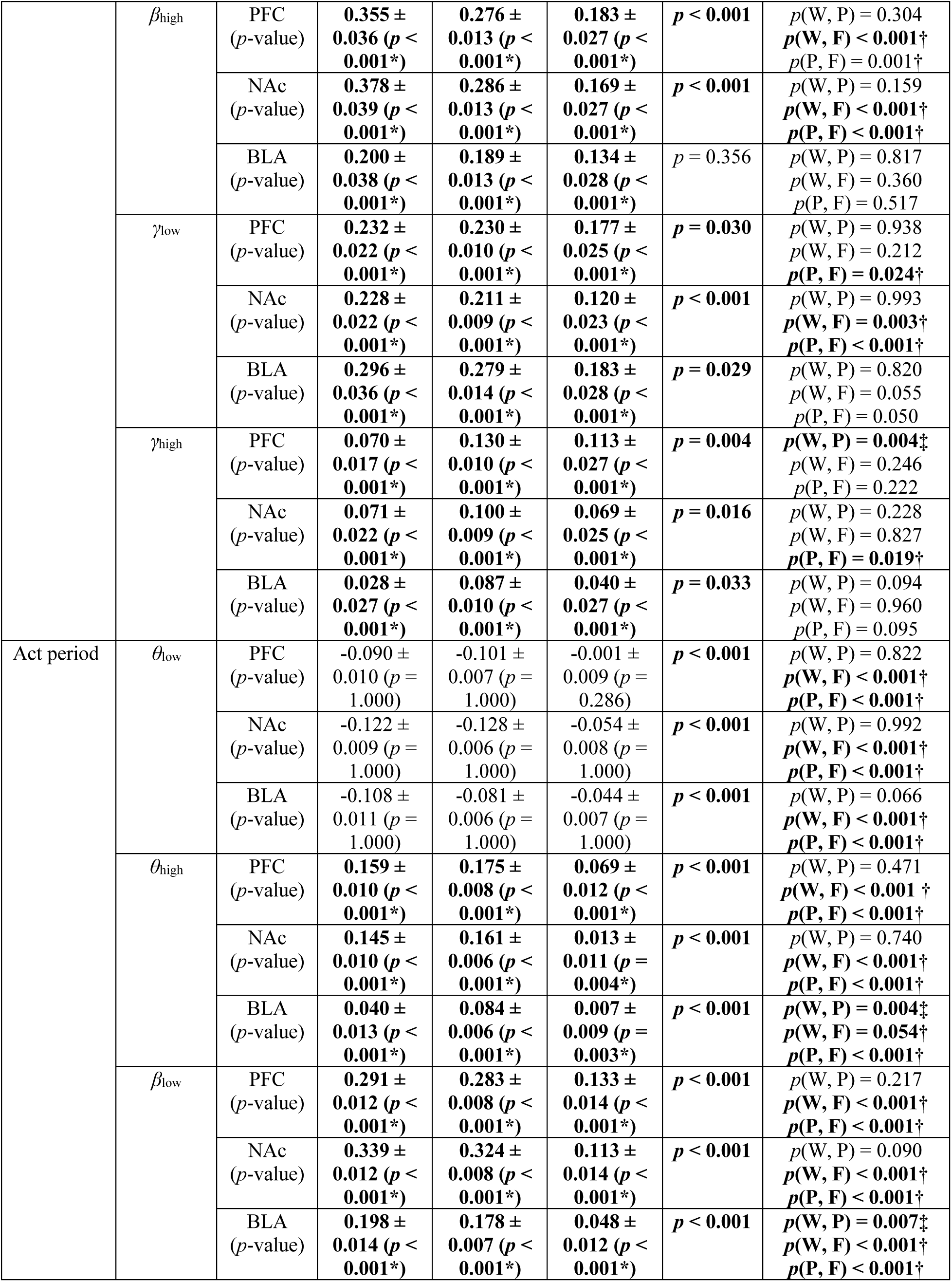
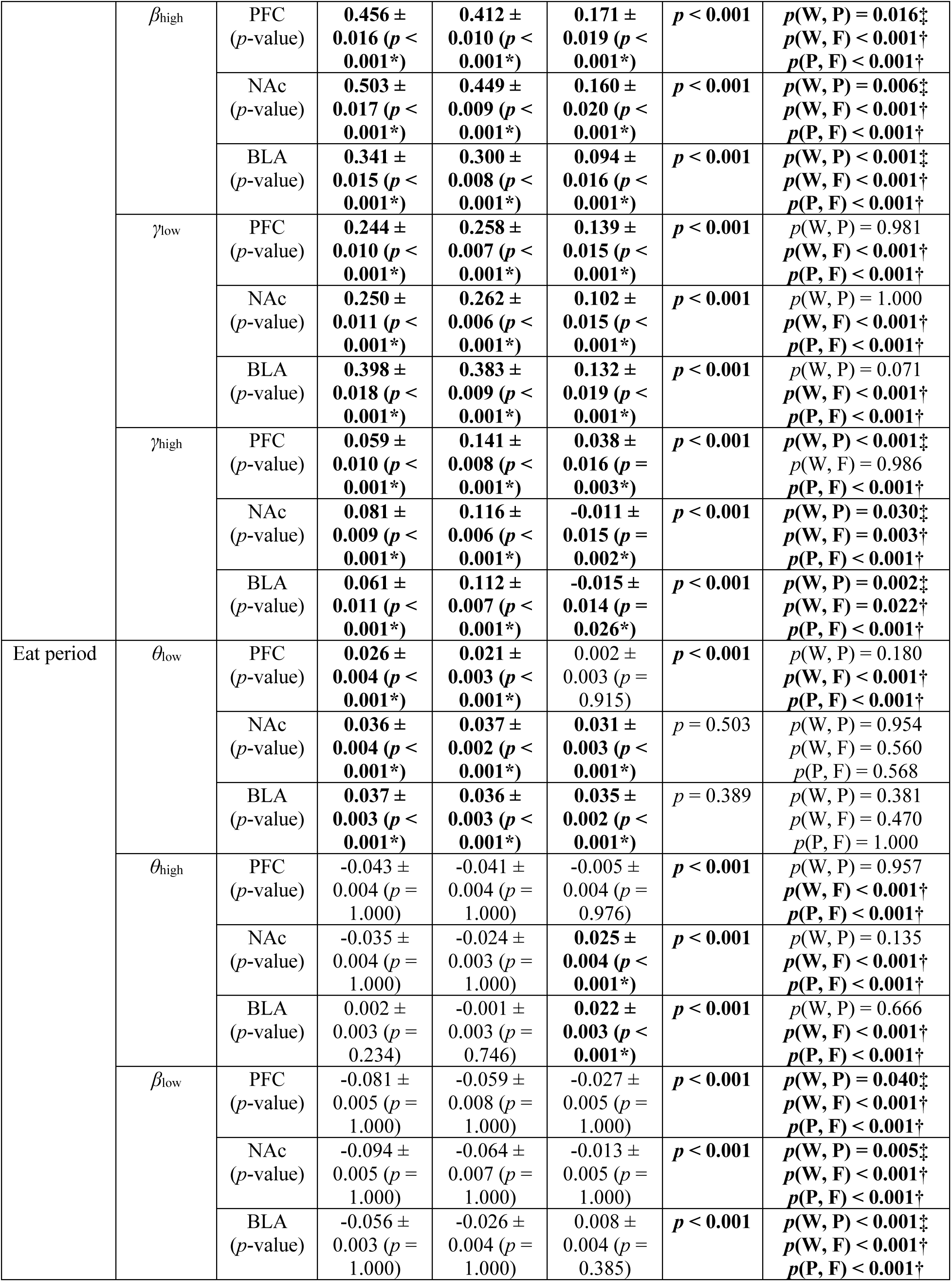
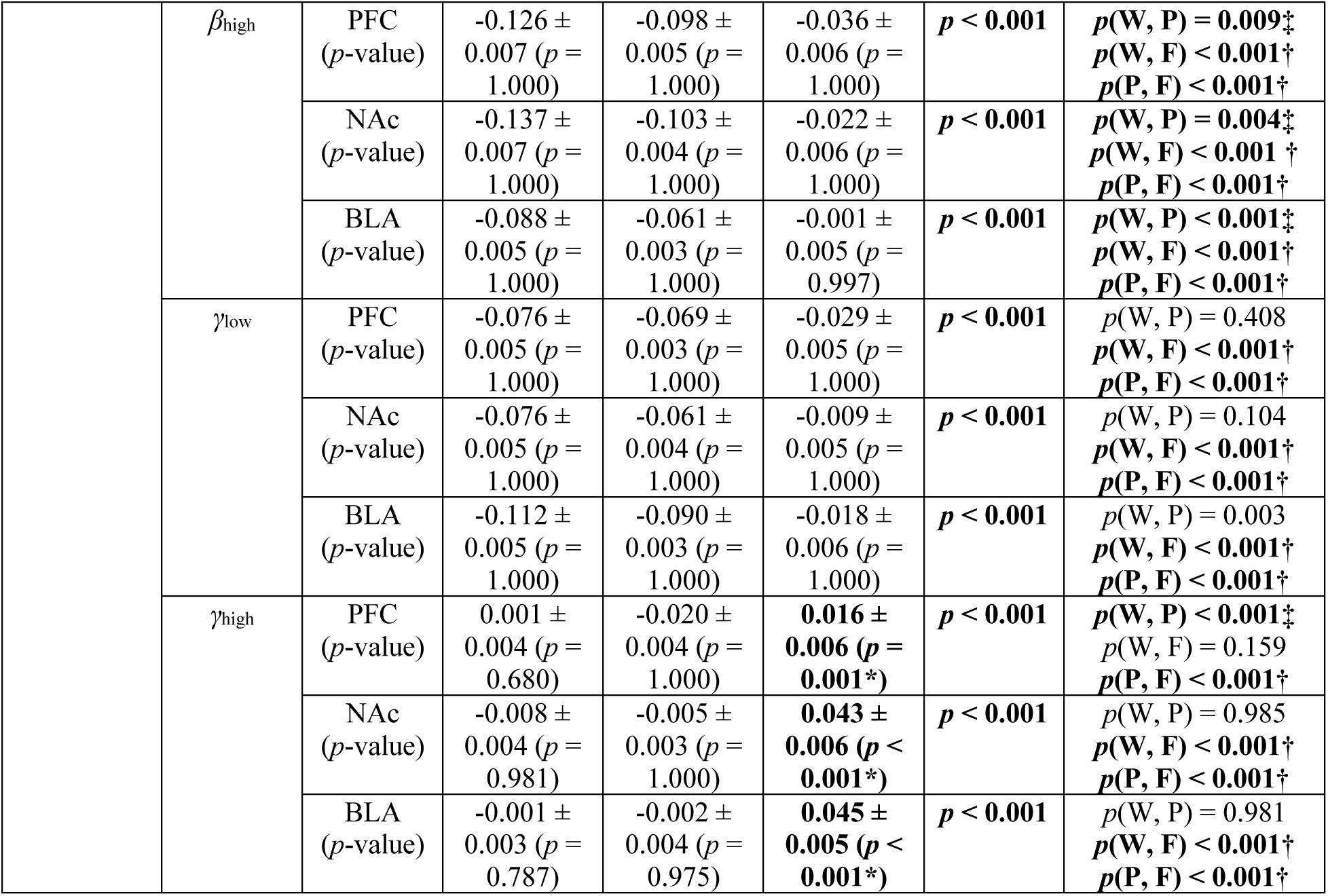
Role-specific z-scored power differences across behavioral states. This table summarizes the z-scored spectral power values across different behaviors (Working, Participating, and Freeriding) during group foraging task (Group A, 162 trials). The results of the null hypothesis test on statistically significant differences from the baseline and indicated by asterisks (*). The results of *post-hoc* pairwise comparisons following the Kruskal-Wallis test are included. Daggers (†) indicate statistically significant differences between freeriding and other roles, while double daggers (‡) indicate significant differences between working and participating. The pairs (W, P), (W, F), and (P, F) denote comparisons between working and participating, working and freeriding, and participating, and freeriding, respectively. *α* = 0.05 for all statistical tests. Frequency bands are defined as follows: *θ*_low_ (4–8 Hz), *θ*_high_ (8–12 Hz), *β*_low_ (18–24 Hz), *β*_high_ (24–32 Hz), *γ*_low_ (35–50 Hz), and *γ*_high_ (70–90 Hz).

**Supplementary Table 7.**
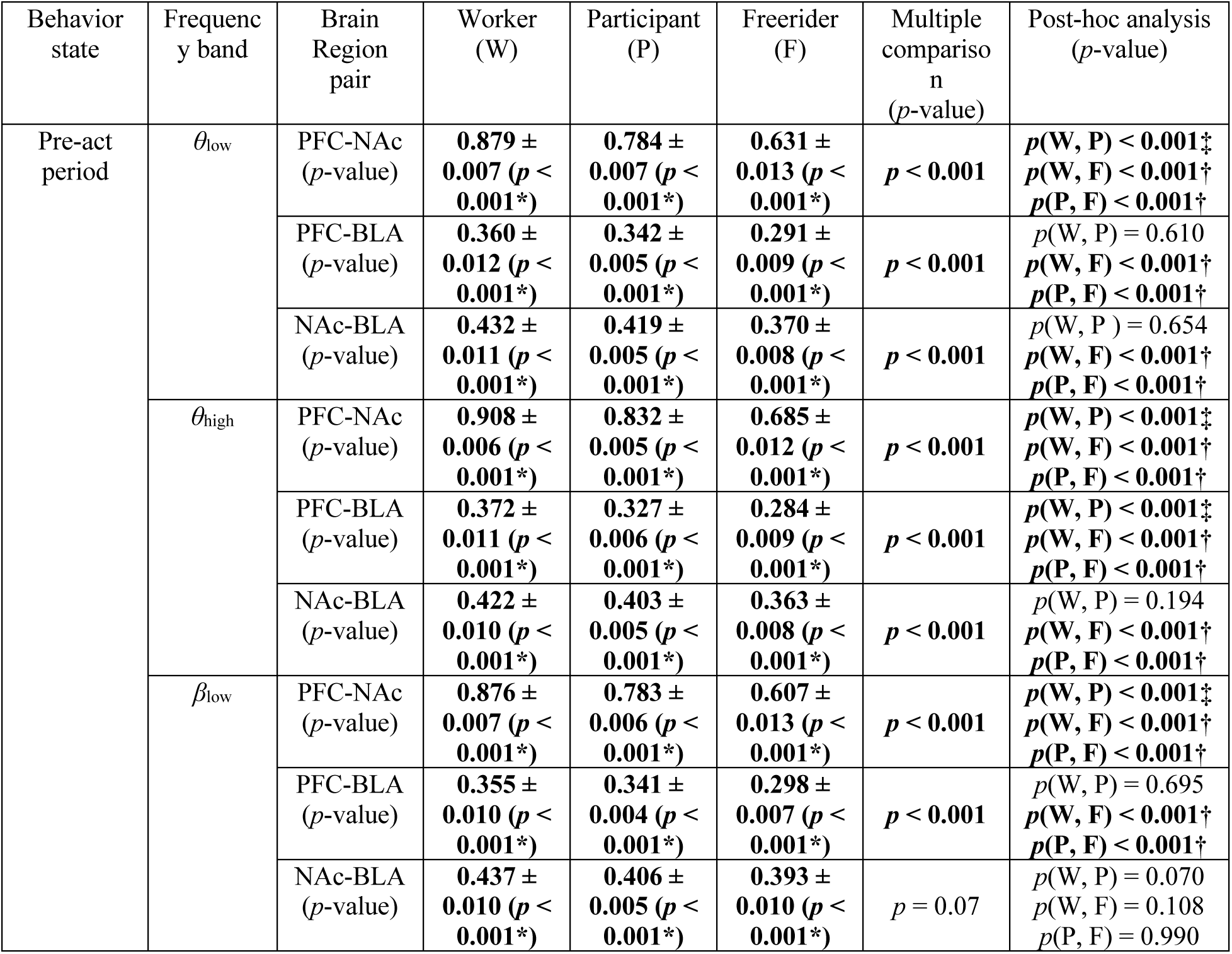
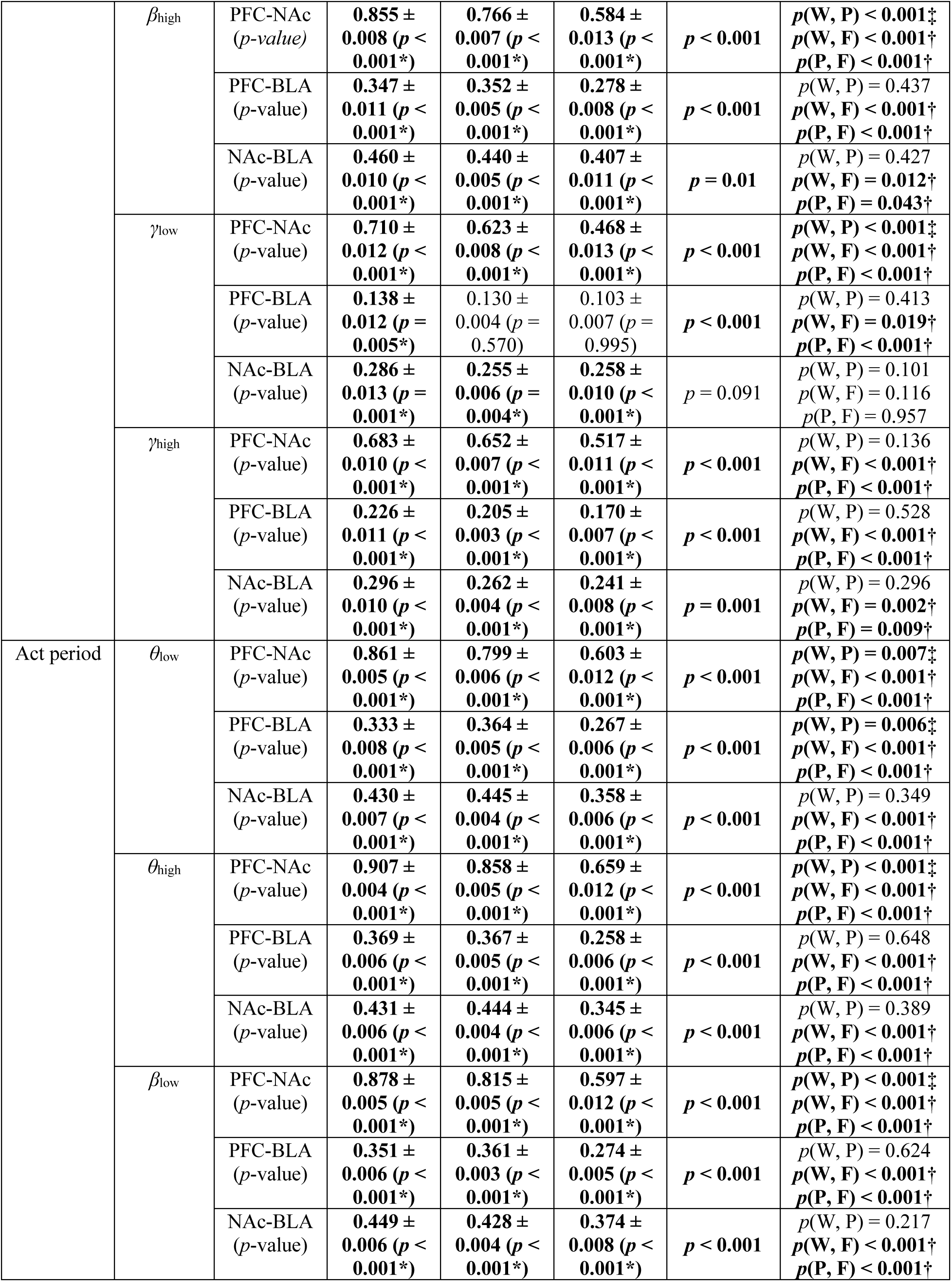
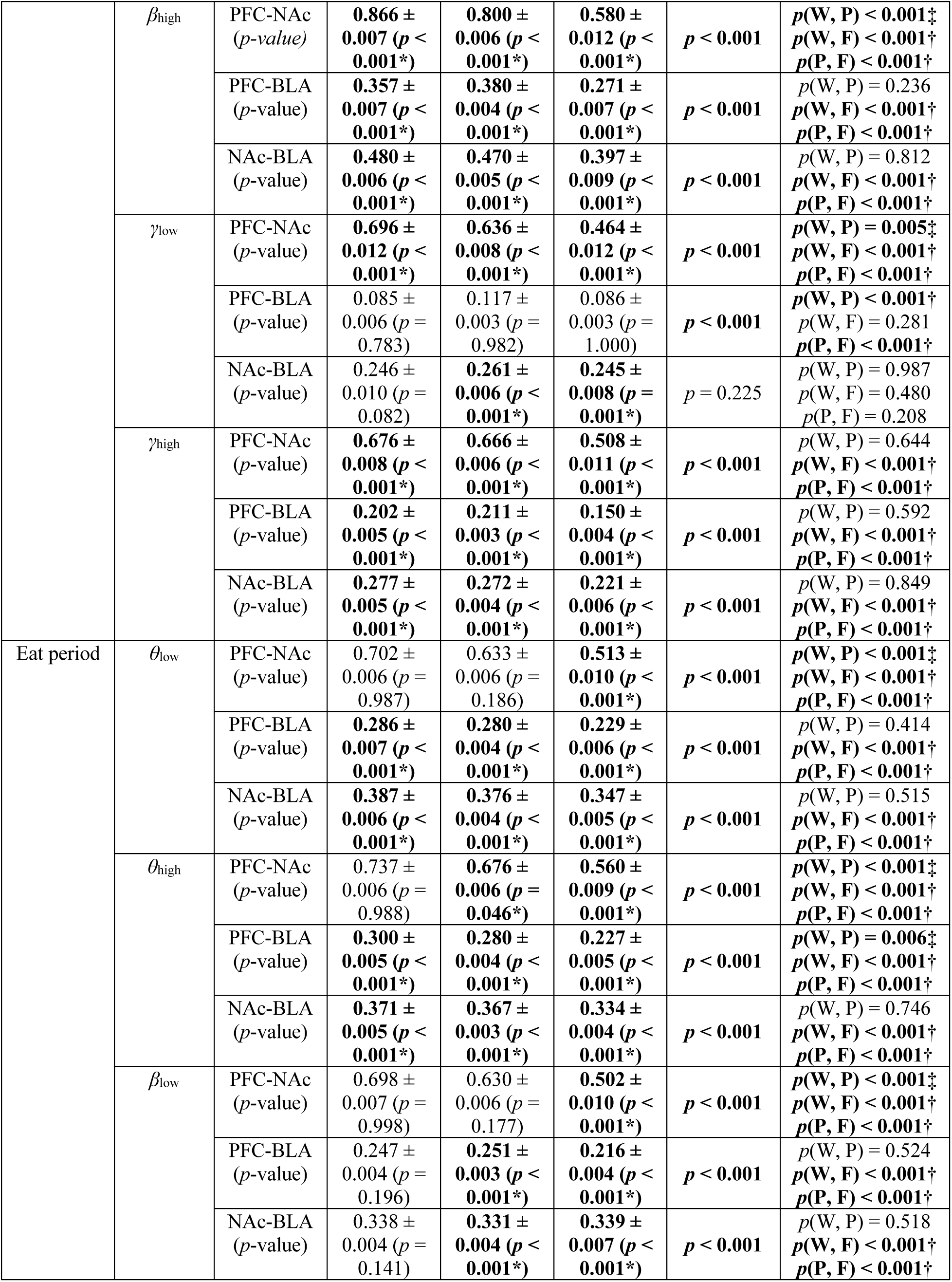
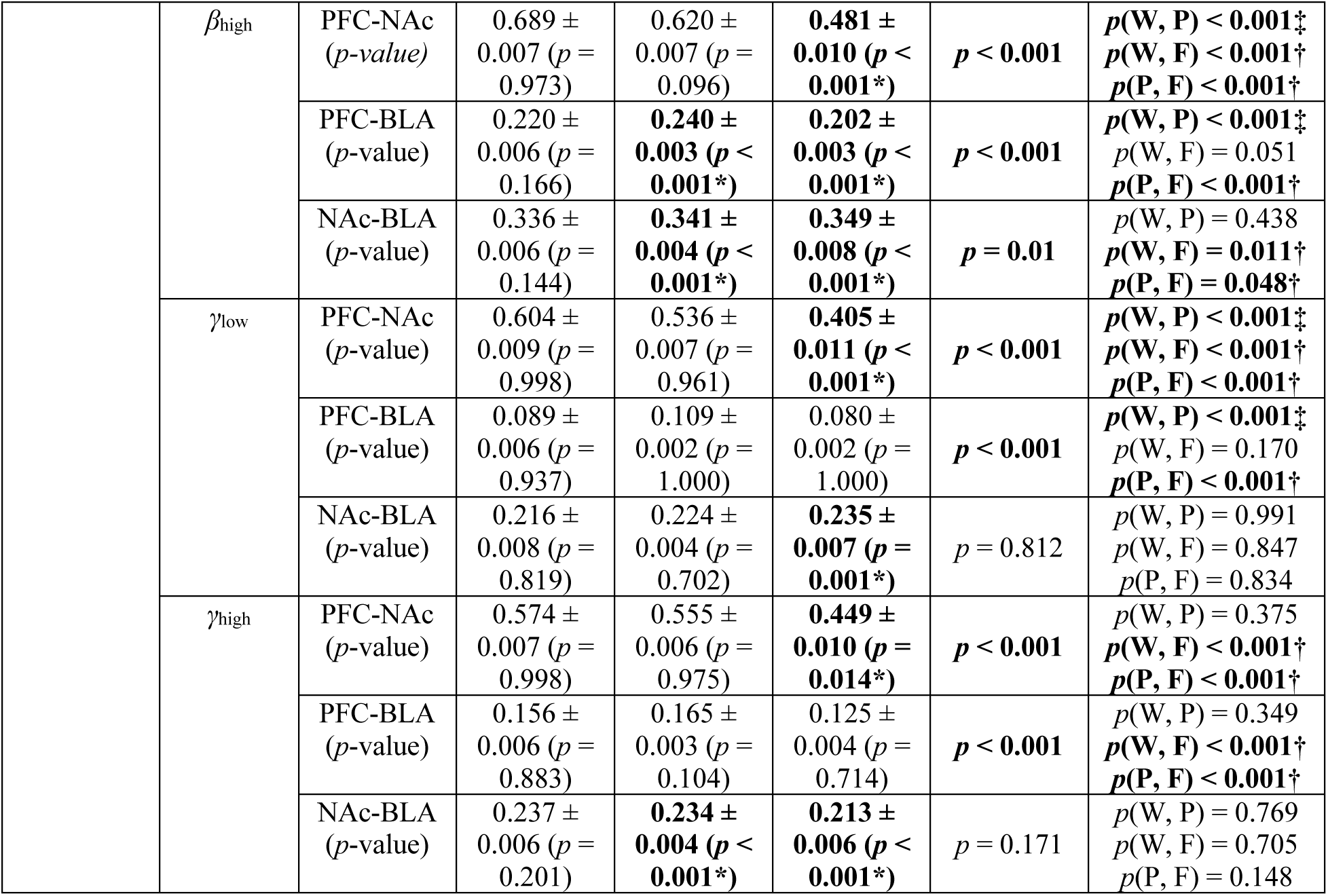
Role-specific network coherence across behavioral states. This table summarizes the spectral coherence between two regions within mPFC-BLA-NAc circuit across different roles (Working, Participating, and Freeriding) during group foraging task (Group A, 162 trials). The results of null hypothesis test on statistically significant differences from the baseline and indicated by asterisks (*). The results of *post-hoc* pairwise comparisons following the Kruskal-Wallis test are included. Daggers (†) indicate statistically significant differences between freeriding and other roles, while double daggers (‡) indicate significant differences between working and participating. The pairs (W, P), (W, F), and (P, F) denote comparisons between working and participating, working and freeriding, and participating, and freeriding, respectively. *α* = 0.05 for all statistical tests. Frequency bands are defined as follows: *θ*_low_ (4–8 Hz), *θ*_high_ (8–12 Hz), *β*_low_ (18–24 Hz), *β*_high_ (24–32 Hz), *γ*_low_ (35–50 Hz), and *γ*_high_ (70–90 Hz).

**Supplementary Table 8.**
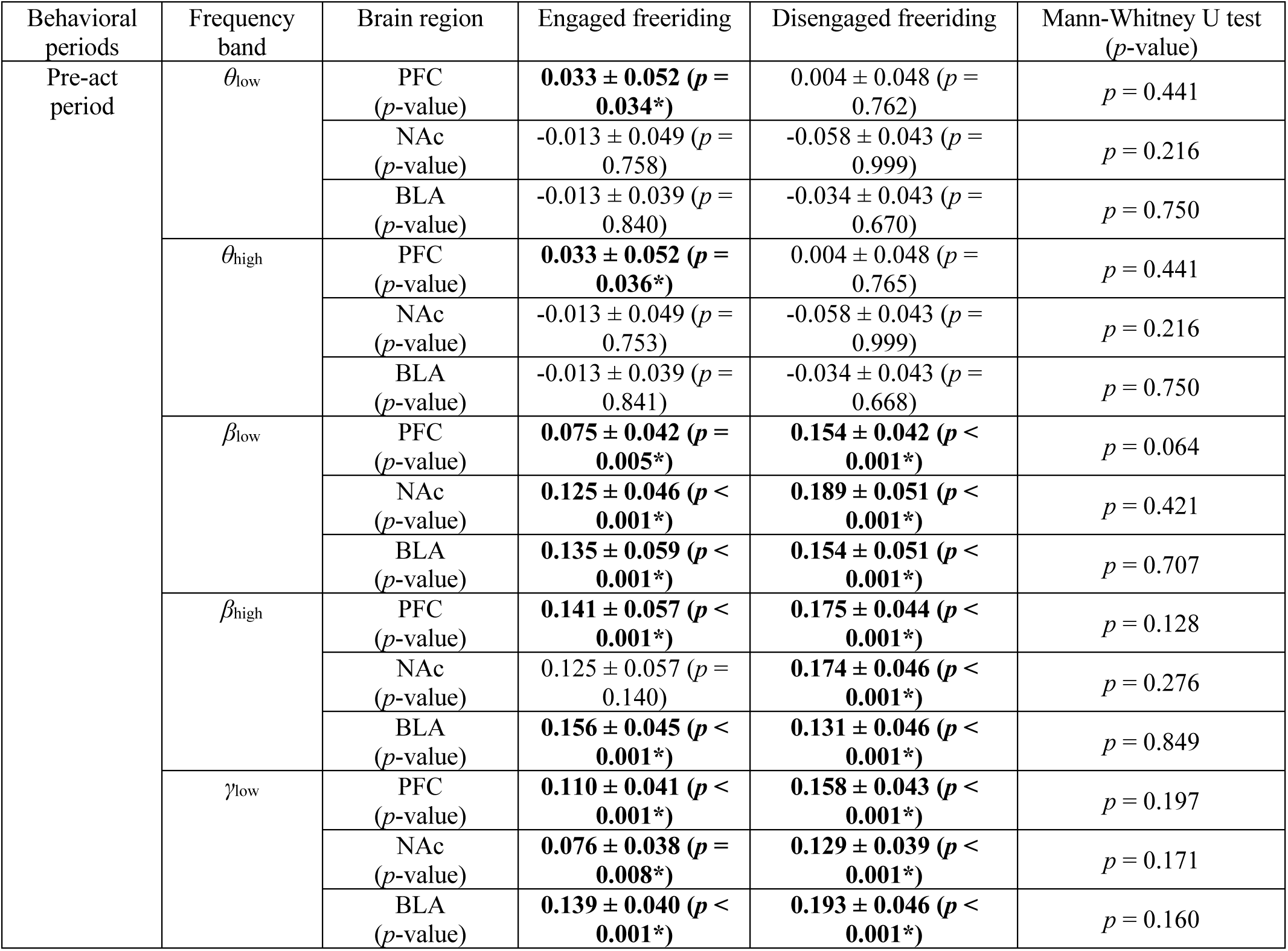
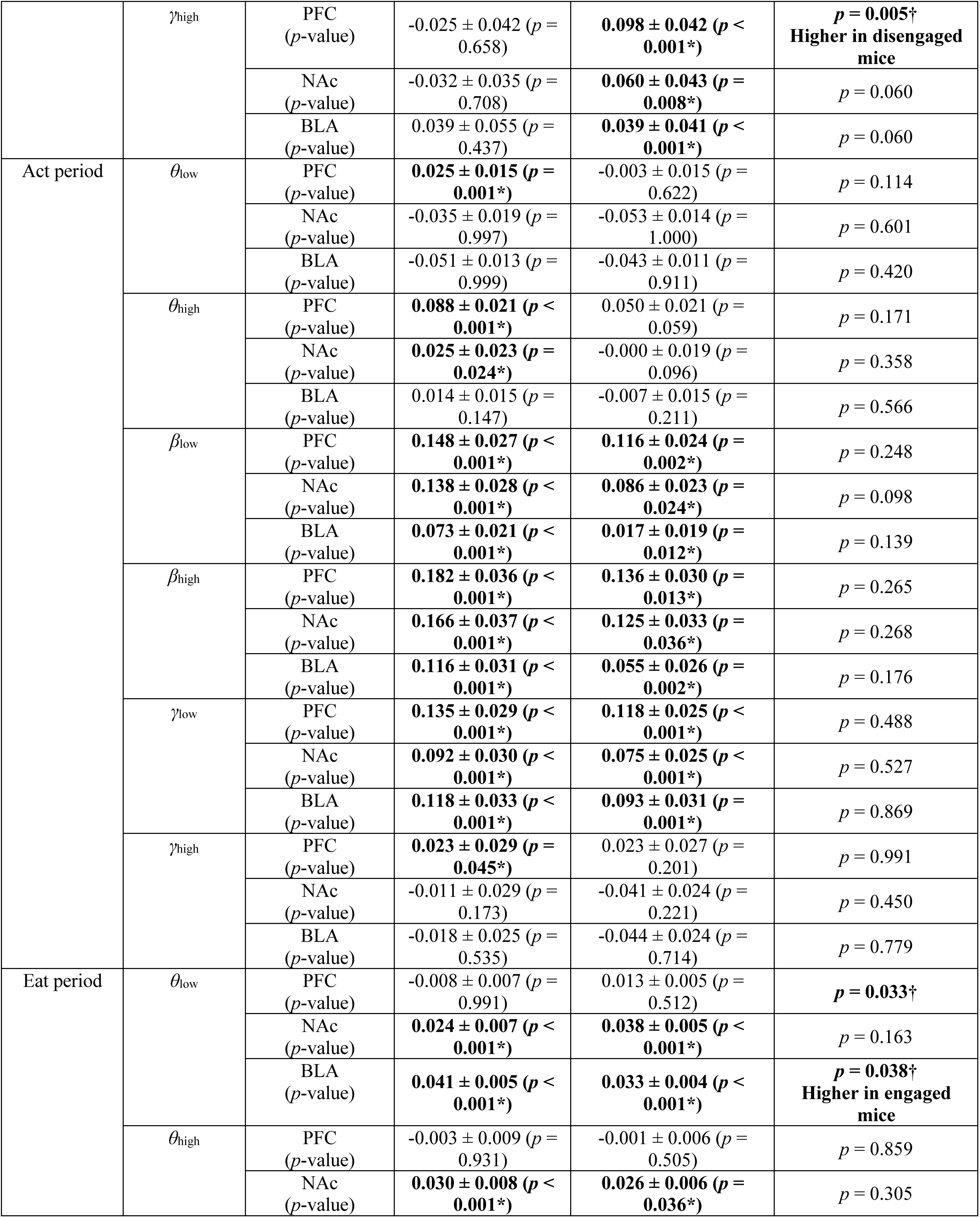
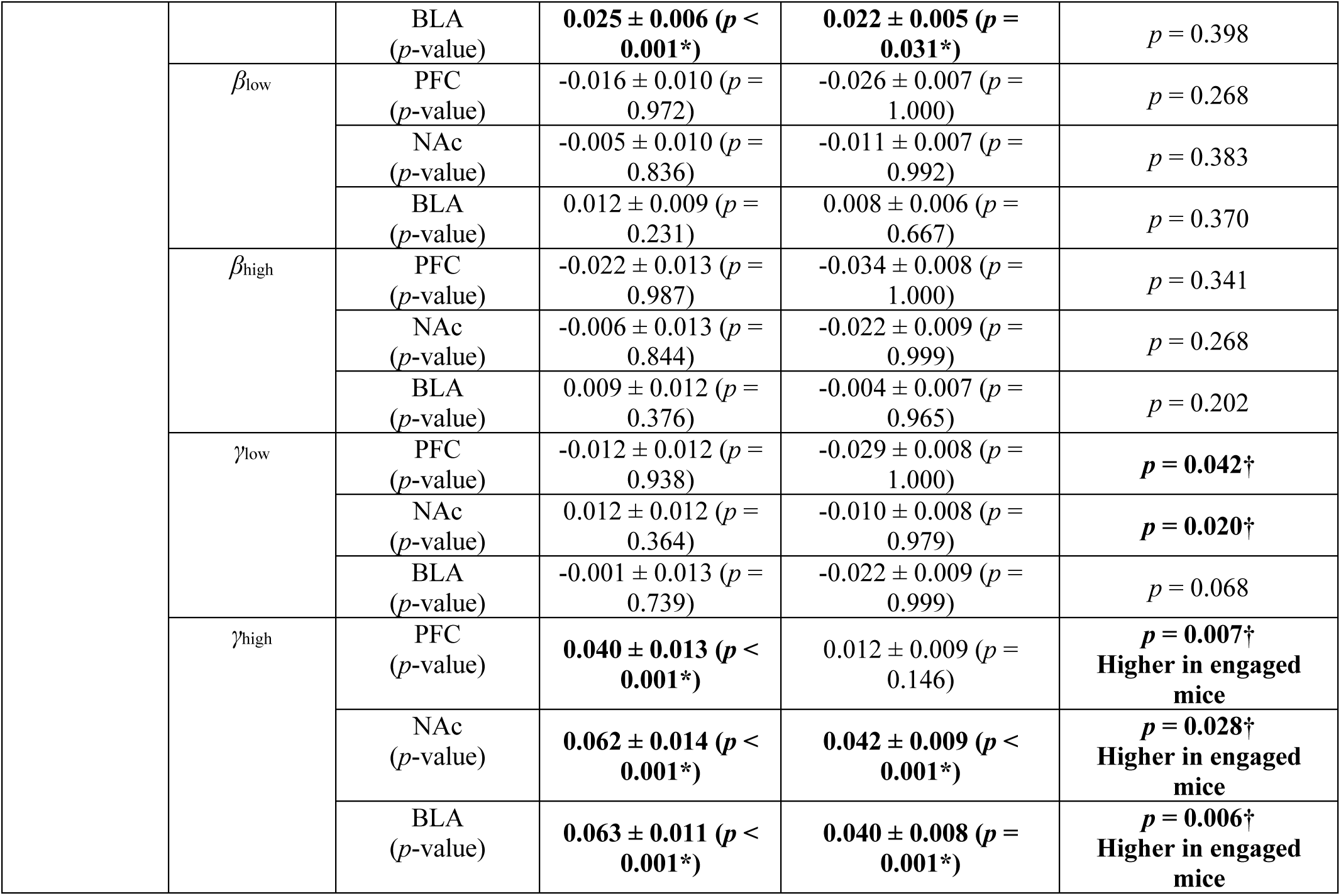
Engaged and disengaged freeriding behavior specific z-scored powers across behavioral states. This table summarizes the z-scored spectral power values in local field potentials (LFPs) of during engaged and disengaged freeriding behaviors (62 trials for engaged freeriding, 107 trials for disengaged freeriding). Engaged freeriding are defined as those who exhibit interest in the task by moving around outside the robot zone while others forage inside the robot zone, whereas disengaged freeriding remain still in the residence zone even after the snack is introduced. Asterisks (*) indicate statistically significant differences from the baseline after testing against the null hypothesis (*α* = 0.05). Daggers (†) denote statistically significant differences between engaged and disengaged freeriding behaviors (Mann-Whitney U test, *α* = 0.05). Frequency bands are defined as follows: *θ*_low_ (4–8 Hz), *θ*_high_ (8–12 Hz), *β*_low_ (18–24 Hz), *β*_high_ (24–32 Hz), *γ*_low_ (35–50 Hz), and *γ*_high_ (70–90 Hz).

**Supplementary Table 9.**
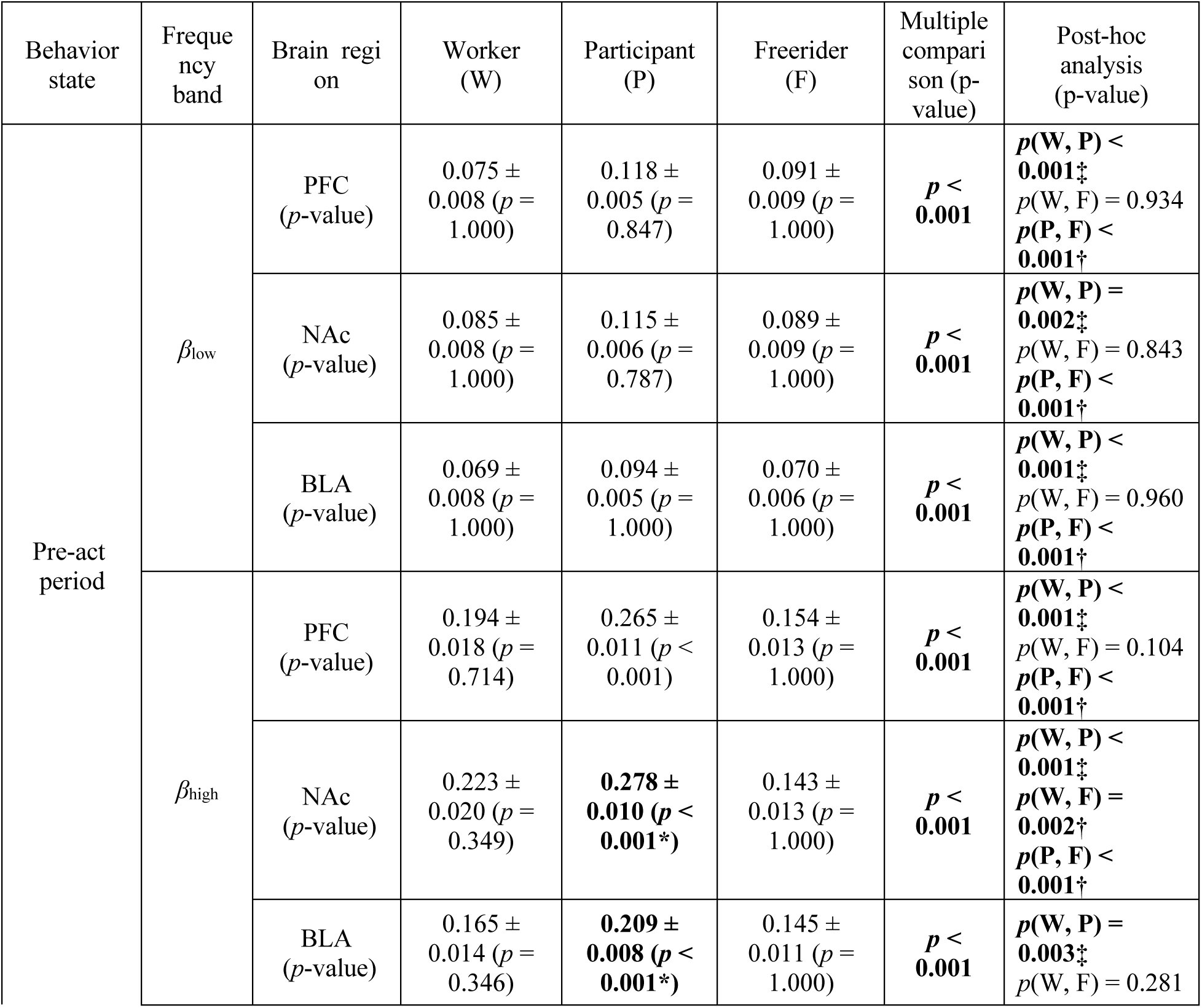
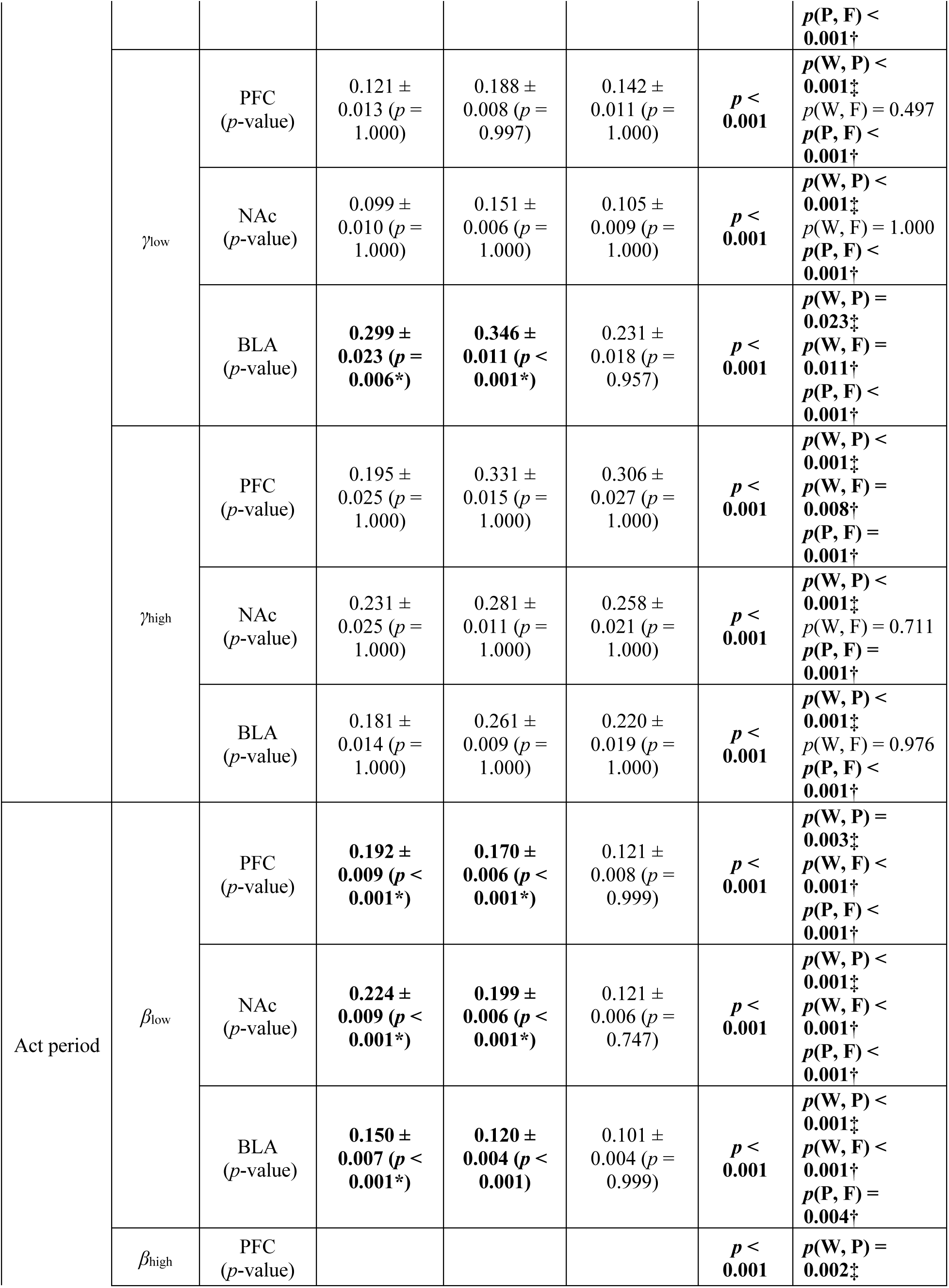
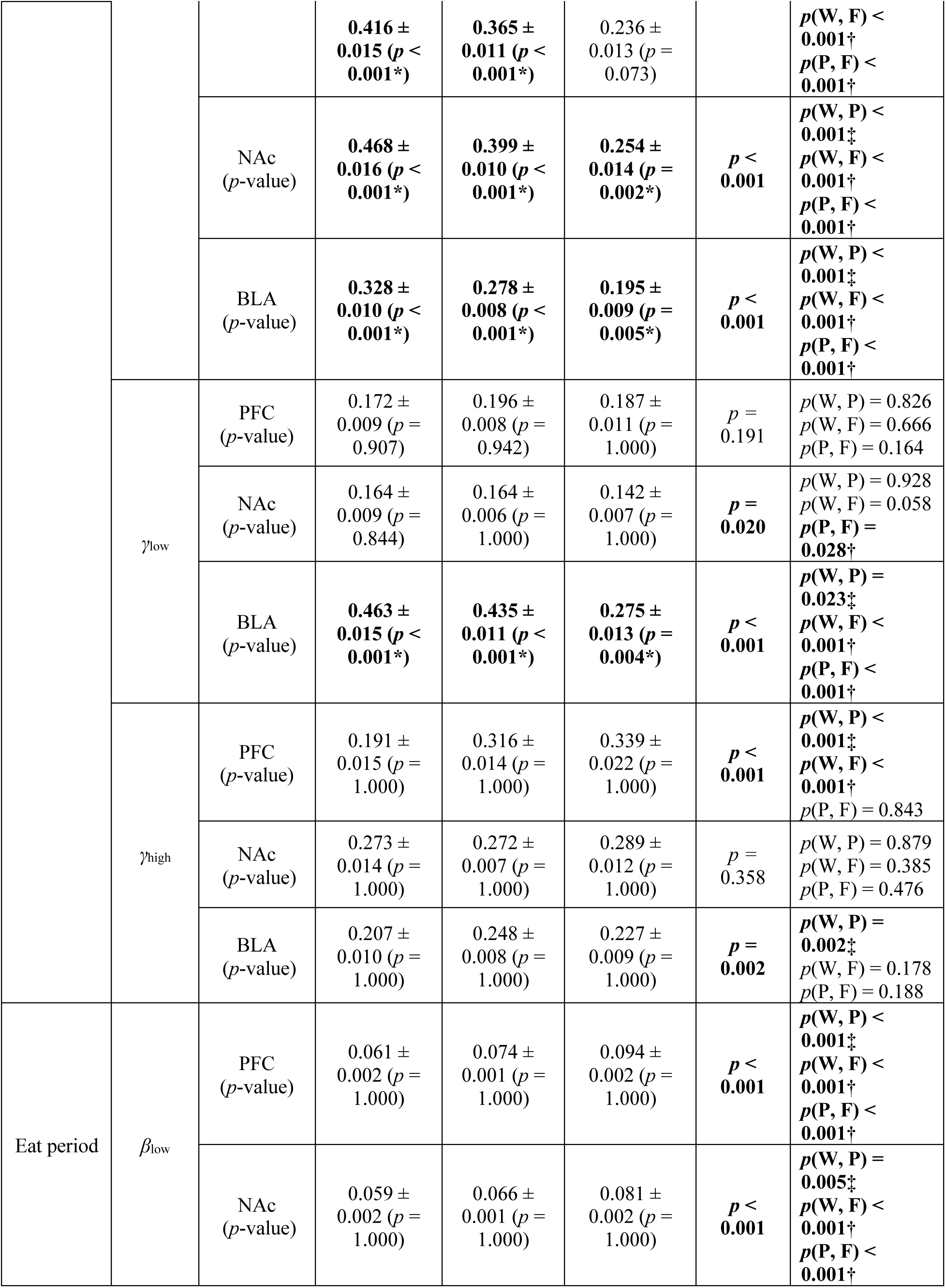
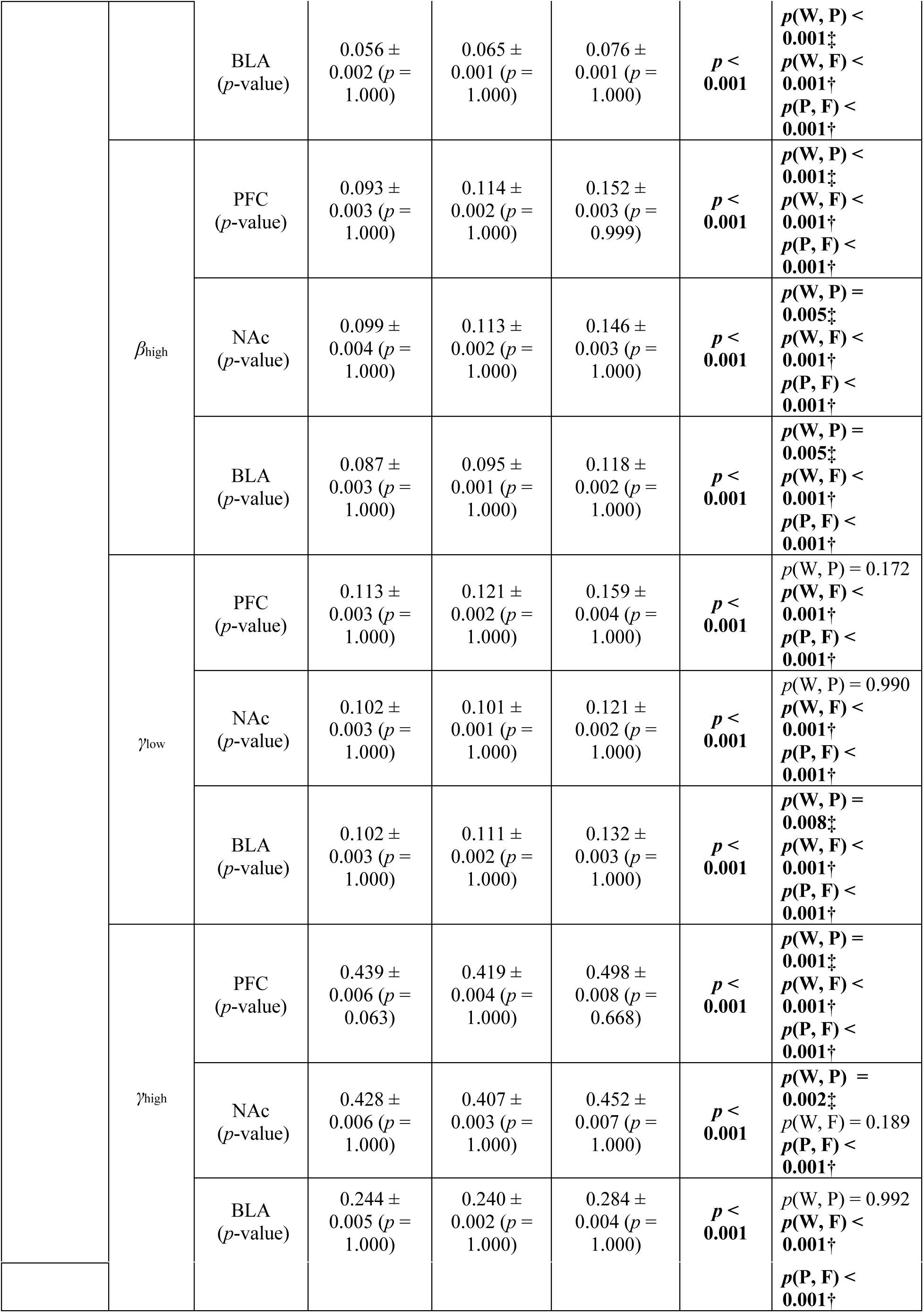
Role-specific burst density differences across behavioral states. This table summarizes the burst density values across different roles (Working, Participating, and Freeriding) during group foraging task (Group A, 162 trials). The results of null hypothesis test on statistically significant differences from the baseline and indicated by asterisks (*). The results of post hoc pairwise comparisons following the Kruskal-Wallis test are included. Daggers (†) indicate statistically significant differences between freeriding and other roles, while double daggers (‡) indicate significant differences between working and participating. The pairs (W, P), (W, F), and (P, F) denote comparisons between working and participating, working and freeriding, and participating, and freeriding, respectively. α = 0.05 for all statistical tests. Frequency bands are defined as follows: θlow (4–8 Hz), θhigh (8–12 Hz), βlow (18–24 Hz), βhigh (24–32 Hz), γlow (35–50 Hz), and γhigh (70–90 Hz).

